# Rooting the deep divergence of land plants

**DOI:** 10.1101/2025.11.29.691271

**Authors:** Hengchi Chen, Jan de Vries

## Abstract

Unravelling the deep phylogenetic structure of embryophytes is essential for understanding their evolutionary success on land. However, pronounced phylogenetic dispute remains: while recent phylogenomic studies have recurrently recovered the bryophyte monophyly in the presence of algal outgroups, absence of such outgroup representatives yields paraphyletic bryophytes. Here, we assembled a large-scale genome dataset encompassing 243 embryophyte species and 12 representative algal species to test the hypothesis of bryophyte monophyly. Across different substitution models and species tree inference methods, we recover strong support for bryophyte monophyly in the presence of algal outgroups; by varying the algal outgroup sampling, we revealed a salient outgroup effect that can significantly alter the embryophyte deep divergence relationship. We show that (1) bryophyte monophyly is consistently supported across three distinct outgroup-free rooting methods; and (2) incomplete lineage sorting, substitution saturation, heterogeneous site and branch profiles through deep time underpin prevalent phylogenomic discordance across genes and sites. Our data explain decades of difficulty in resolving the deep dichotomy of land plants. Bryophyte monophyly is not an artefact by outgroups but underpinned by genuine phylogenetic signal.

## Introduction

The successful colonization of land by plants marked a pivotal evolutionary transition that gave rise to embryophytes (land plants), ushering in complex terrestrial life^1^. A string of key phenotypic and genomic innovations^2^ underpin their remarkable ecological and evolutionary success. To trace the exact evolutionary path that embryophytes took to adapt to and diversify on land, we must identify the root from which all embryophyte descendants emerged. The question of where the embryophyte root truly lies has been studied for decades^3–13^ but remains unsolved.

Early studies in the 1980s placed the root of embryophytes at the branch splitting liverworts from the rest of embryophytes (i.e., hornworts, mosses, tracheophytes) based on morphological characters^14–16^ (Fig. 1; Table S1). Nonetheless, the morphology-based inference appeared labile that a number of subsequent studies^4^ had deduced discrepant root positions based on different sets of morphological characters and/or different phylogenetic methods. For instance, Garbary et al. (1993) inferred the embryophyte root at the divergence of monophyletic bryophytes and tracheophytes based on 90 male gametogenesis-related characters using the maximum parsimony (MP) method^17^. During the 1990s, numerous studies based on a few marker genes emerged^9,18–26^, although still yielding discordant results (Fig. 1; Table S1). For instance, Hedderson et al. (1996) inferred the embryophyte root at the divergence of hornworts and the rest of embryophytes applying both the MP and maximum likelihood (ML) methods on 18s rRNA gene sequences^22^. Whereas Lewis et al. (1997) placed liverworts as the root applying both MP and ML methods on *rbcL* gene^23^. More alternative rooting scenarios were proposed as more marker genes became available and more sophisticated methods emerged, e.g., placing mosses as the root^24,27^, or placing setaphytes (the clade consisting of all mosses and liverworts) as the root^28–30^. Nevertheless, the most recent large-scale nuclear genome-based phylogenomic studies all recovered the monophyletic bryophytes as the best root position^31–37^ (Fig. 1; Table S1). Moreover, many recent plant systems biology studies pertinent to embryophytes either assume or directly apply the bryophyte monophyly in their analyses or interpretations^2,38–48^, entrenching it as the prevailing paradigm in current practice.

**Figure 1.**
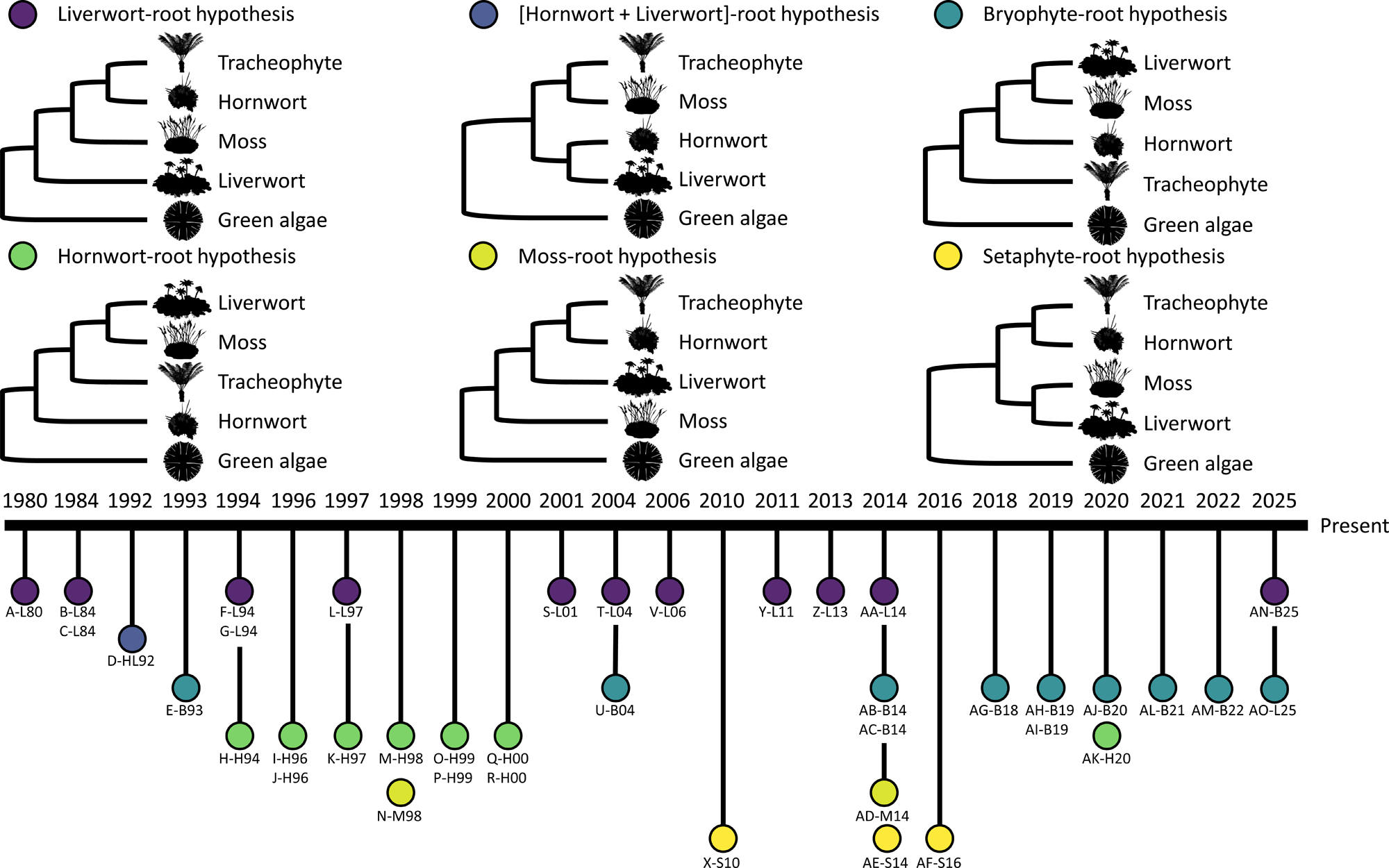
Competing hypotheses regarding the root position of embryophytes. Phylogenetic studies resolving the deep divergence relationship of embryophytes were inspected from 1980 until present and we pinpointed 6 competing hypotheses that were derived by modern phylogenetic methods and clearly preferred by the original studies when alternative scenarios were present. The publication year of each considered study is denoted on the bottom time scale, whose length is not to scale. Each competing hypothesis is marked by a distinct color. Each considered study is represented by a unique code, for instance, ‘A-L80’ indicates the study published in 1980 that supported the liverwort-root hypothesis with a unique index as ‘A’, see Table S1 for detailed information. Silhouette images are from Phylopic (license: CC0 1.0 Universal Public Domain Dedication), in which the hornwort image represents *Anthoceros agrestis*, the liverwort image represents *Marchantia polymorpha*, the moss image represents *Sphagnum* sp., the tracheophyte image represents *Cycas circinalis*, and the green algae image represents *Micrasterias papillifera*.

However, when inspecting the key evidence supporting the bryophyte monophyly from all those phylogenetic and phylogenomic studies, we find that the gene families used in their phylogenetic analyses always include algal outgroups. In particular, Harris et al. (2022)^34^ did conduct outgroup-free rooting analyses utilizing ALE^49^, STRIDE^50^, and the AU test^51^ based on 249 single-copy gene families (SOG) derived from only 24 embryophyte genomes: both the bryophyte monophyly and the bryophyte paraphyly scenarios stood all the tests. In fact, the bryophyte monophyly was never the best rooting scenario in any of those tests. Only by adding algal outgroups into the gene families and reconducting the topology test did yield results that rejected all rooting scenarios except for the monophyletic bryophyte scenario, as described in many previous studies^31–33^. What is the evidence for bryophyte monophyly? Is it an artefact from algal outgroups?

Outgroup rooting is the prevalent method nowadays to root a gene tree or species tree^52–55^. Nonetheless, this method is not without pitfalls. The accuracy of outgroup rooting is strongly dependent on the choice of outgroup representatives^52,56–59^. If inappropriately chosen, they can interfere with phylogenetic relationships among the ingroup^54,59–62^. As such, fast-evolving or compositionally shifted lineages could be misplaced towards outgroups^63^, manifested as early diverging members of the ingroup in a paraphyletic manner due to factors such as long branch attraction (LBA). Moreover, the outgroup-rooting method represents only one of the methods available for rooting a phylogenetic tree^54^. Outgroup-free rooting methods include 1) midpoint rooting^64^, 2) minimal ancestor deviation (MAD) rooting^65^, 3) minimum variance rooting^66^, 4) molecular clock-based rooting^67^, 5) non-reversible substitution model-based rooting^68^, 6) paralogue-based rooting^69^, 7) multispecies coalescent model-based rooting^70^, 8) gene family evolution model-based rooting^71^, such as the rooting analyses presented in Harris et al. (2022)^34^ using ALE^49^ and STRIDE^50^, 9) “Hennigian” methods relying on rare molecular events^63^ (e.g., translocation^72^, insertion and deletion^73–76^, shift in amino acid usage^77^, intron^78^, and protein domain^79^), and 10) fossil-based rooting^54,80^. To the best of our knowledge, no study has systematically dissected the effect of outgroups on the deep divergence of embryophytes, and the current majority of phylogenetic inferences by outgroup rooting remains to be corroborated by outgroup-free rooting methods.

Here we set out to resolve the discordance in the deep genetic structure of the land plant phylogeny. We assembled a large-scale embryophyte genome dataset encompassing 154 bryophyte species and 89 tracheophyte species with 12 algal species as outgroup representatives to investigate the rooting question of embryophytes, leveraging the outgroup-rooting method and 3 different outgroup-free rooting methods, i.e., non-reversible substitution model-based rooting, MAD rooting, and paralogue-based rooting. We show that the monophyletic bryophyte rooting scenario has exclusively obtained consistent support from both the outgroup-rooting and outgroup-free rooting methods. Nonetheless, we detected widespread phylogenomic discordance across gene families and amino acid sites with or without algal outgroups, derived from ancestral incomplete lineage sorting (ILS), substitution saturation, branch-wise and site-wise heterogeneity. We demonstrate that failing to account for these confounding factors will lead to discordant deep divergence scenarios of embryophytes, which provides a plausible explanation for the prevalent discrepancy emerged from previous embryophyte phylogenetic studies relying on several to a few marker genes without meticulous curation of informative sites or model comparison. Altogether, this study constitutes the first evidence showing that the bryophyte monophyly is not an artefact by outgroups but underpinned by genuine phylogenetic signals.

## Results

### Construction of single-copy gene family datasets

We clustered protein-coding genes into gene families (i.e., orthogroups via OrthoFinder^81^) using a genome datasets of 243 embryophyte species and of 243 embryophyte and 12 algal species, referred to as the 243-genome dataset and the 255-genome dataset, respectively. Since there was no single-copy gene family (SOG) deduced, we constructed the refined SOGs (rSOGs) based on multi-copy orthogroups that have species coverage of 100% (see Methods), yielding 273 and 137 rSOGs for the 243-genome dataset and the 255-genome dataset, respectively, henceforth the 273-rSOG and 137-rSOG datasets. Besides the de novo inference of SOGs, we also constructed SOGs upon our 243-genome dataset using the reference SOGs curated in the embryophyta_odb12 dataset from OrthoDB^82^, leveraging the reference hidden Markov model (HMM) profiles (see Methods). This led to 2,026 SOGs, henceforth the 2026-RefSOG dataset. Unlike the rSOG dataset, the RefSOG dataset can have species coverage less than 100% due to factors such as lineage-specific loss; >50% of the RefSOGs have at least 224 species present and >75% of the RefSOGs have at least 208 species present (Fig. S1), indicating an overall good species representation.

#### Outgroup rooting

##### Bryophyte monophyly recovered when rooted with 12 algal outgroup representatives

To decipher the embryophyte deep divergence scenario with the outgroup-rooting method, we conducted the species tree inference using the (i) concatenation and (ii) multispecies coalescent (MSC) methods on the 137-rSOG dataset (including 12 algal outgroup representatives; see Methods). Both approaches support the embryophyte monophyly and bryophyte monophyly (see below). The local posterior probability (LPP) calculated in the MSC method is 1.00 and 0.91 for the crown nodes of embryophytes and bryophytes (Fig. 2), respectively. The ultrafast bootstrapping (UFBoot), SH-aLRT test, approximate Bayes test, and the local bootstrapping in the concatenation method all provide full support for the embryophyte monophyly and bryophyte monophyly (see Methods; Fig. S2). The same deep divergence topology is recovered when applying the optimal substitution model evaluated by ModelFinder^83^ in terms of BIC (Fig. S3 and S4).

**Figure 2.**
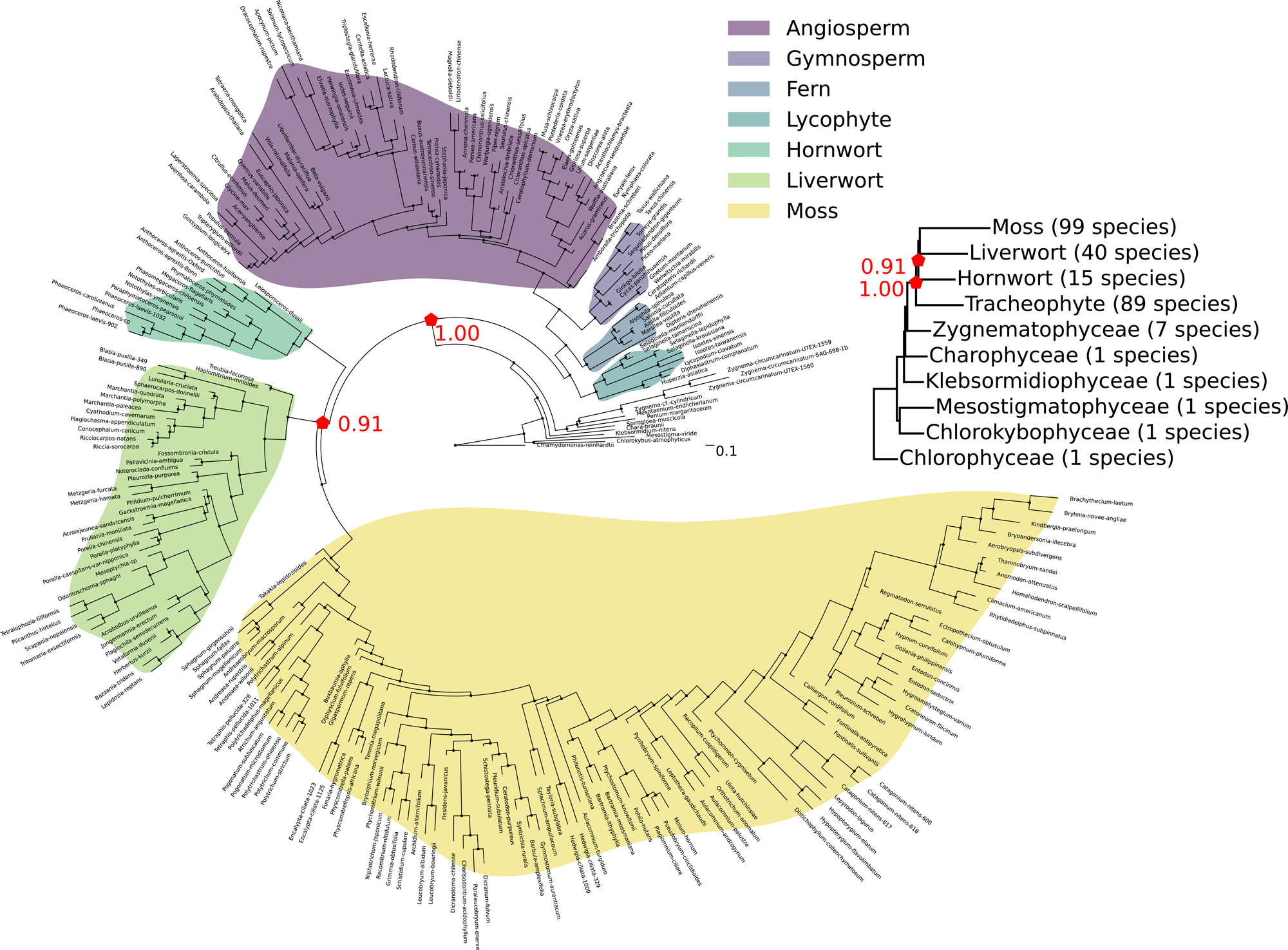
Species tree inferred by the MSC method based on the 137-rSOG dataset. Major embryophyte clades are highlighted in distinct colors, including angiosperms, gymnosperms, ferns, lycophytes, hornworts, liverworts, and mosses. The supporting level for the crown node of bryophytes and embryophytes is measured by the local posterior probability. The branch length represents the number of coalescent unit. A summarizing streptophyte backbone phylogeny is exhibited. The number of sampled species in different clades are denoted within parentheses and clades with more than 1 sampled species have been collapsed into a single representative branch whose length is represented by the median root-to-tip distance among the affiliated sampled species in the summarizing streptophyte backbone phylogeny.

##### Incomplete lineage sorting (ILS) underpins the discordance of embryophyte deep divergence

Previous marker-based phylogenies often resulted in discrepant deep divergence scenarios^6,11,12,84,85^, suggesting the presence of phylogenetic discordance across gene families. The biological sources underpinning such discordance remain elusive. Here, we quantified the level of discordance across gene families regarding the deep divergence of embryophytes and dissected possible biological processes responsible for the observed discordance with a series of triplet frequency analyses (see Methods). Firstly, we calculated the frequency of the two minor discordant triplets regarding the crown group of embryophytes (i.e., q2 and q3; Extended Data Fig. 1a-b), which are 0.286 and 0.283, respectively, indicating a pronounced level of discordance across gene trees. The chi-square goodness-of-fit test shows that the frequency of q2 and q3 is not significantly different (*P* > 0.05; Extended Data Fig. 1b). To mitigate random sampling error^86^, we repeated the triplet frequency analyses on 200 replicate datasets of gene trees (see Methods). The chi-square test of independence shows that the distribution of q2 and q3 is not significantly different (*P* > 0.05; Extended Data Fig. 1c). Such symmetry in the minor triplet frequencies likely excludes gene flow as the major force driving the discordance^87,88^ across gene trees. We further show that the observed discordant triplet frequencies (i.e., q2 + q3) are within the 1σ confidence interval of the null distribution of simulated gene trees under the MSC model (see Methods; Extended Data Fig. 1d), suggesting the consistency within the expectation of a moderate ILS-only scenario. Thus, we propose that a moderate level of ILS emerging in ancestral embryophytes may underpin the observed discordance across gene families in our dataset.

##### An outgroup effect revealed by the stepwise removal of outgroup representatives

The choice of outgroups is pivotal for the accuracy of outgroup rooting. To examine the outgroup effect on the inference of embryophyte deep divergence, we adopt a stepwise removal strategy in which we remove one outgroup representative at a time in the descending order of phylogenetic distance (i.e., from the most distant to the closest) and repeat the species tree inference analysis (see Methods). For the deep divergence inferred by the MSC method, the bryophyte monophyly has been consistently recovered, albeit with varying LPP ranging from 0.49 to 1 (Extended Data Fig. 2b). For the inference of deep divergence by the concatenation method, however, the results vary between bryophyte monophyly and bryophyte paraphyly (Extended Data Fig. 2a). The concatenation method assumes that every SOG has the same evolutionary history and the discordance across gene families is derived only from the gene tree estimation error, which the concatenated MSA can overcome by amplifying the true phylogenetic signals. The presence of prominent discordance derived from ILS as we have shown earlier, however, makes this assumption violated. While this appears to constitute a plausible explanation for the varied results, the concatenation method still has recovered the bryophyte monophyly under certain outgroup settings; other factors than ILS must thus shape the phylogenetic signal of the embryophyte deep divergence during the stepwise removal process of outgroup representatives, which we collectively refer to as the outgroup effect and investigated further.

##### Declined number of decisive sites, increased saturation level, and declined clock-likeness may underpin the outgroup effect

To scrutinize potential factors underpinning the outgroup effect, we analyzed the dynamics of 1) the informative sites, 2) the substitution saturation level, and 3) the tree clock-likeness following the stepwise removal of outgroup representatives (see Methods). We calculated the number of informative sites decisive for the crown node of monophyletic bryophytes, which turned out to monotonically decline (Fig. 3a). The associated site concordance factor (sCF) also monotonically declined from 37.66% to 32.25% (Fig. 3a), which was ultimately surpassed by the site discordance factors for alternative quartets (i.e., sDF1 or sDF2; see Methods). Although the declining number of decisive sites may have weakened the phylogenetic signal supporting bryophyte monophyly, the sCF of up to 37.66% and sDF of up to 38.87% indicate no overwhelming preference for any alternative topology. There rather is substantial conflicting signals across sites. Notably, when the sCF dropped below the null expectation of one-third as the number of decisive sites declined (e.g., upon the removal of the outgroup representative from Charophyceae), the branch support for the bryophyte monophyly also declined from full support to undetermined (Extended Data Fig. 2a). This underscores the importance of sufficient sampling of decisive sites so that the deep divergence signal is less likely to be overwhelmed by noise by chance. The level of substitution saturation is significantly different across datasets that recovered different deep divergence scenarios (*P* = 0.025; Kruskal Wallis test) and datasets recovering the bryophyte monophyly have the lowest level of substitution saturation (Fig. 3b). We calculated the coefficient of variance of root-to-tip distance as a proxy for tree clock-likeness^65^ for each inferred gene tree and found that trees recovering different scenarios exhibit significantly different clock-likeness (*P* = 0.014; Kruskal-Wallis test; Fig. 3c). Bryophyte monophyly is predominantly recovered in the trees with the highest clock-likeness.

**Figure 3.**
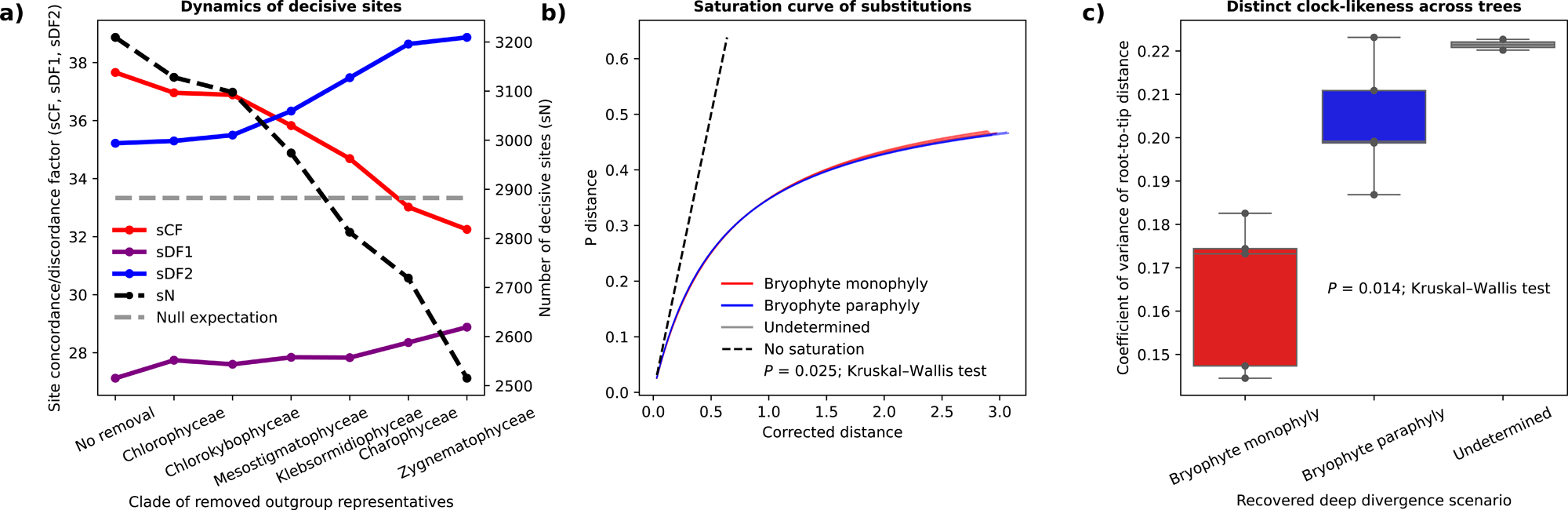
Factors that may affect the concatenation-based phylogenetic inference. (a) The dynamics of the number of informative sites that are decisive for the crown node of monophyletic bryophytes (sN) as well as the associated sCF, SDF1, and SDF2 following the step-wise removal of outgroup representatives. (b) The change of the sequence saturation level across datasets that recovered the bryophyte monophyly, paraphyly, and undetermined scenarios. (c) The distinct clock-likeness across trees that recovered the bryophyte monophyly, paraphyly, and undetermined scenarios.

##### Fast-evolving sites and site heterogeneity jointly confound the embryophyte deep divergence

To unravel the cause of the bryophyte paraphyly, we scrutinized species tree inference by 1) selecting sites according to their site-specific rates and 2) selecting branches according to their distances to the root, based on the dataset with *Zygnema* and *Mesotaenium* retained as outgroup representatives in that the bryophyte paraphyly was first recovered in this concatenated dataset. Slow-evolving sites have been proven useful in deciphering deep divergence relationship^63,89,90^. We calculated the maximum likelihood estimation (MLE) of substitution rates for each site on the concatenated MSA dataset (see Methods). For each concatenated MSA, we created 10 subsets, comprising the top 10^th^, 20^th^, 30^th^, 40^th^, 50^th^ percentiles of the slowest-evolving sites, and the top 10^th^, 20^th^, 30^th^, 40^th^, 50^th^ percentiles of the fastest-evolving sites, respectively. In parallel, to examine the effect of long branches, we removed the top 5^th^, 10^th^, 15^th^, 20^th^, 25^th^, 30^th^, 35^th^, 40^th^, 45^th^, 50^th^ percentiles of the longest branches from the concatenated MSA dataset, leading to another 10 subsets (see Methods). Bryophyte paraphyly persists regardless of the removal of long branches (Fig. S5) or fast-evolving sites (Fig. S6). Notably, the species trees inferred from the subsets comprising only the slowest-evolving sites manifest a decaying gradient of the support level for the recovered bryophyte paraphyly (Fig. S6), ranging from 100% UFBoot support in the subsets comprising the top 30^th^ , 40^th^ or 50^th^ percentiles of the slowest-evolving sites, to 77.3% and 56.7% in the subsets comprising the top 20^th^ and 10^th^ percentiles of the slowest-evolving sites, respectively. Whereas the species trees inferred from the subsets comprising only the fastest-evolving sites exhibit conspicuous inconsistencies at numerous deep phylogenetic positions, for instance, failures in recovering the monophyly of any major embryophyte clade (Fig. S6). These results not only reinforce the importance of using slow-evolving sites when resolving deep divergence relationship but also illustrate the lability of the recovered bryophyte paraphyly facing the most conserved slow-evolving sites.

Previous studies have shown that compositional heterogeneity exerts a pronounced effect on the inferred deep divergence relationship^37,91^. To further test the effect of compositional heterogeneity, we applied the C60 mixture model to infer the deep divergence relationship (see Methods). As shown in Fig. S7, when using only the top 10^th^ or 20^th^ percentiles of the slowest-evolving sites, the bryophyte monophyly was recovered; adding more faster-evolving sites would instead recover the bryophyte paraphyly as in the homogeneous profile model. Whereas applying the C60 mixture model consistently failed to recover the bryophyte monophyly regardless of how many long branches were removed (Fig. S8). Altogether, these results point to a joint confounding effect of fast-evolving sites and site heterogeneity on the phylogenetic signal regarding the embryophyte deep divergence. Importantly, removing the long branches while retaining the fast-evolving sites appears ineffective in the recovery of deep phylogenetic signals even under a site-heterogeneity model.

#### Outgroup-free rooting

##### Non-reversible model lends support for the bryophyte-root scenario

Non-reversible model integrates the root position as a parameter in the inference of ML gene tree^68^, which makes alternative rooting scenarios directly comparable. Leveraging the non-reversible model implemented in IQTREE^92^, we calculated the rooting likelihood on every possible branch on the concatenated MSA (see Methods). The result shows that the likelihood reaches its maximum when rooted with bryophytes (Table 1; Fig. 4). Applying the rule-of-thumb for model selection^93^ in terms of ΔAIC, the second- and third-best root positions at hornworts and setaphytes have considerably weaker support than the ML root position (i.e., bryophytes; Table 1; Fig. 4). The fourth-best root position at mosses has very little support compared to the ML root position (Table 1; Fig. 4). The fifth- to ninth-best root positions have essentially no support compared to the ML root position (Table 1; Fig. 4). We further calculated the root likelihood on individual MSAs of the 273-rSOG and 2026-RefSOG datasets across five frequently debated rooting scenarios (i.e., the bryophyte-root, hornwort-root, liverwort-root, moss-root, and setaphyte-root scenarios; see Methods). Analysis on the individual MSAs of the 273-rSOG dataset shows that the number of gene families inferred with the highest likelihood for the bryophyte-root and moss-root scenarios is significantly greater than expected by chance (Bonferroni-adjusted *P* = 5.187 × 10□^12^ and 0.025, respectively; binomial test; Table 2), whereas it is significantly lower for the liverwort-root and setaphyte-root scenarios (Bonferroni-adjusted *P* = 0.020 and 3.121 × 10□^12^, respectively; Table 2). Further, the bryophyte-root scenario obtained significantly more support than the moss-root scenario (*P* = 0.005, Fisher’s exact test). Analysis on the 2026-RefSOG dataset shows that the number of gene families inferred with the highest likelihood for the bryophyte-root scenario is tremendously greater than expected by chance (Bonferroni-adjusted *P* = 4.039 × 10□^94^; Table 2), whereas it is significantly lower for all other scenarios (Bonferroni-adjusted *P* = 0.004, 2.932 × 10□^4^, 1.206 × 10□^7^ and 1.772 × 10□^25^ for the hornwort-root, liverwort-root, moss-root, and setaphyte-root scenarios, respectively; Table 2). Altogether, the analyses leveraging the non-reversible model bolster the bryophyte-root scenario.

**Figure 4.**
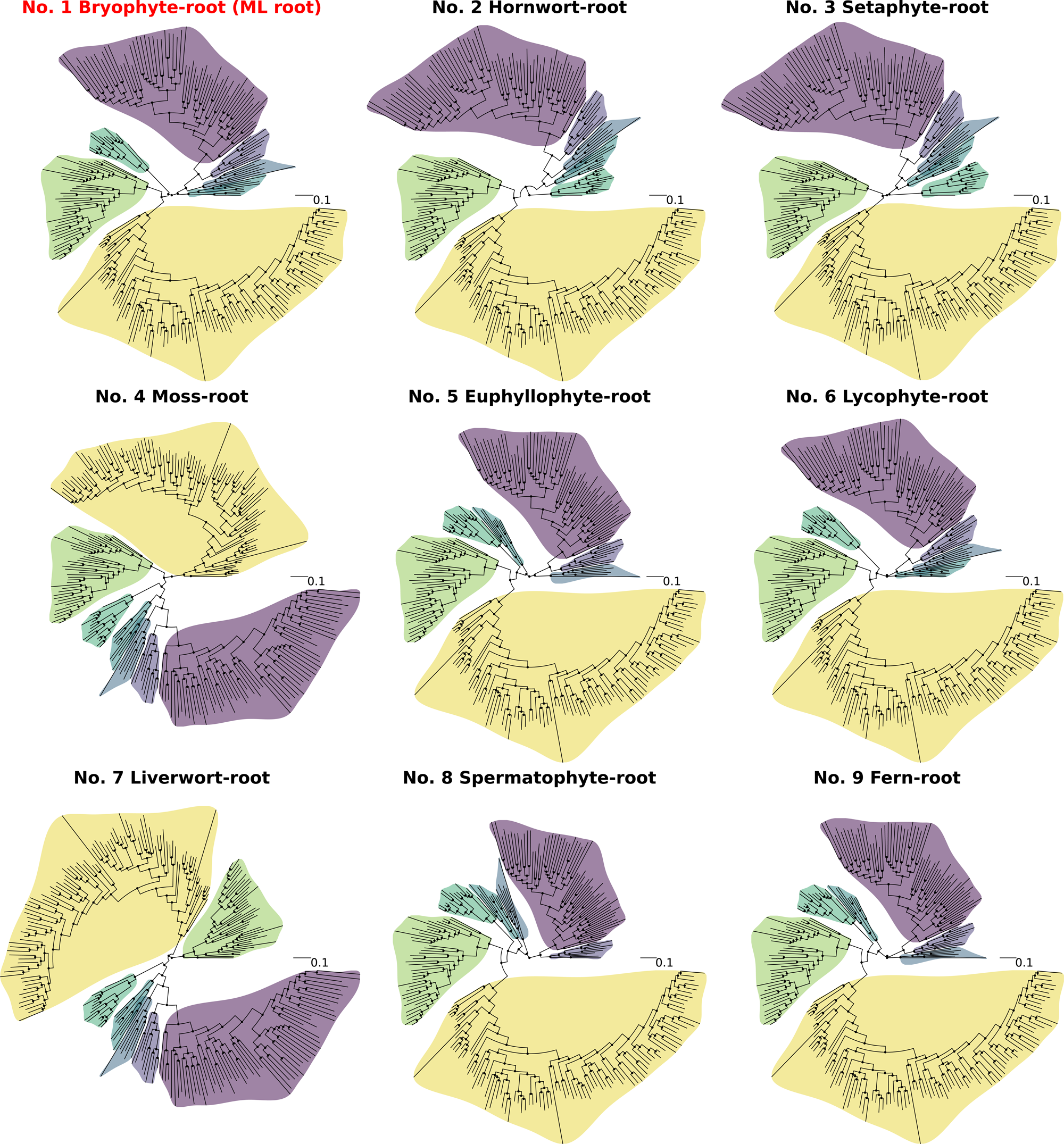
The top 9 best root positions estimated by the non-reversible model based on the concatenated MSA of the 273-rSOG dataset. The top 9 best root positions in terms of likelihood and the associated ML gene trees are shown with their rankings. Major embryophyte clades are highlighted in distinct colors using the same color code as in Fig. 2. The ML root position (i.e., the bryophyte-root) is marked in red. Detailed likelihood information can be found in Table 1.

**Table 1.**
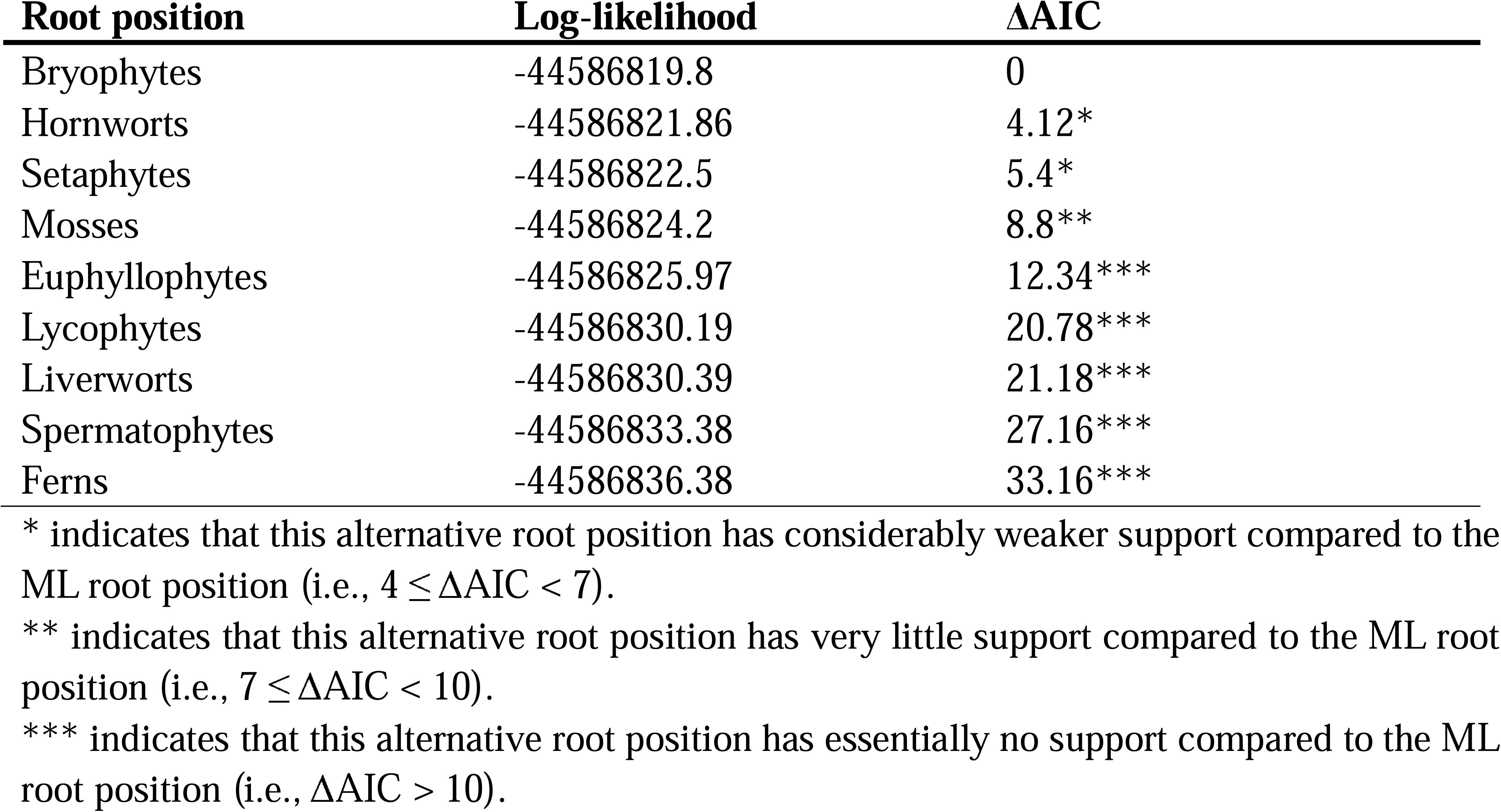
Log-likelihood of the top 9 best root positions and their ΔAIC with the ML root position.

**Table 2.**
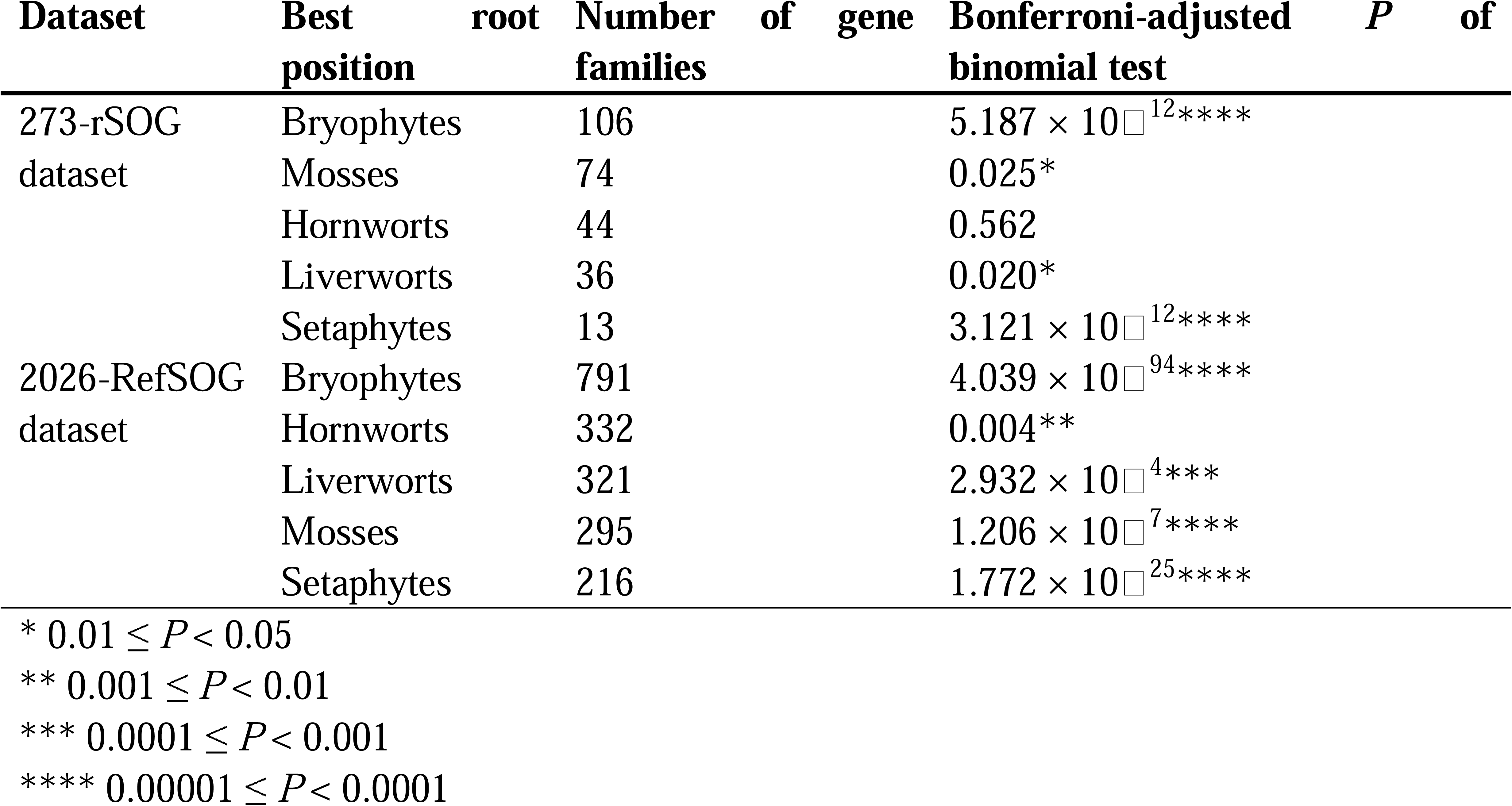
The number of gene families supporting different root positions.

##### MAD rooting supports the bryophyte-root scenario despite considerable phylogenomic discordance

We further applied the MAD^65^ rooting method on unrooted gene trees to find the best root position. Unrooted gene trees were inferred on the 273-rSOG and 2026-RefSOG datasets based on different substitution models (see Methods). In total, there are 15 possible topologies regarding the interrelationship of hornworts, mosses, liverworts, and tracheophytes (Fig. 5a). Notably, we found that each distinct topology is indeed supported by a number of gene families (Table 3; Fig. 5b). For the unrooted gene trees inferred from the 273-rSOG dataset, the chi-square goodness-of-fit test shows that the numbers of gene families supporting different topologies are significantly different in all applied substitution models (*P* = 2.842 × 10^−29^, 6.409 × 10^−28^, 1.894 × 10^−19^ for the LG model, BIC-best model, and the C20 mixture model, respectively). The number of gene families with the root position inferred at the crown node of bryophytes is consistently larger than expected by chance (Table 3; Fig. 5b). Whereas the numbers of gene families with the root position inferred at other nodes are consistently no difference or smaller than expected by chance (Table 3; Fig. 5b). In particular, regarding the 3 possible interrelationships within bryophytes, the one with hornworts being sister to setaphytes (i.e., T1 in Fig. 5a) is consistently favored by the largest numbers of gene families (Table 3; Fig. 5b). It is noteworthy that applying the C20 mixture model will increase the number of gene families supporting the other 2 alternative interrelationships within bryophytes, implying the confounding effect of compositional heterogeneity across sites on the deep divergence signal of individual gene families. Nevertheless, the outstanding support for T1 still holds after accounting for compositional heterogeneity. For the unrooted gene trees inferred from the 2026-RefSOG dataset, the numbers of gene families supporting different topologies are also significantly different (*P* < 1 × 10^−223^ for all applied models; chi-square goodness-of-fit test). Likewise, the number of gene families supporting T1 is also consistently the largest among the 15 possible scenarios and significantly larger than expected by chance (Table 3; Fig. 5b). Compared to the results based on the 273-rSOG dataset, the results based on the 2026-RefSOG dataset differs in the liverwort-root scenario which is consistently supported by significantly more gene families than expected by chance (Bonferroni-adjusted *P* = 1.070 × 10^−5^, 7.650 × 10□^7^, 8.101 × 10□^6^, for the LG model, BIC-best model, and the C20 mixture model, respectively; binomial test; Fig. 5b), which points to the standing phylogenomic discordance across more sampled gene families. Above all, the bryophyte-root scenario exclusively stands the tests with varied sampling of gene families and varied substitution models.

**Figure 5.**
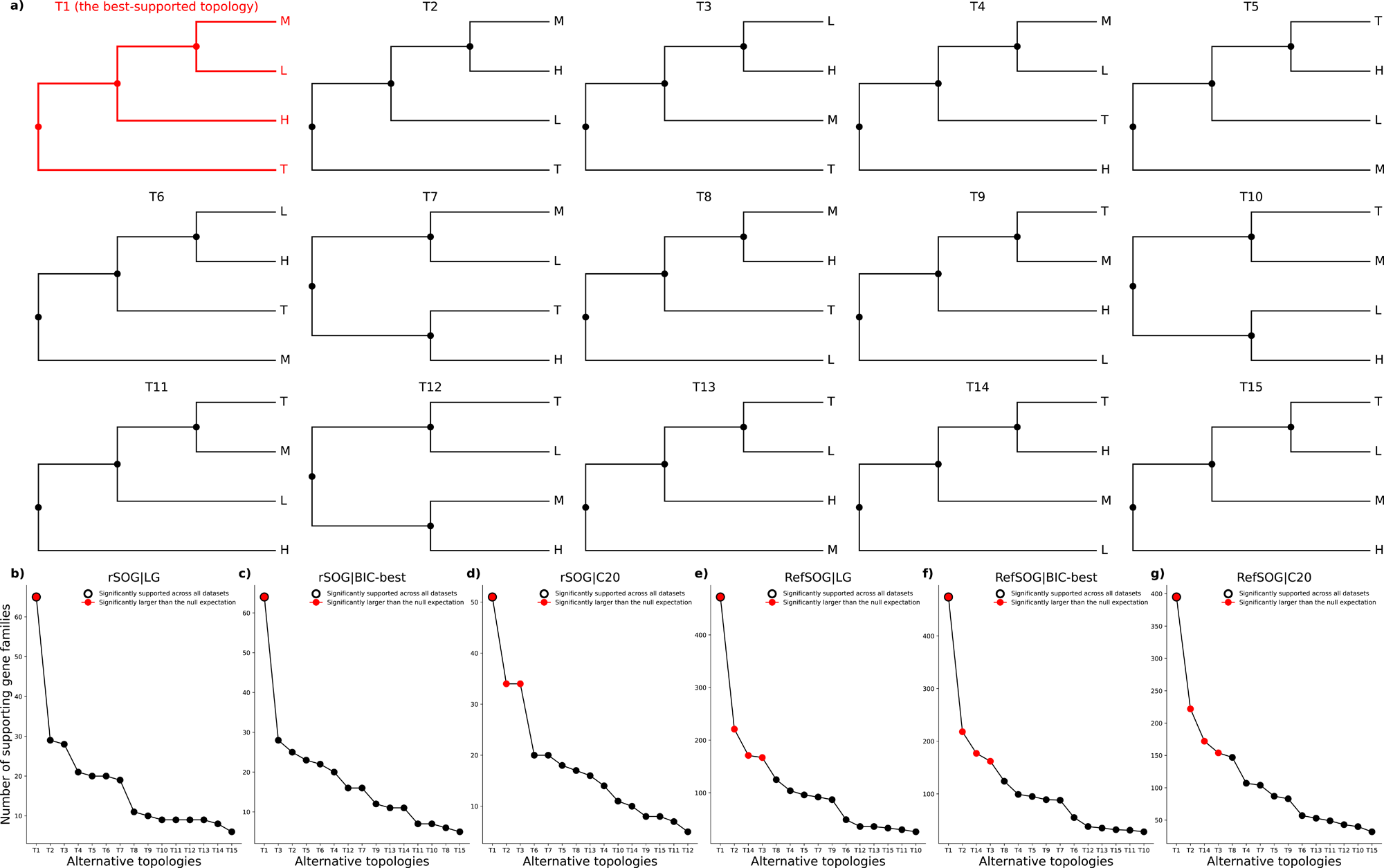
The 15 distinct rooted topologies inferred by MAD and the number of supporting gene families of the 273-rSOG dataset and the 2026-RefSOG dataset. (a) The 15 distinct rooted topologies inferred by the MAD method. The best-supported topology (i.e., T1) is highlighted in red with thicker lines. (b) The number supporting gene families across different datasets. Topologies that received significantly more support than the null expectation are highlighted in red. The topology that received significantly more support than the null expectation across all datasets (i.e., T1) is marked with a black ring. Detailed information of the significance level can be found in Table 3.

**Table 3.**
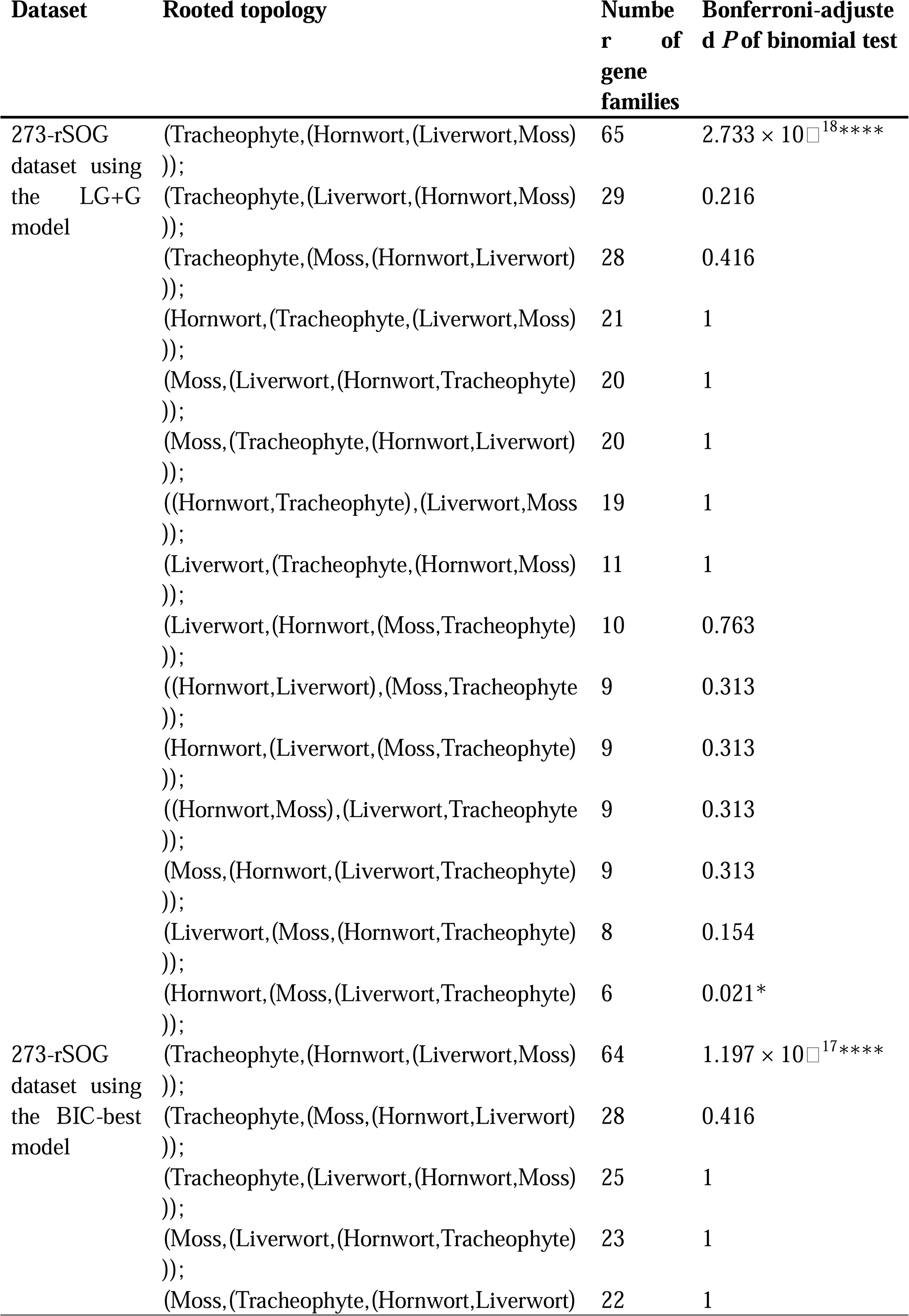
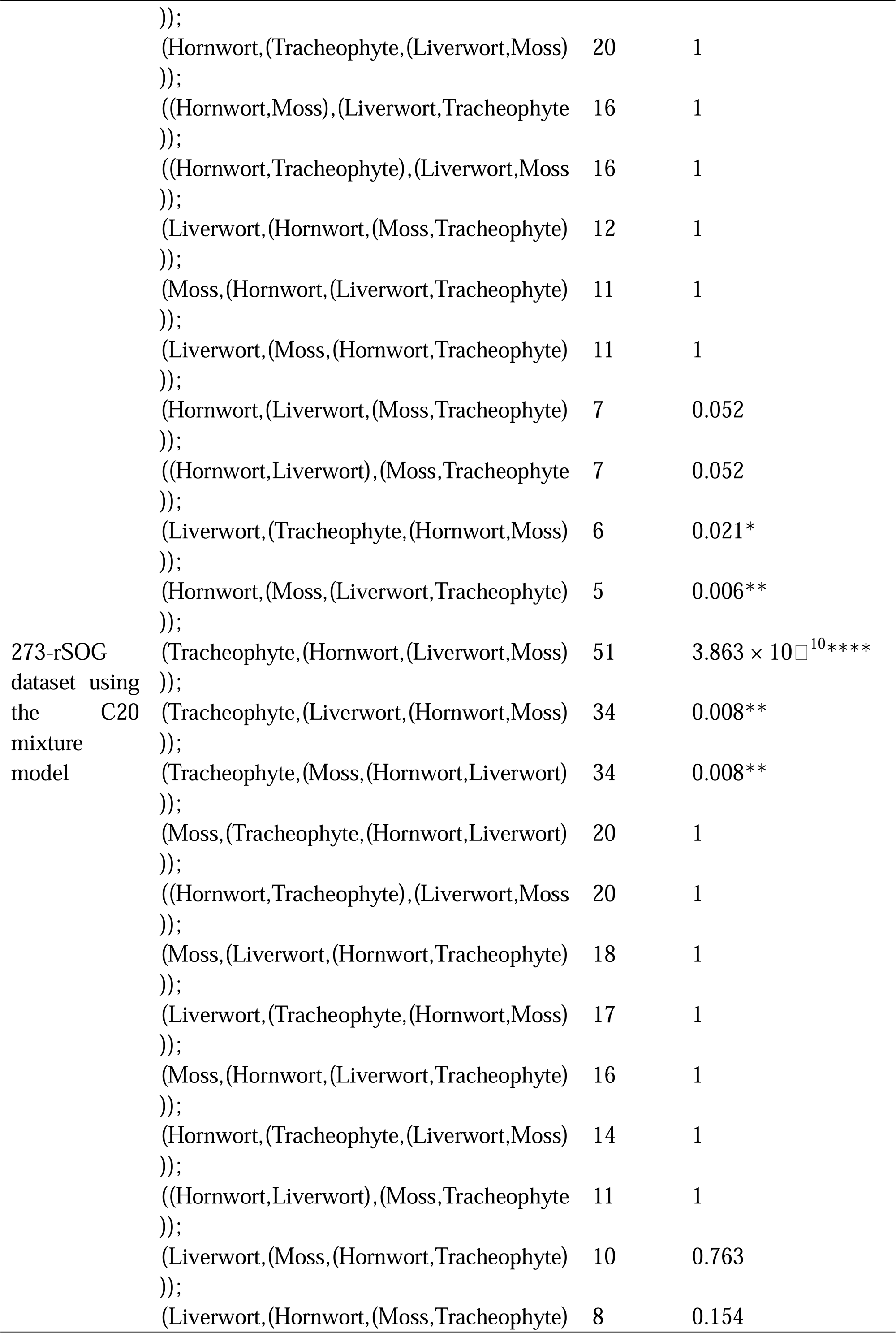
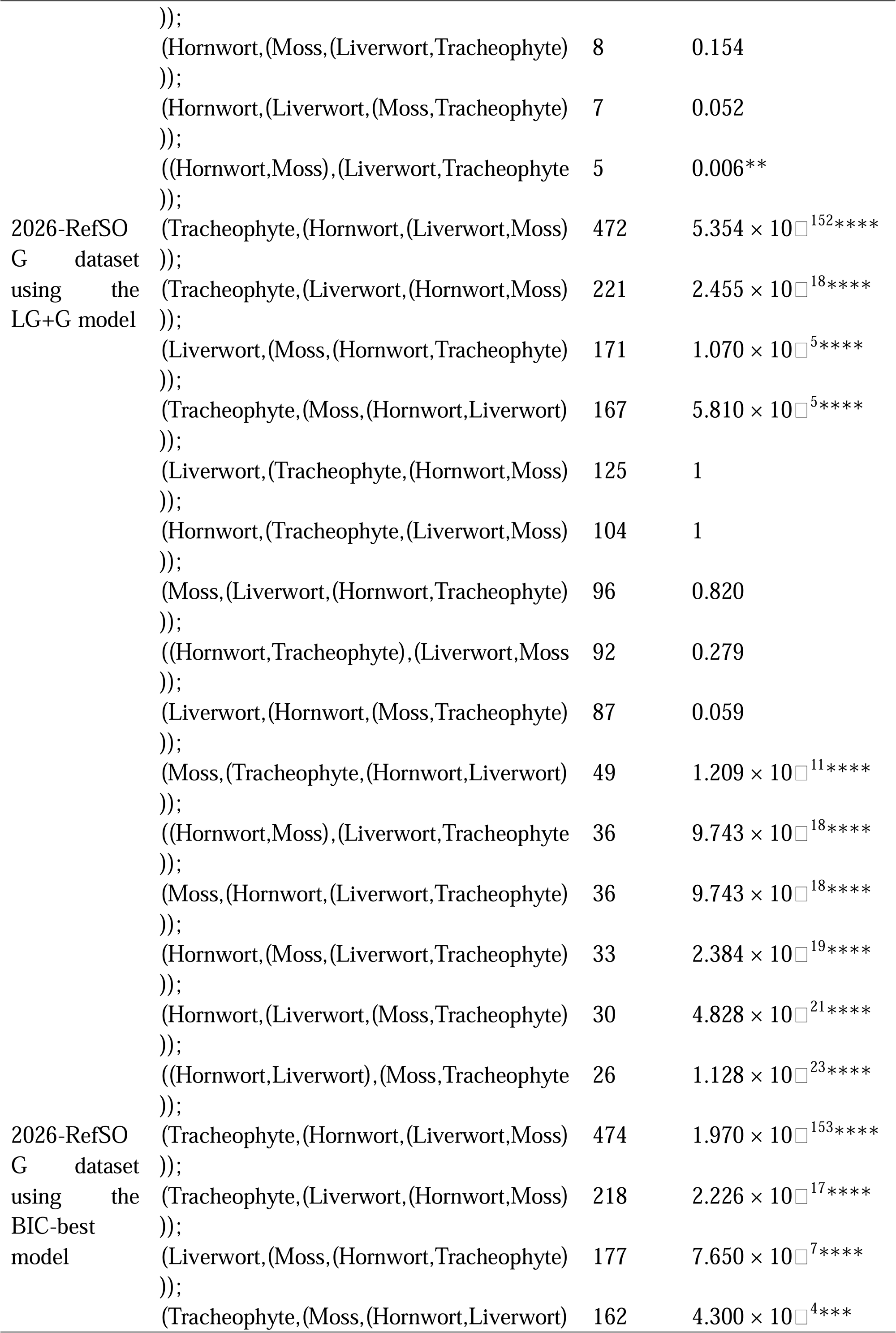
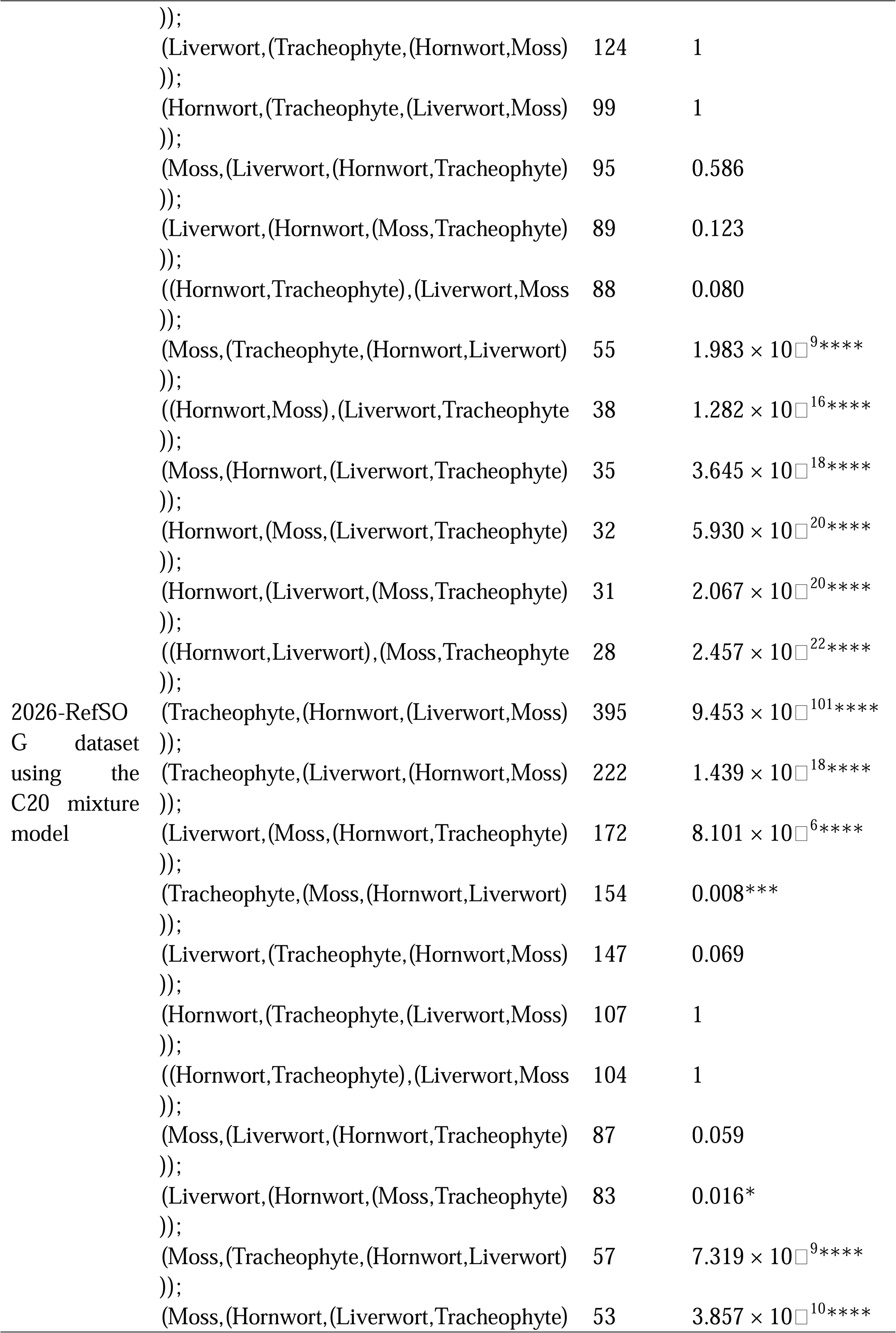
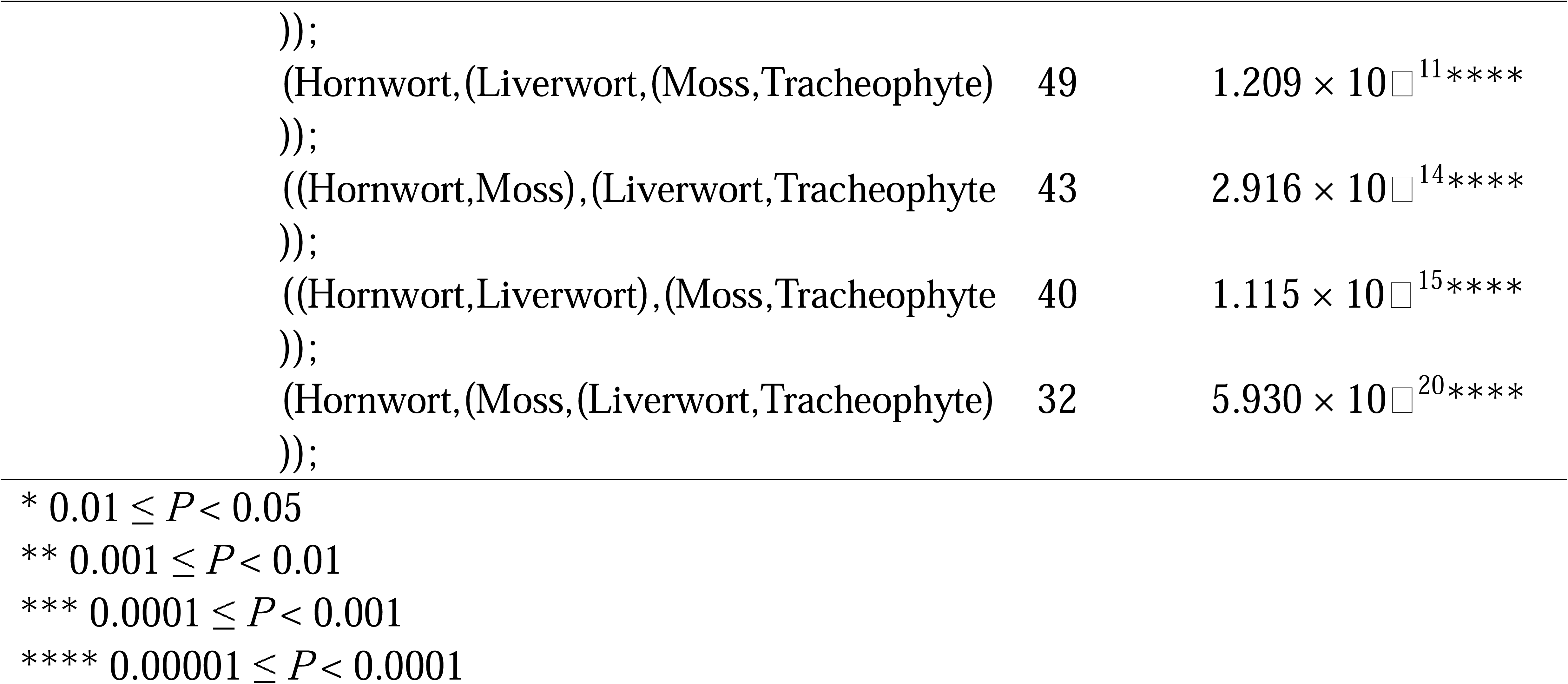
The number of gene families supporting different rooted topologies.

##### Paralogue-based rooting supports the bryophyte-root scenario

Rooting by paralogues (i.e., duplicated genes) can be traced back to 1980s^69^ when Iwabe et al.^69^ utilized a few gene pairs that were assumed to be duplicated predating the divergence of the last universal common ancestor to determine the interrelationship of the three primary domains (i.e., Eukarya, Bacteria, and Archaea). As for embryophytes, despite the fact that gene duplication is pervasive among different major clades^94,95^, established episodes of gene duplication that emerged unequivocally predating the divergence of embryophytes are scarce. Auxin response factors (ARFs) are crucial components of the nuclear auxin response in embryophytes^96–98^. ARFs can be divided into three major clades (i.e., clade A, clade B, and clade C), originating from ancient gene duplications that predate the divergence of embryophytes^97,99–101^. In particular, the presence of clade-B ARFs in hornworts is under question^99–101^. We thus opted to retrieve paralogous gene sequences of clade-A and clade-C ARFs from our 243-genome dataset (see Methods). Given that paralogue-based rooting studies often suffer from the low resolution and divergent functional evolution of individual paralogues, giving rise to discordant topologies across different paralogues^69,102^, we constructed a concatenated composite alignment consisting of clade-A and clade-C ARFs to decipher the root of embryophytes using the LG and the BIC-best model (see Methods). Both models recovered the same topology (Extended Data Fig. 3 and Fig. S9). As expected, the inferred ML gene tree utilizing the BIC-best model fully supports the monophyly of clade-A and clade-C ARF sequences (Extended Data Fig. 3). The bryophyte-root scenario is recovered for both clade-A and clade-C ARF subtrees, despite the lack of full support in the clade-A ARF subtree both in the BIC-best model (Extended Data Fig. 3) and the LG model (Fig. S9). We further applied the C60 mixture model, which again recovered the same topology but markedly increased the supporting level of the undetermined crown node of bryophytes in the clade-A ARF subtree (Extended Data Fig. 3).

## Discussion

### Overlooked outgroup effect on the embryophyte deep divergence and its causes

Since its formalization by Watrous and Wheeler^103^, outgroup rooting has remained a fundamental and widely used method for rooting gene trees and species trees^52–55^, which requires credible prior information of the phylogenetic relationship of the outgroup as to the ingroup. Inappropriate sampling of outgroups^104^, for instance, outgroups that are too distantly related to ingroups, can bias the topology of ingroups due to factors such as LBA^57,105^, homoplasy^106,107^ and compositional shift^61,108^. Nonetheless, little is known about the effect of sampled algal outgroups on the deep divergence relationship of embryophytes.

Using a stepwise outgroup-removal strategy on the 137-rSOG dataset (see Methods), we uncovered a strong algal outgroup effect that can obscure deep phylogenetic signals within embryophytes (Extended Data Fig. 2). When outgroups from multiple major clades were retained, concatenation analyses consistently recovered bryophyte monophyly with full support. As outgroups were reduced within Zygnematophyceae, support for bryophyte monophyly declined gradually and then shifted abruptly to strongly supported bryophyte paraphyly; throughout, both the number of sites informative for the bryophyte crown node and the corresponding sCF decreased steadily (Fig. 3a). Notably, once the sCF fell below the one-third null expectation, support for bryophyte monophyly dropped concurrently, indicating that outgroup choice can qualitatively alter ingroup topology when decisive phylogenetic signal falls below a critical threshold. This is relevant especially when the degree of phylogenetic discordance across genes and sites is high (see further). Alongside the declined number of decisive sites, we also find that the level of substitution saturation increased and the tree clock-likeness decreased in the inferred gene trees (Fig. 3b-c). The increased level of saturation suggests that removing an outgroup representative may not only risk reducing the number of decisive sites but also risk reducing the overall informativeness of sites, for instance, when the removed outgroup representative held crucial sites informative for the determination of ancestral state and/or multiple hits. Declined clock-likeness signals an elevated level of rate heterogeneity across branches, which increases susceptibility to LBA where fast-evolving lineages with elongated branches may spuriously cluster together due to homoplasy^109^ or saturation^110^; the inclusion of fast-evolving sites (Fig. S6) and the inadequate accounting for site-heterogeneity (Fig. S7) have jointly confounded the deep phylogenetic signal of embryophytes. Using the C60 mixture model, we recovered monophyletic bryophytes (Fig. S7), which the homogeneous profile model failed to do (Fig. S6) on the datasets comprising the top 10^th^ or 20^th^ percentiles of slowest-evolving sites. On the other hand, we find that simply removing the longest branches while keeping all the fast-evolving sites in place could not recover the bryophyte monophyly no matter how many long branches were removed (Fig. S5) even under the site-heterogeneity model (Fig. S8). Importantly, the ever-growing availability of genomes in the genomic era, increasing taxon sampling, does not necessarily improve the phylogenetic inference for deep nodes. Instead, increased taxon sampling must be combined with meticulous curation of conserved and informative sites and adequate accounting for the accumulated compositional heterogeneity and saturation through deep time to improve the robustness of the recovered deep divergence over different sampling strategies, datasets, and models.

Diverse biological processes can underpin the observed phylogenomic discordance across gene families. We find that it is the ILS rather than gene flow that can plausibly explain the observed triplet frequency distribution (Extended Data Fig. 1a-c). In particular, the observed discordant level aligns with the expectation of a moderate level of ILS (Extended Data Fig. 1d): the concatenation method that combines all the concordant and discordant sites across gene families still holds promise of recovering the correct deep phylogeny under such moderate level of model violation (Extended Data Fig. 2a).

### Bryophyte monophyly stands various tests of outgroup-free rooting despite pervasive phylogenomic discordance

Although with the outgroup-rooting method we successfully recovered the bryophyte monophyly with a string of meticulous analyses with various datasets and models, as in many other recent large-scale phylogenetic studies of embryophytes^31–37^, it does not fundamentally answer the initial question we posed—could the bryophyte-root scenario simply be an artefact resulting from algal outgroups? To address this, we applied three distinct outgroup-free rooting methods (i.e., the non-reversible model-based rooting, MAD rooting, and paralogue-based rooting) to our 273-rSOG dataset and 2026-RefSOG dataset to determine the optimal embryophyte root position without relying on algal outgroups.

Conventional Markov models of substitutions usually assume stationarity and time-reversibility in which substitutions are equally likely in both directions, which is computationally convenient but leads to unrooted gene trees. The non-reversible model instead integrates the root position as a parameter into the formation of the likelihood function^111^, making different root positions directly comparable. Utilizing the non-reversible model on the concatenated MSA of the 273-rSOG dataset, we show that the ML root position lies on the branch separating bryophytes and tracheophytes, which also significantly outcompetes all the other possible root positions (Table 1; Fig. 4). When further testing on individual gene families five particular rooting scenarios that have recurrently gained support from previous phylogenetic studies^3,6,11,12,112^ (i.e., the bryophyte-root, hornwort-root, liverwort-root, moss-root, and setaphyte-root scenarios), the number of gene families supporting the bryophyte-root scenario has been consistently higher than expected by chance and outcompeted other scenarios in both the 273-rSOG dataset and 2026-RefSOG dataset (Table 2). Notably, for the 273-rSOG dataset, in addition to the bryophyte-root scenario, the moss-root scenario also stood out with support from a significantly large number of gene families (Table 2). Whereas for the 2026-RefSOG dataset, only the bryophyte-root scenario gained significantly strong support (Table 2). It appears that the labile support for the moss-root scenario is caused by the insufficient sampling. At any rate, the presence of the consistently larger number of gene families that support other rooting scenarios than the bryophyte-root scenario in both datasets unequivocally points to the prevalent phylogenomic discordance across gene families. This may explain why numerous previous embryophyte phylogenetic studies relying on several to a few individual marker genes obtained discordant results regarding the root placement. In the presence of pervasive discordant deep phylogenetic signals across gene families, phylogenetic inference based on only a small fraction of the entire available gene repertoire is bound to be sensitive to sampling bias. In that regard, our gene family datasets based on the de novo inference of 243 phylodiverse embryophyte genomes and the latest reference gene family dataset of OrthoDB^82^ should constitute a well-sampled example.

Besides testing alternative rooting scenarios directly by the likelihood under the non-reversible model, we applied the MAD rooting method on the representative four-taxon subtrees derived from unrooted gene trees inferred by different substitution models and datasets (see Methods). The representative four-taxon subtree is formed by the least divergent species-combination of the four major embryophyte clades (i.e., hornworts, mosses, liverworts, and tracheophytes), which is supposed to best preserve the deep phylogenetic signal that can be otherwise blurred by homoplasy and saturation. In theory, there can be up to 15 different rooted topologies regarding a four-taxon bifurcating tree (Fig. 5a). In practice, all the 15 alternative rooted topologies have been recovered by MAD with different numbers of gene families from both the 273-rSOG dataset and 2026-RefSOG dataset under different substitution models (Table 3; Fig. 5b). The topology T1 (i.e., bryophytes form a monophyly with hornworts sister to setaphytes) is consistently recovered as the best topology regardless of which substitution model and which dataset was used. Nonetheless, there are subtle differences between the results derived from different substitution models and datasets. For the 273-rSOG dataset, when applying the LG model or the BIC-best model, only T1 stood out as the significantly supported topology, while applying the C20 mixture model decreased the support for T1 but increased the support for T2 and T3 (Table 3; Fig. 5b), both of which still represent the bryophyte monophyly but differ in the interrelationship within bryophytes. This signifies the presence of pronounced compositional heterogeneity among some gene families that could biasedly cluster lineages with convergent structural and/or functional constraints together. On the other hand, analyzing the 2026-RefSOG dataset did not reveal a prominent effect of compositional heterogeneity in that the significantly supported rooting scenarios remain the same across different substitution models (Table 3; Fig. 5b), although the number of gene families supporting T1 also declined when applying the C20 mixture model. This indicates that despite the confounding effect of compositional heterogeneity among some gene families, sufficient gene sampling can overcome these noises. Nevertheless, in addition to T1, T2, and T3, results derived from the 2026-RefSOG dataset show that T14 has consistently emerged as one of the significantly supported topologies (Table 3; Fig. 5b), i.e., liverworts being sister to the rest of embryophytes. This result is not totally unexpected. As increasing the number of sampled genes from 273 to 2,026 will inevitably cover more genes in more functional categories, the known reductive evolution underwent by liverworts^32^ that distinguishes their selection profiles and structural constraints from other embryophytes probably gives rise to a number of genes supporting the liverwort-root scenario^11,78^, which do not necessarily reflect the true underlying species tree topology but the entangled domain and/or sequence composition alongside their unique reductive evolution. This constitutes another plausible biological explanation for the prevalent phylogenomic discordance across gene families, especially for deep nodes whose signals have already been erased by factors such as multiple hits and homoplasy.

Gene duplication is a defining feature of embryophytes^34^. Nonetheless, well-characterized gene pairs that were duplicated predating the divergence of embryophytes are scarce. We leveraged the established clade-A and clade-C ARFs to perform a paralogue-based rooting analysis. Previous paralogue-based rooting studies have often reported discrepant rooting placements across individual paralogues and gene families, probably due to the low phylogenetic signal and the divergent alignment profile across individual genes. We applied a novel method that concatenates orthologues from different species and preserves the local alignment profiles of different paralogues in the final global alignment profile to construct a concatenated composite alignment that magnifies the phylogenetic signal while accounting for the local structural variation across paralogues (see Methods). Concordant results have been obtained for both the clade-A and clade-C ARF subtrees in which the bryophyte-root scenario is supported (Extended Data Fig. 3 and Fig. S9). Notably, the bryophyte crown node in the clade-A ARF subtree did not obtain full support using the homogeneous profile model (Extended Data Fig. 3 and Fig. S9). While applying the C60 mixture model recovered the same topology but with prominently stronger support for the bryophyte crown node in the clade-A ARF subtree (Extended Data Fig. 3). This reinforces the presence of heterogeneous site-specific substitution profiles of deep nodes, probably reflecting the divergent selection pressure and functional constraints across different paralogues through deep time.

## Conclusions

The current prevailing bryophyte monophyly topology can be robustly recovered across a range of algal outgroup settings and species tree inference methods, yet a remarkable outgroup effect that can significantly alter the deep phylogeny of the embryophyte ingroup. Using slow-evolving sites and site-heterogeneity model can mitigate such outgroup effect and restore the deep phylogenetic signal. In parallel, the bryophyte monophyly scenario exclusively stands the tests of various outgroup-free rooting methods among a range of competing deep divergence scenarios, addressing the initial question we proposed that the bryophyte monophyly could be a mere artefact by the introduction of outgroups. Alongside different lines of evidence supporting the bryophyte monophyly scenario, we demonstrate nonetheless that the presence of pervasive phylogenomic discordance derived from biological processes such as ILS and methodological issues such as model inadequacy constitutes an essential reason for the long-lasting debate over the deep divergence of embryophytes. Using this much expanded sampling of hornworts, mosses, and liverworts—balancing out the abundantly sampled tracheophytes—and methodological rigour we provide a framework that robustly roots the embryophytes at their deepest divergence, yielding monophyletic Tracheophyta and monophyletic Bryophyta. Endorsing the monophylum Bryophyta has profound implications for our understanding of plant evolution of form and function: character states uniquely found in tracheophytes and bryophytes have an equal likelihood to reflect the state in the last common ancestor of land plants. Here, the use of algal sampling comes full circle, because only they can resolve this stalemate of trait inference.

## Methods

### Taxon sampling

Leveraging the accumulated wealth of publicly available embryophyte genomes, particularly the recent release of numerous bryophyte genomes^35,113^, we constructed a large-scale embryophyte genome dataset encompassing 154 bryophyte species composed of 15 hornwort, 40 liverwort, and 99 moss species, and 89 tracheophyte species composed of 9 lycophyte, 7 fern, 10 gymnosperm, and 63 angiosperm species. Our sampling of tracheophyte genomes covers all the orders that currently have published genomes. We further collated genome data of 12 algal outgroup representatives that include *Chlamydomonas reinhardtii* from Chlorophyceae, *Klebsormidium nitens*, *Chlorokybus atmophyticus*, and *Mesostigma viride* from the KCM grade, *Chara braunii*, *Spirogloea muscicola*, *Mesotaenium endlicherianum*, *Penium margaritaceum*, *Zygnema cf. cylindricum*, *Z. circumcarinatum SAG 698-1b*, *Z. circumcarinatum UTEX 1559*, and *Z. circumcarinatum UTEX 1560* from the ZCC grade. The source of data is summarized in Table S2.

### Construction of single-copy orthologous gene datasets

We constructed our single-copy orthologous (SOG) datasets leveraging 1) de novo inferred orthogroups (i.e., orthologous gene families) using OrthoFinder (v2.5.4)^81^ and 2) reference-based SOGs from OrthoDB (v12)^82^. For the de novo inferred orthogroups, we first conducted the gene family clustering analysis employing OrthoFinder with the inflation factor set as 3 and other parameters set as default based on protein sequences. We ran the gene family clustering analysis two times. The first run is without algal outgroup representatives, i.e., only 243 embryophyte genomes, referred to as the 243-genome dataset. The second run is with all algal outgroup representatives, i.e., 243 embryophyte genomes with 12 algal genomes, referred to as the 255-genome dataset. Both runs ended up with no SOG deduced, probably due to the pervasive episodes of gene and genome duplications among embryophytes^34^. We thus opted to construct refined SOGs (rSOGs) from multi-copy orthogroups. The procedure is as follows. We first select orthogroups with 100% species coverage and conduct an all-against-all sequence similarity search of each orthogroup to itself using diamond (v2.1.11.165)^114^. For multiple gene copies from the same species, we selected the gene that has the highest overall bit-score to other species as the representative gene for this species. While for species that only have one gene copy, that gene is then selected as the representative gene. Such that we constructed our rSOG datasets, with 273 rSOGs for the embryophyte-only genome dataset and 137 rSOGs for the genome dataset including algae, referred to as the 273-rSOG and 137-rSOG dataset, respectively. For the reference-based SOGs, we leveraged the 2,026 SOGs (referred to as the 2026-RefSOG dataset) curated in the embryophyta_odb12 dataset within the latest OrthoDB.v12 BUSCO dataset^82^. In brief, we first scanned the hidden Markov model (HMM) profile database of the 2,026 reference SOGs against our 243 embryophyte genomes using hmmscan from hmmer (v.3.3.2)^115^ with the e-value threshold set as 1 × 10^−10^. Using the bit-score cutoffs provided by the embryophyta_odb12 dataset, we then filtered out homologous genes that had lower bit-scores than the cutoff for each reference SOG. For the species with multiple homologous genes remained, we retained the gene that had the highest bit-score as the representative orthologue for that species. Such that we constructed our 2,026 reference SOGs.

### Construction of paralogous gene datasets

The emergence of the auxin response factor (ARF) clade A and clade C is originated from an ancient duplication event predating the divergence of embryophytes^97,99–101^. Tracing by the gene ids of *Arabidopsis thaliana*, we retrieved the two corresponding gene families of the ARF clade A and clade C from our gene family profile. We performed an all-against-all sequence similarity search for each of them as to itself using diamond (v2.1.11.165)^114^. For each ARF clade, we selected for each of the four major embryophyte clades (i.e., hornworts, mosses, liverworts, tracheophytes) the top gene from 15 different species (i.e., the species number of the least sampled hornwort clade) in terms of the sequence similarity represented by the bit-score with other species, leading to in total 60 genes per ARF clade. PRANK (v170427)^116^ was used to construct multiple sequence alignment (MSA) for each ARF clade with default parameters. We then concatenated for each of the four major embryophyte clades the 15 aligned protein sequences for each ARF clade, leading to 4 concatenated sequences per ARF clade. Clustal-Omega (v1.2.4)^117^ was then applied to infer a composite alignment from the pre-aligned multiple sequence profiles of the ARF clade A and clade C so as to preserve the local alignment profile of each paralogue.

### Maximum likelihood inference of gene trees

For the 137-rSOG, 273-rSOG and 2026-RefSOG datasets, we conducted the maximum likelihood (ML) gene tree inference analyses as follows. We first constructed MSAs based on amino acid sequence for each SOG using PRANK (v170427)^116^ with default parameters. The resulting peptide MSAs were then used as input for IQTREE (v3.0.1)^92^ to perform gene tree inference. We ran IQTREE two times. The first run applied the commonly used LG+G model. The second run applied the ModelFinder^83^ to select the best model among the 1,232 alternative protein substitution models implemented in IQTREE in terms of BIC (i.e., the BIC-best model). We launched an additional run that applied the LG+C20+F+G+PMSF model which accounts for among-site compositional heterogeneity on the outgroup-free 273-rSOG and 2026-RefSOG datasets. The phylogenetic uncertainties are measured by 1,000 ultrafast bootstrap replicates^118^, SH-like approximate likelihood ratio test^119^, approximate Bayes test^120^, and the local bootstrap probability^121^, optimized by the nearest neighbor interchange. For the 137-rSOG dataset, we further concatenated all individual MSAs into a supermatrix and then repeated the ML gene tree inference analyses. We conducted the step-wise removal of outgroup representatives from the supermatrix and repeated the ML gene tree inference analyses in the order of *Chlamydomonas reinhardtii*, *Chlorokybus atmophyticus*, *Mesostigma viride*, *Klebsormidium nitens*, *Chara braunii*, *Spirogloea muscicola*, *Penium margaritaceum*, *Mesotaenium endlicherianum*, *Zygnema cf. cylindricum*, *Z. circumcarinatum UTEX 1560*, and *Z. circumcarinatum SAG 698-1b*, following a distant-to-close order in terms of the phylogenetic distance according to the latest algal phylogenetic system^122–124^. The site concordance factor (sCF), site discordance factor for alternative quartet 1 (sDF1) and alternative quartet 2 (sDF2), as well as the number of informative sites decisive for the crown node of monophyletic bryophytes are calculated as the mean over 100 quartets using the updated likelihood-based method^125^ implemented in IQTREE that can account for multiple hits at the same site and mitigate the homoplasy effect. For the composite alignment of paralogues, we conducted the ML gene tree inference analyses same as above, in addition to a third run using the LG+C60+F+G+PMSF model. In particular, we follow the principle that applying the C20 mixture model on individual MSAs and the C60 mixture model on concatenated MSAs to avoid model overfitting.

### Measurement of substitution saturation

Multiple substitutions at the same site can lead to substitution saturation compromising the phylogenetic signal for deep divergence. We calculated the raw proportion of differing sites (i.e., the p distance) and the corrected distance from the inferred ML gene tree for each species pair on the 137-rSOG dataset under different outgroup settings (see above). For the dataset of each outgroup setting, we fit a Michaelis–Menten saturation curve of the p distance against the corrected distance using the ‘curve_fit’ function implemented in the SciPy (v1.5.4)^126^ library. The saturation level is then calculated as the maximum p distance divided by the estimated asymptotic maximum. We also fit an alternative exponential saturation curve but abandoned it in the end due to its worse model fit compared to the Michaelis–Menten saturation curve in terms of *R*^2^ (Table S3).

### Refinement of single-copy orthologous gene datasets

Faster evolving sites are more likely to contain noises from multiple substitutions and turn saturated compared to slower evolving sites^89^, which may lead to systematic bias (e.g., LBA). We calculated the MLE substitution rate for each site of the concatenated MSAs of the 137-rSOG dataset for each outgroup setting using an empirical Bayesian approach^127^ implemented in IQTREE (v3.0.1)^92^ using the best substitution model evaluated by ModelFinder^83^. We then selected the top 10^th^, 20^th^, 30^th^, 40^th^ , 50^th^ percentiles of the slowest-evolving sites and also the top 10^th^, 20^th^, 30^th^, 40^th^ , 50^th^ percentiles of the fastest-evolving sites, respectively, to conduct the ML gene tree inference analyses using the LG+G and the LG+C60+F+G+PMSF model with the same parameters as above. Exceptionally long branches within a tree are known to affect the topology^57^ due to factors such as LBA. Thus, we also removed the top 5^th^, 10^th^, 15^th^, 20^th^, 25^th^, 30^th^, 35^th^, 40^th^, 45^th^, 50^th^ percentiles of the longest branches (i.e., tip-to-root distances), respectively, to conduct the same ML gene tree inference analyses as above.

### Non-reversible model to infer the best root position

Conventional substitution models assume stationarity and reversibility, i.e., substitutions are equally likely in both directions, leading to likelihood functions that are independent on the root position and thus yielding unrooted trees. We utilized the non-reversible model^111^ implemented in IQTREE (v3.0.1)^92^ that takes the root position also as a parameter in the likelihood function to test different rooting scenarios. For the concatenated MSA of the 273-rSOG dataset, we calculated the likelihood of every possible root position. For the individual MSAs in the 273-rSOG and 2026-RefSOG dataset, we tested five particular rooting scenarios (i.e., the bryophyte-root, hornwort-root, liverwort-root, moss-root, and setaphyte-root scenario) by comparing their likelihoods. For clarity, the bryophyte-root scenario entails the phylogenetic tree rooted at the crown node of monophyletic bryophytes, same interpretation for other rooting scenarios. We conducted a binomial test to determine which root position is significantly more supported than expected by chance, assuming that each of the 5 alternative root positions has an equal probability (i.e., 0.2) of being the best. Bonferroni correction is performed to control for multiple comparisons.

### Four-taxon minimal ancestor deviation rooting

Grounded in clock-like reasoning, the rooting method of minimal ancestor deviation (MAD)^65^ estimates the root position by considering all branches and quantifies the root-mean-square (RMS) of the pairwise relative deviations from the molecular clock expectation at different positions of each branch. The optimal root position inferred by MAD is the branch and position that minimizes such deviation. Four-taxon unrooted gene trees have been shown effective for an accurate MAD rooting^128^ when properly curated. We constructed our four-taxon datasets for each unrooted gene tree inferred by different substitution models (i.e., the LG+G model, BIC-best model, and C20 mixture model) based on the 273-rSOG dataset and 2026-RefSOG dataset. For each SOG, we select one representative species from each of the four major embryophyte clades (i.e., hornworts, mosses, liverworts, tracheophytes) that gave the shortest overall pairwise distance to form a representative four-taxon operating unit, which is most likely to retain ancient phylogenetic signals^89^. The MAD algorithm is then implemented upon the four-taxon datasets to determine the best root position.

### Multispecies coalescent method of species tree inference and the triplet analysis

We inferred species trees from pools of unrooted gene trees using ASTRAL-Pro3 (v1.22.3.6)^129^ which accounts for incomplete lineage sorting (ILS) as modelled by the multispecies coalescence (MSC) model^130,131^. The reliability of ASTRAL-Pro3 is highly dependent on the accuracy of the inferred gene trees. We thus filtered out branches that did not have at least 50% support from any of the uncertainty measures and utilized the retained branches for the species tree inference. The supporting value of each branch in the inferred species tree is measured by the local posterior probability. Default parameters of ASTRAL-Pro3 were applied except for the ‘-u’ option that we set as ‘2’ to produce detailed triplet information for each branch (see further).

We retrieved the triplet frequencies calculated by ASTRAL-Pro3 regarding the crown group of embryophytes, focusing on the interrelationship between the three major embryophyte clades (i.e., hornworts, setaphytes, and tracheophytes). Regarding this rooted three-taxon subtree, the coalescent model^132^ predicts that gene trees arising from the species tree can have three different topologies as illustrated in Extended Data Fig. 1a, in which the dominant topology is regarded as the major triplet (i.e., q1) while the other two minor triplets (i.e., q2 and q3) occur in symmetry. The presence of gene flow between non-sister taxa will give rise to the asymmetry between the two minor triplets due to factors such as exchanging alleles. As such, testing the difference between the two minor triplets can reflect the presence or absence of gene flow^132^. We applied the chi-square goodness-of-fit test to determine whether the frequency of the two minor triplets is significantly different. To alleviate the impact of random sampling error, we generated 200 replicate datasets, each comprising one bootstrapped gene tree randomly sampled per gene family in the 137-rSOG dataset. ASTRAL-Pro3 was used to calculate the frequency of the three alternative triplets on the 200 replicate datasets as above. We then compared the distribution of the frequency of the two minor triplets applying the chi-square test of independence.

To investigate whether ILS alone can explain the observed discordance level across gene families, we applied a simulation-based method following Cai et al. (2021)^87^. In short, we first generated 200 bootstrapped species trees from our 200 replicate datasets using ASTRAL-Pro3 with the default parameters. For each of the bootstrapped species tree, we simulated the same number of gene trees as in the empirical dataset under the coalescent model using the “sim.coal.mpest” function in Phybase^133^. ASTRAL-Pro3 was then used to calculate the frequency of the two minor triplets on the 200 bootstrapped species tree-derived datasets of simulated gene trees with the default parameters. As such, we constructed a null frequency distribution of the discordant triplets simulated under the coalescent model. We then calculated the 1σ confidence interval of the simulated null distribution and made comparison to the observed frequency of the discordant triplets from the 200 replicate datasets.

## Data and code availability

The sequences and MSA profiles of the 137-rSOG, 273-rSOG, and 2026-RefSOG datasets are deposited at https://doi.org/10.6084/m9.figshare.30620423.v1 together with the inferred gene trees by different substitution models. The code to construct rSOGs and curate representative four-taxon datasets can be found in the GitHub repository heche-psb/RootingLandPlant.

## Acknowledgements

H.C. acknowledges funding from the Walter Benjamin Programme of the German Research Foundation (DFG, project number 566819120, CH 3905/1-1). J.d.V. is grateful for funding by the German Research Foundation grant 509535047 (VR 132/10-1) and grants 440231723 (VR 132/4-1) and 528076711 (VR 132/13-1) within the framework of the Priority Programme “MAdLand—Molecular Adaptation to Land: Plant Evolution to Change” (SPP 2237). J.d.V. further thanks the European Research Council for funding under the European Union’s Horizon 2020 research and innovation programme (grant agreement no. 852725; ERC-StG “TerreStriAL”). We thank Profs. Fai-Wei Li from Boyce Thompson Institute and Günter Theißen from FSU Jena for their precious suggestions and insightful discussion on this study during the MAdLand Annual Meeting 2025.

## Author contributions

H.C. and J.d.V. conceived and managed this project. H.C. conducted the analysis in this study. J.d.V. supervised this project. H.C. and J.d.V. wrote the manuscript. All authors read and approved the manuscript.

