## Supplementary figures and images for "Rooting the deep divergence of land plants"

### Extended Data Figure 1

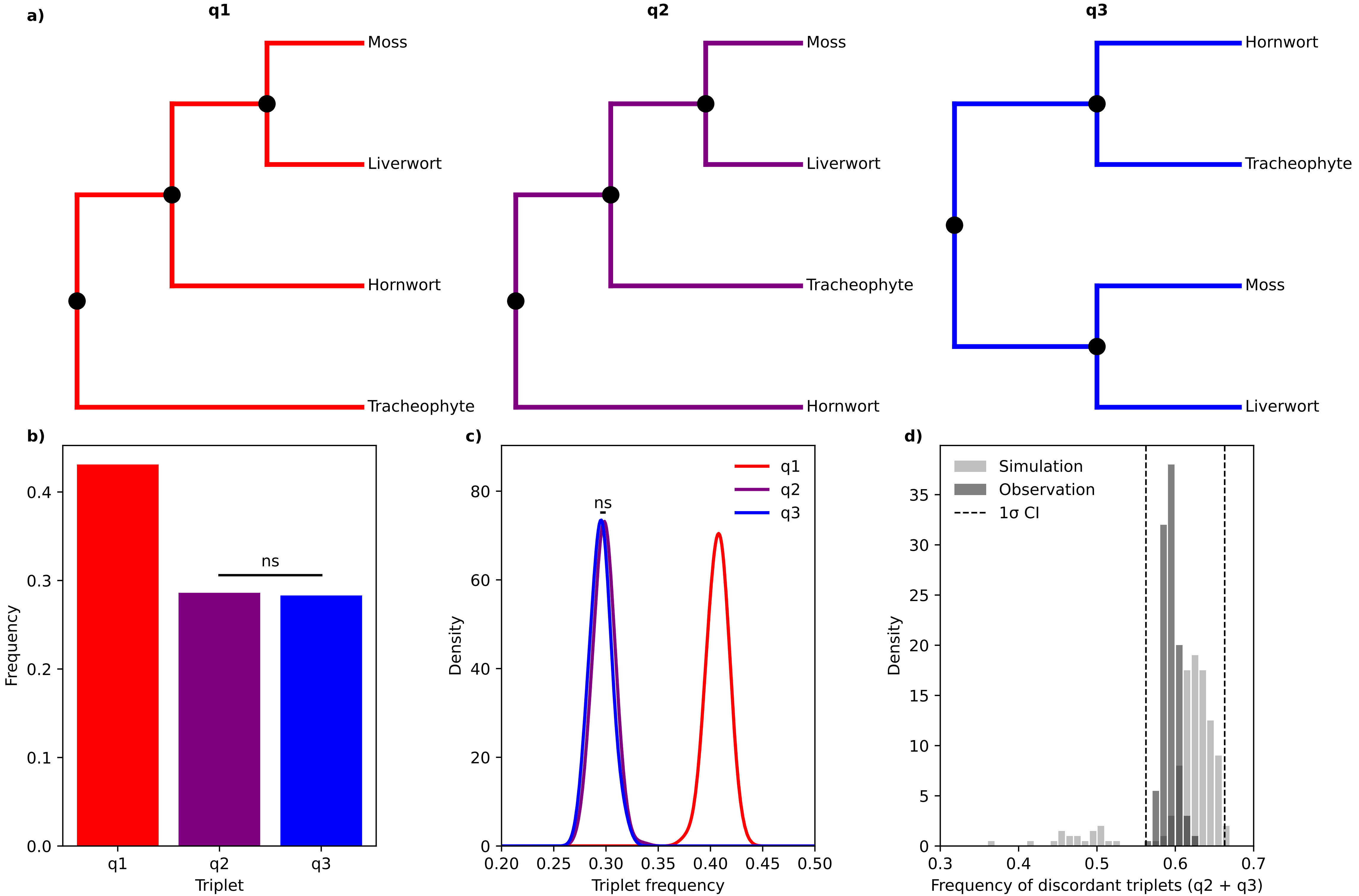

### Extended Data Figure 2

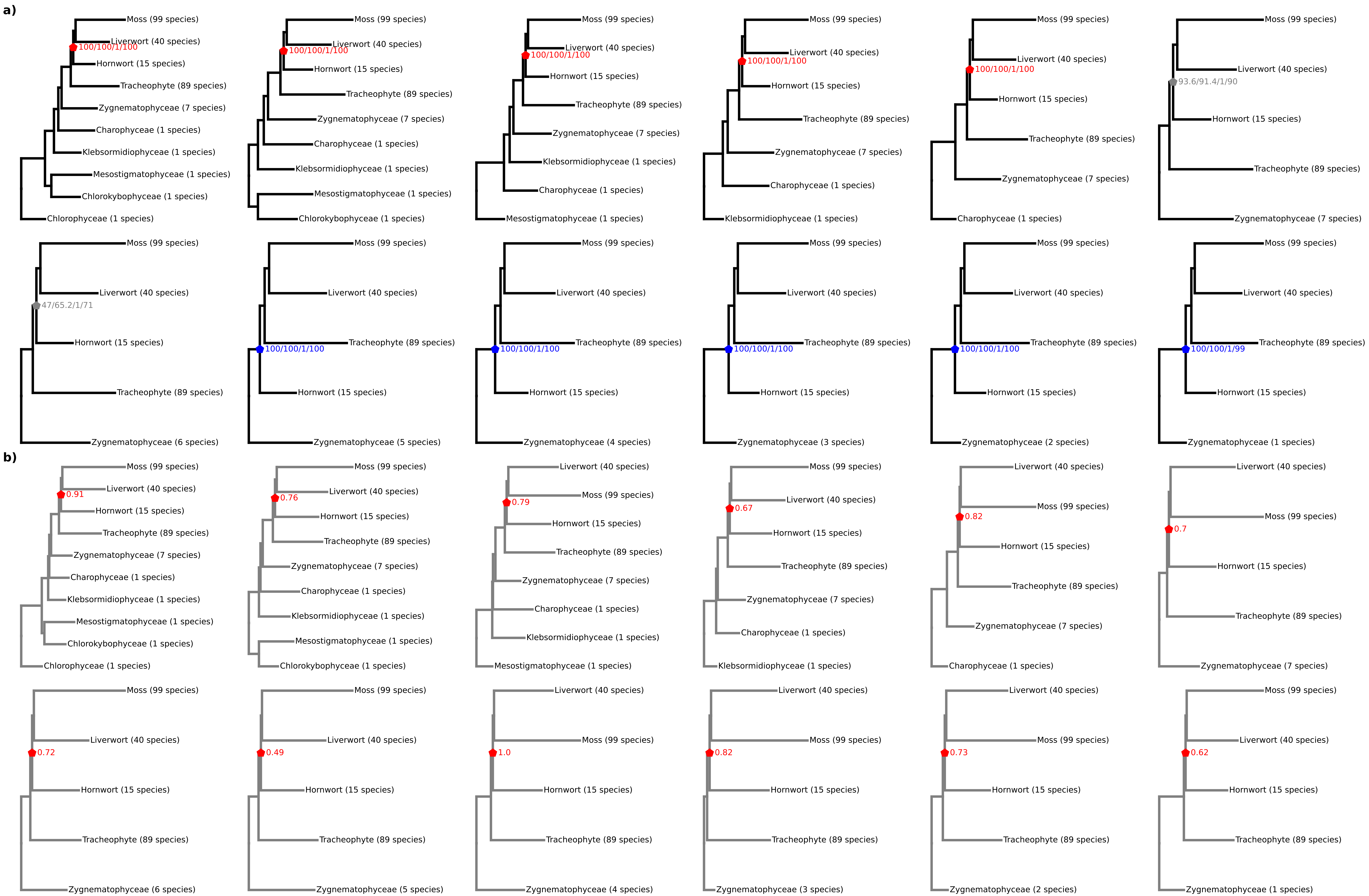

### Extended Data Figure 3

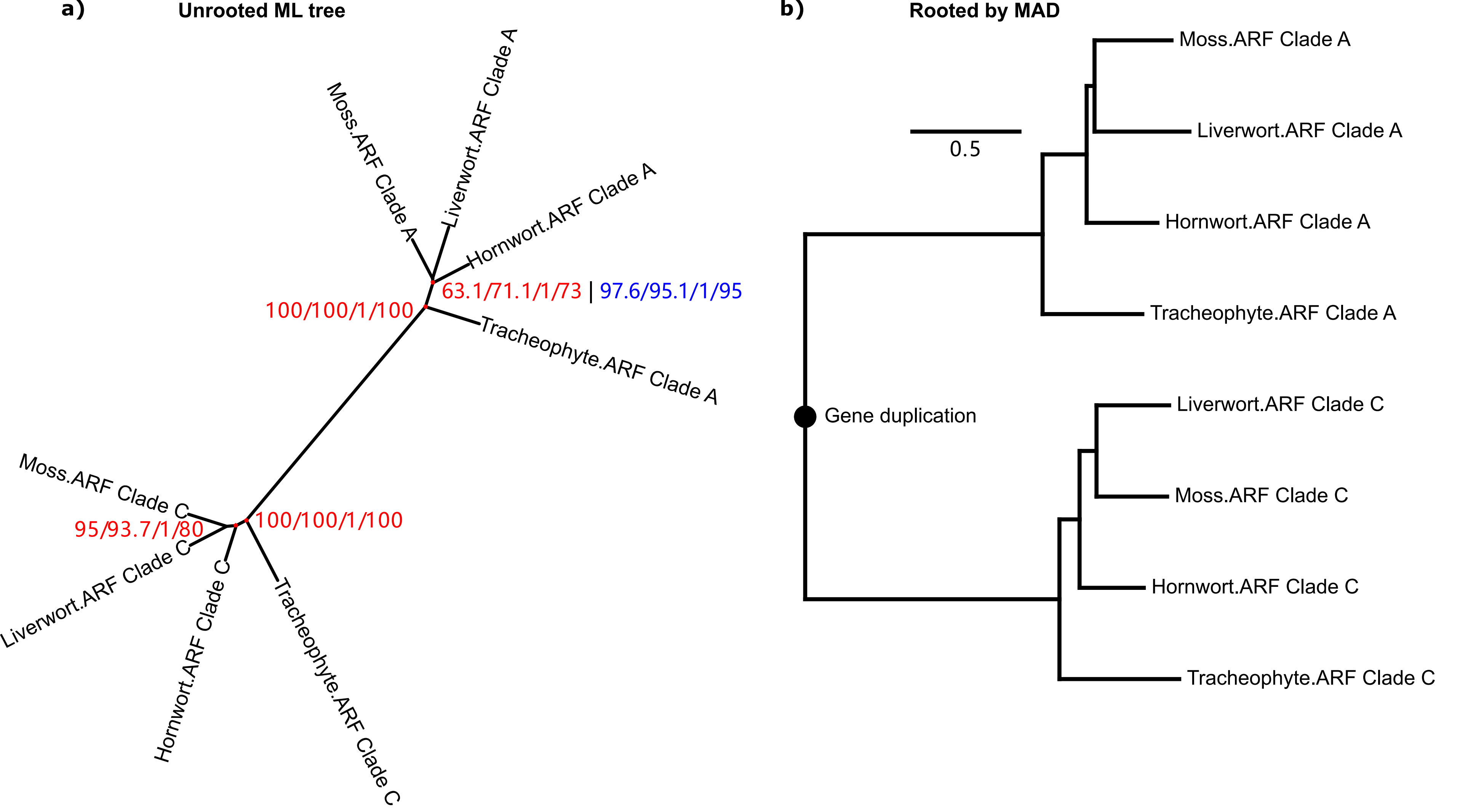
