## Supplementary Figures S1 to S9 for "Rooting the deep divergence of land plants"

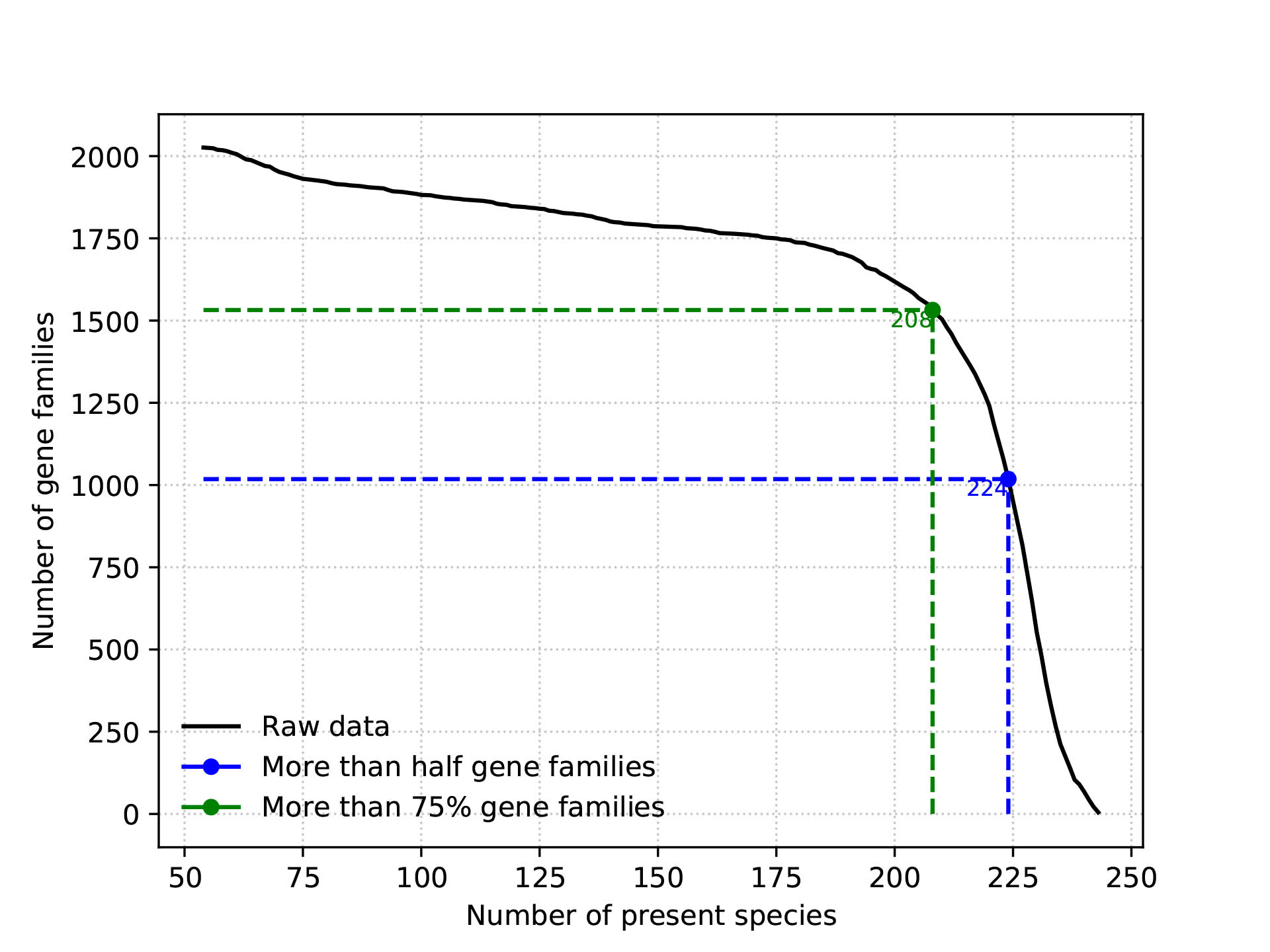

**Figure S1. The relationship between the number of present species and the number of gene families in the 2026-RefSOG dataset.**

More than half of the 2,026 RefSOGs have at least 224 species present while more than 75% of the 2,026 RefSOGs have at least 208 species present.

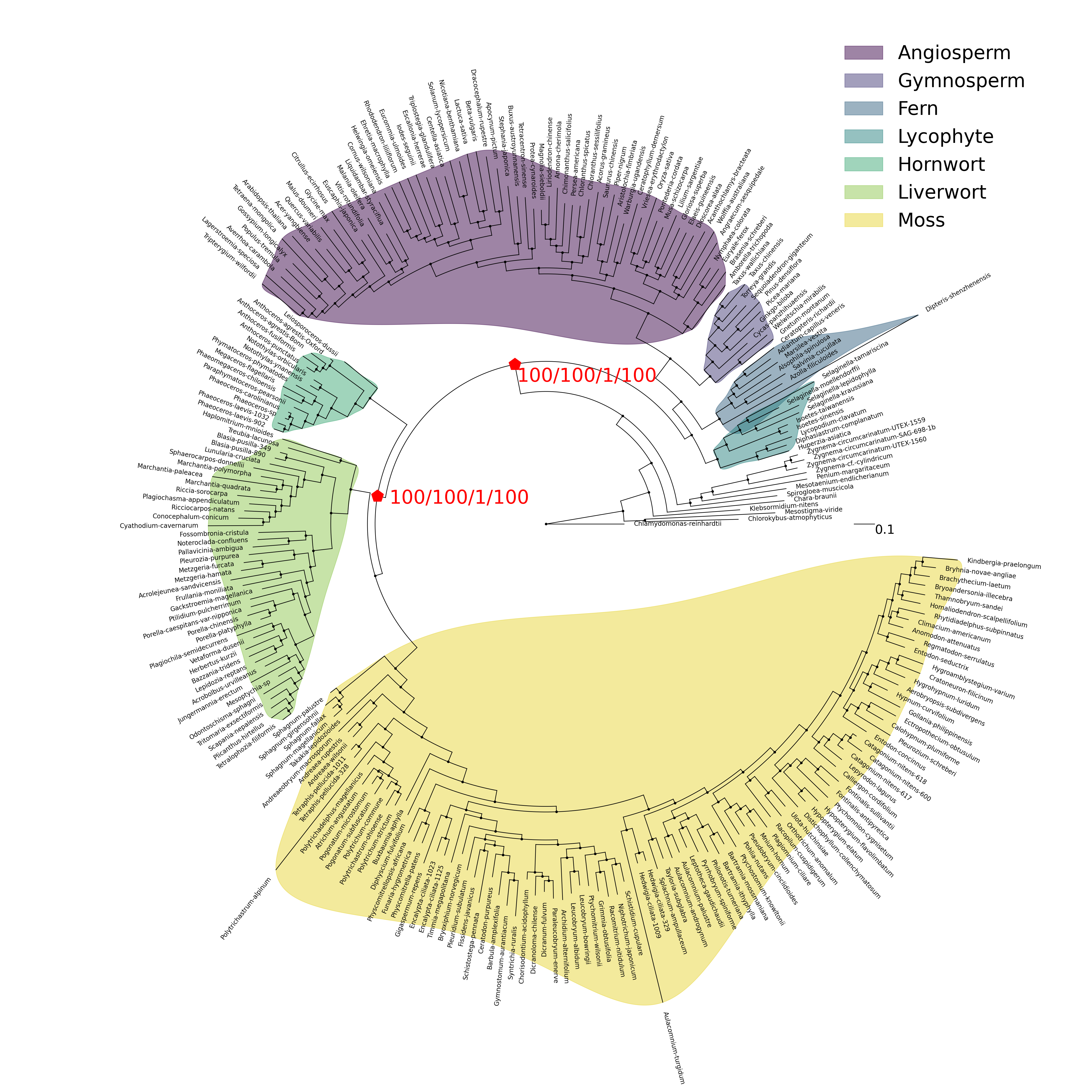

**Figure S2. Species tree inferred by the concatenation method based on the 137-rSOG dataset.**

Major embryophyte clades are highlighted in distinct colors, including angiosperms, gymnosperms, ferns, lycophytes, hornworts, liverworts, and mosses. The supporting level for the crown bryophyte and embryophyte nodes is measured by 1,000 ultrafast bootstrap replicates, SH-like approximate likelihood ratio test, approximate Bayes test, and the local bootstrap probability. The branch length represents the number of substitutions per site.

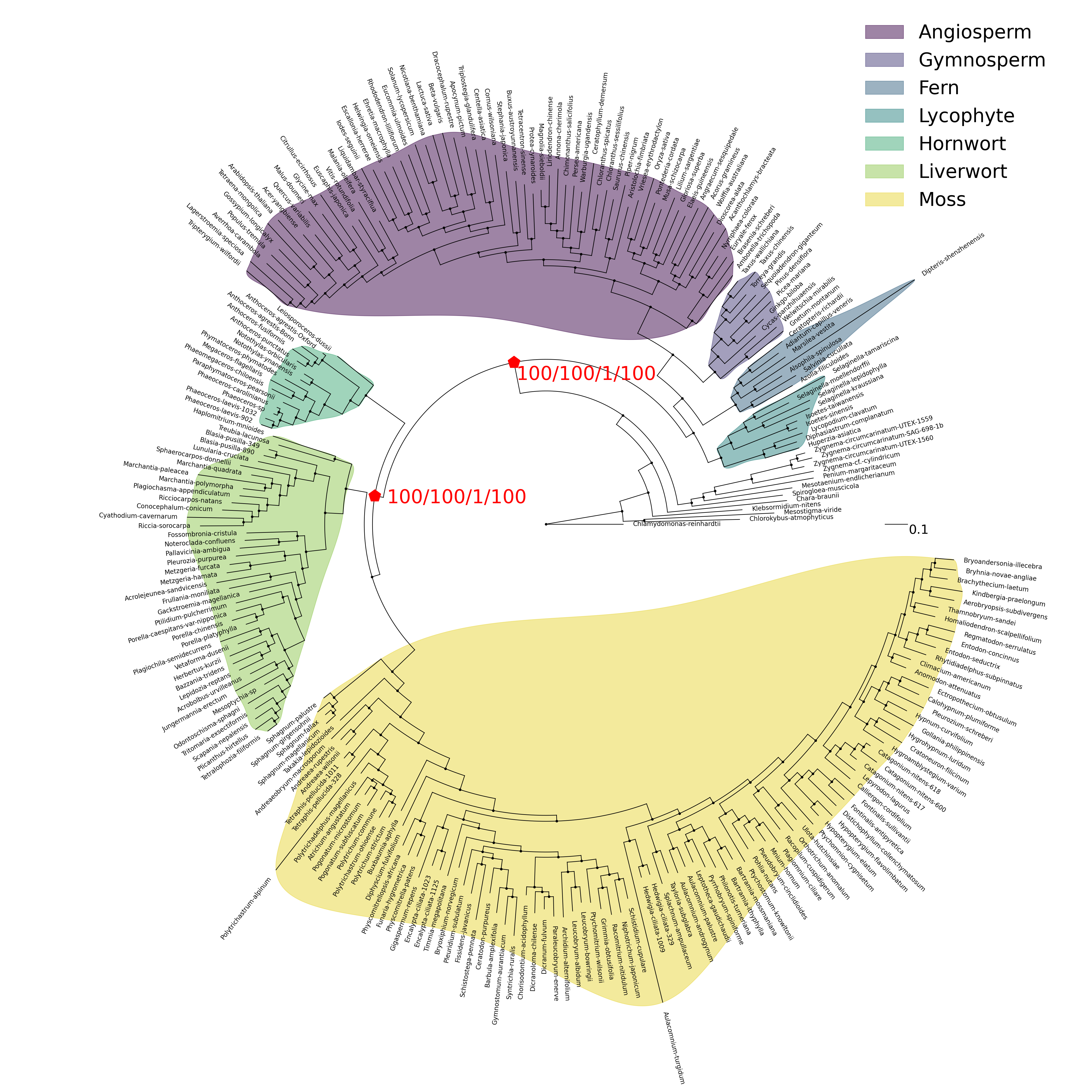

**Figure S3. Species tree inferred by the concatenation method based on the 137-rSOG dataset using the BIC-best model.**

Major embryophyte clades are highlighted in distinct colors, including angiosperms, gymnosperms, ferns, lycophytes, hornworts, liverworts, and mosses. The supporting level for the crown bryophyte and embryophyte nodes is measured by 1,000 ultrafast bootstrap replicates, SH-like approximate likelihood ratio test, approximate Bayes test, and the local bootstrap probability. The branch length represents the number of substitutions per site.

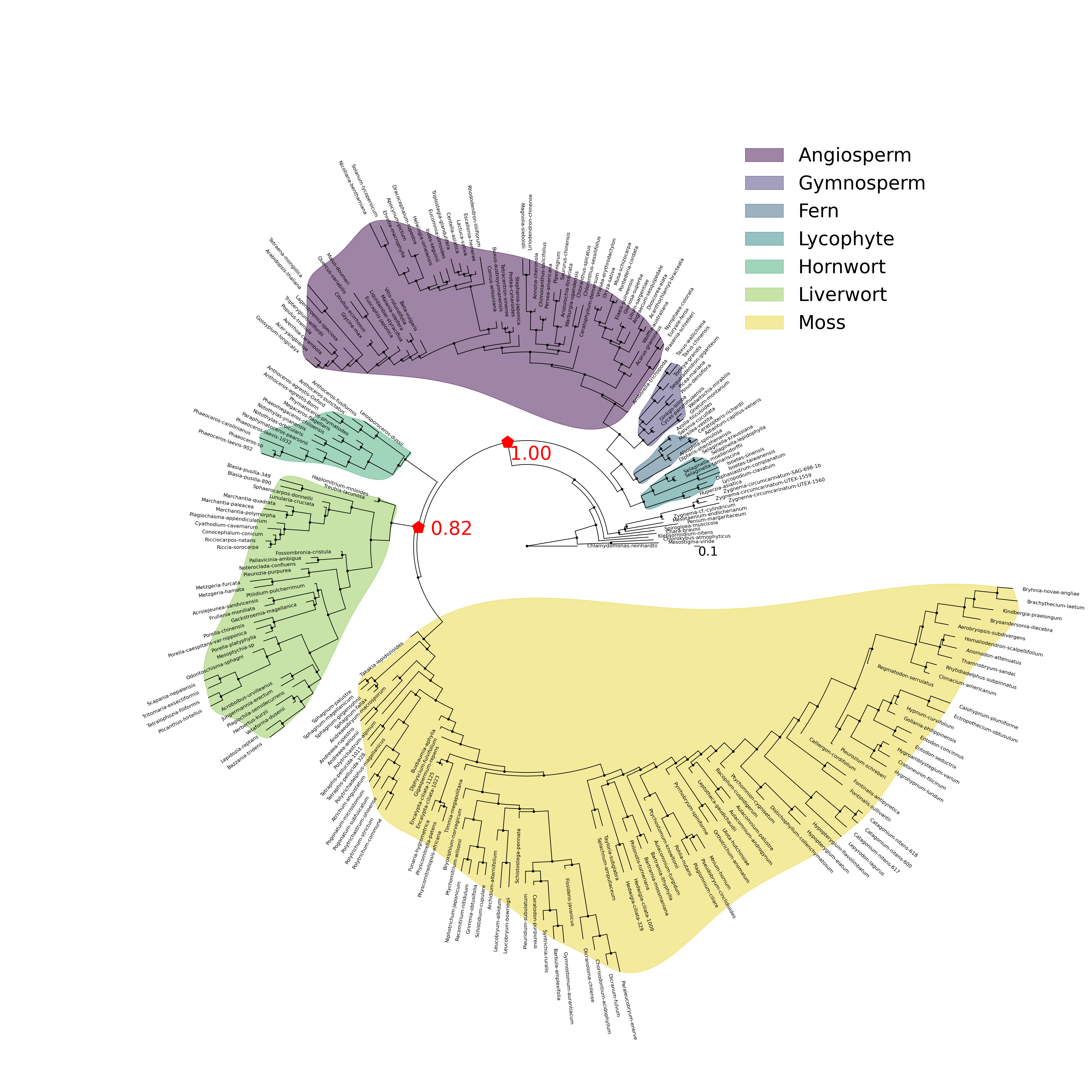

**Figure S4. Species tree inferred by the MSC method based on the 137-rSOG dataset using the BIC-best model.**

Major embryophyte clades are highlighted in distinct colors, including angiosperms, gymnosperms, ferns, lycophytes, hornworts, liverworts, and mosses. The supporting level for the crown node of bryophytes and embryophytes is measured by the local posterior probability. The branch length represents the number of coalescent unit.

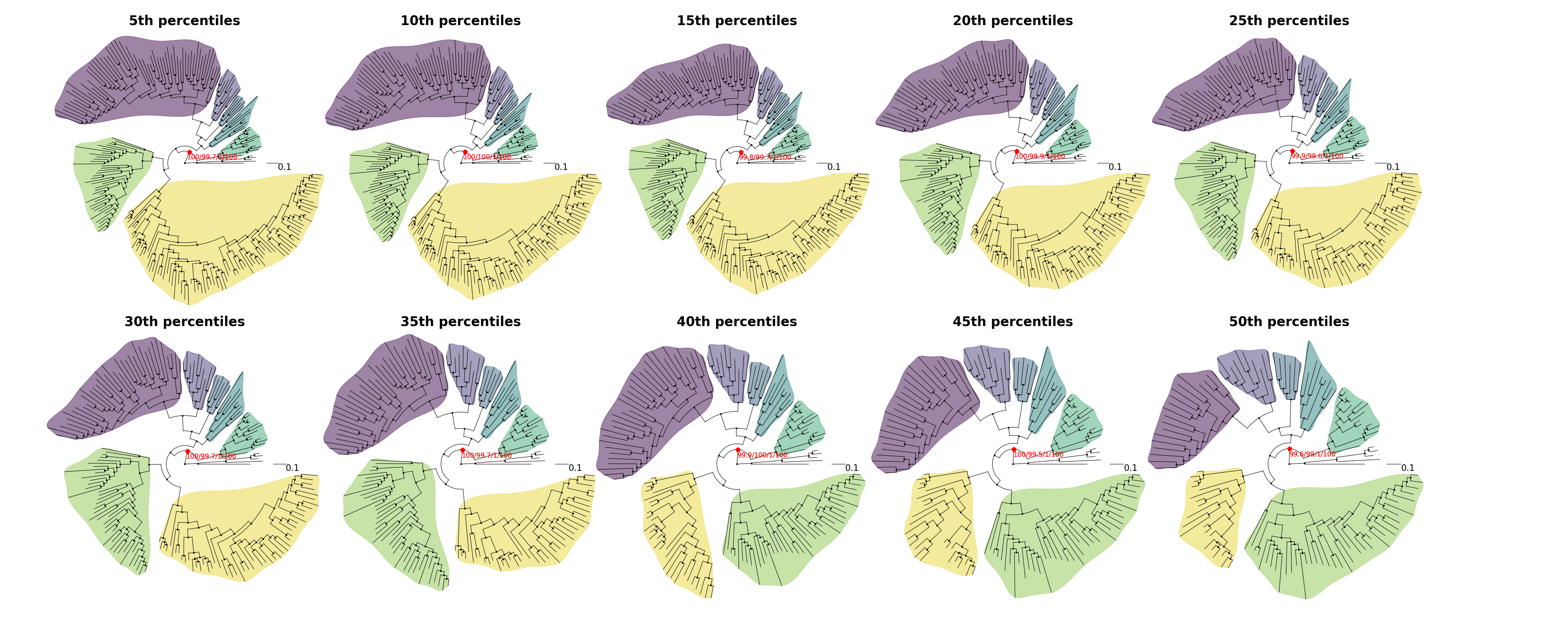

**Figure S5. Species tree inferred by the concatenation method based on the 137-rSOG dataset with *Zygnema* and *Mesotaenium* retained as outgroup representatives while varying the percentage of removed long branches.**

Major embryophyte clades are highlighted in distinct colors with the same color code as in Fig. S2. The supporting level for the crown bryophyte and embryophyte nodes is measured by 1,000 ultrafast bootstrap replicates, SH-like approximate likelihood ratio test, approximate Bayes test, and the local bootstrap probability. The branch length represents the number of substitutions per site. The number of the percentile is denoted atop of each tree.

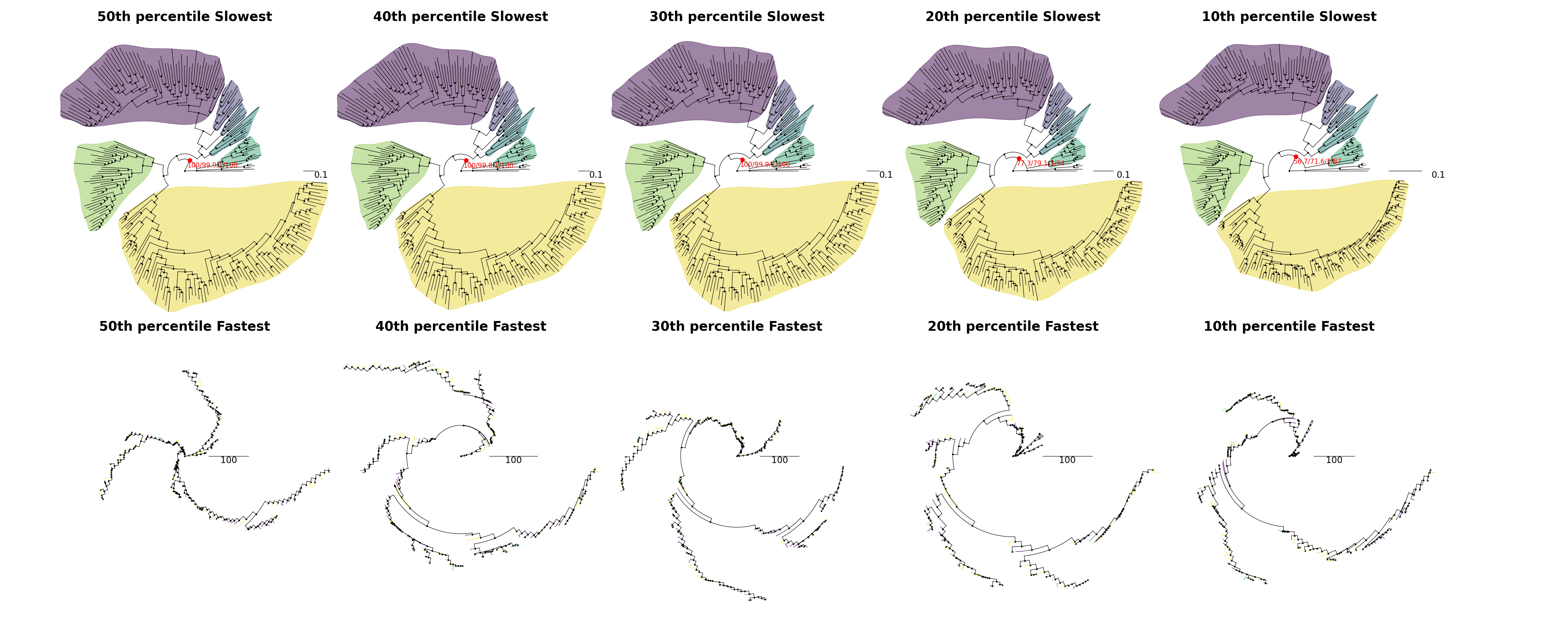

**Figure S6. Species tree inferred by the concatenation method based on the 137-rSOG dataset with *Zygnema* and *Mesotaenium* retained as outgroup representatives while varying the percentage of retained slowest- or fastest-evolving sites.**

Major embryophyte clades are highlighted in distinct colors with the same color code as in Fig. S2 for the datasets consisting of only slowest-evolving sites. Tip branches are colored according to their clade affiliations with the same color code as in Fig. S2 for the datasets consisting of only fastest-evolving sites. The supporting level for the crown bryophyte and embryophyte nodes is measured by 1,000 ultrafast bootstrap replicates, SH-like approximate likelihood ratio test, approximate Bayes test, and the local bootstrap probability. The branch length represents the number of substitutions per site. The number of the percentile is denoted atop of each tree.

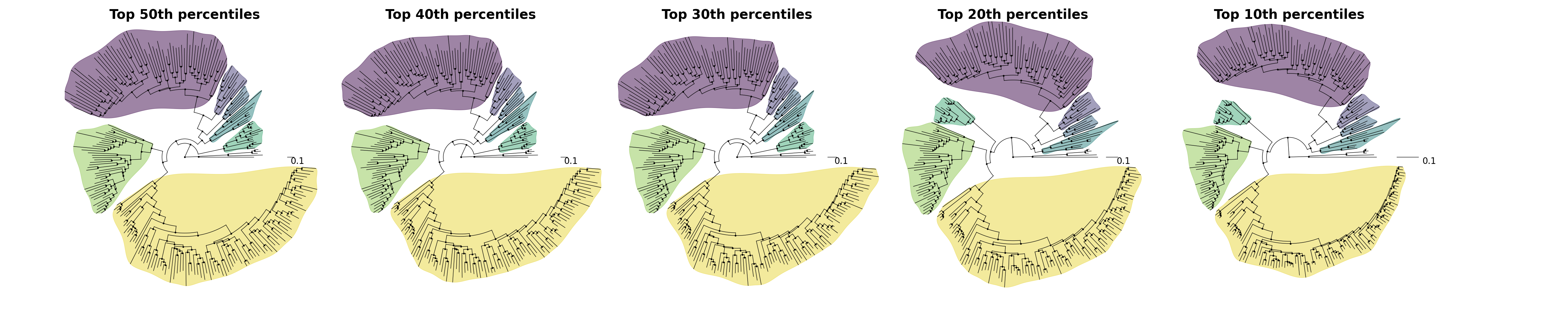

**Figure S7. Species tree inferred by the concatenation method based on the 137-rSOG dataset with *Zygnema* and *Mesotaenium* retained as outgroup representatives while varying the percentage of retained slowest-evolving sites using the LG+C60+F+G+PMSF mixture model.**

Major embryophyte clades are highlighted in distinct colors with the same color code as in Fig. S2. The branch length represents the number of substitutions per site. The number of the percentile is denoted atop of each tree.

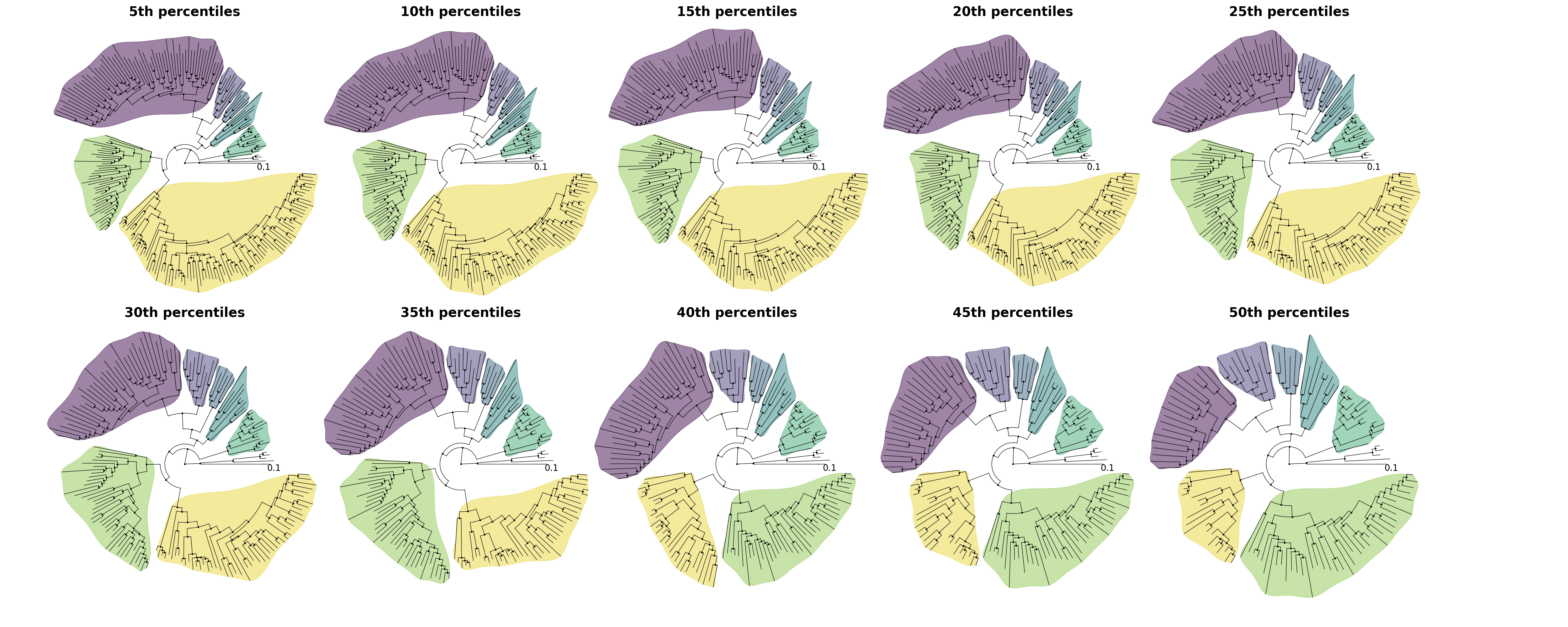

**Figure S8. Species tree inferred by the concatenation method based on the 137-rSOG dataset with *Zygnema* and *Mesotaenium* retained as outgroup representatives while varying the percentage of removed long branches using the LG+C60+F+G+PMSF mixture model.**

Major embryophyte clades are highlighted in distinct colors with the same color code as in Fig. S2. The branch length represents the number of substitutions per site. The number of the percentile is denoted atop of each tree.

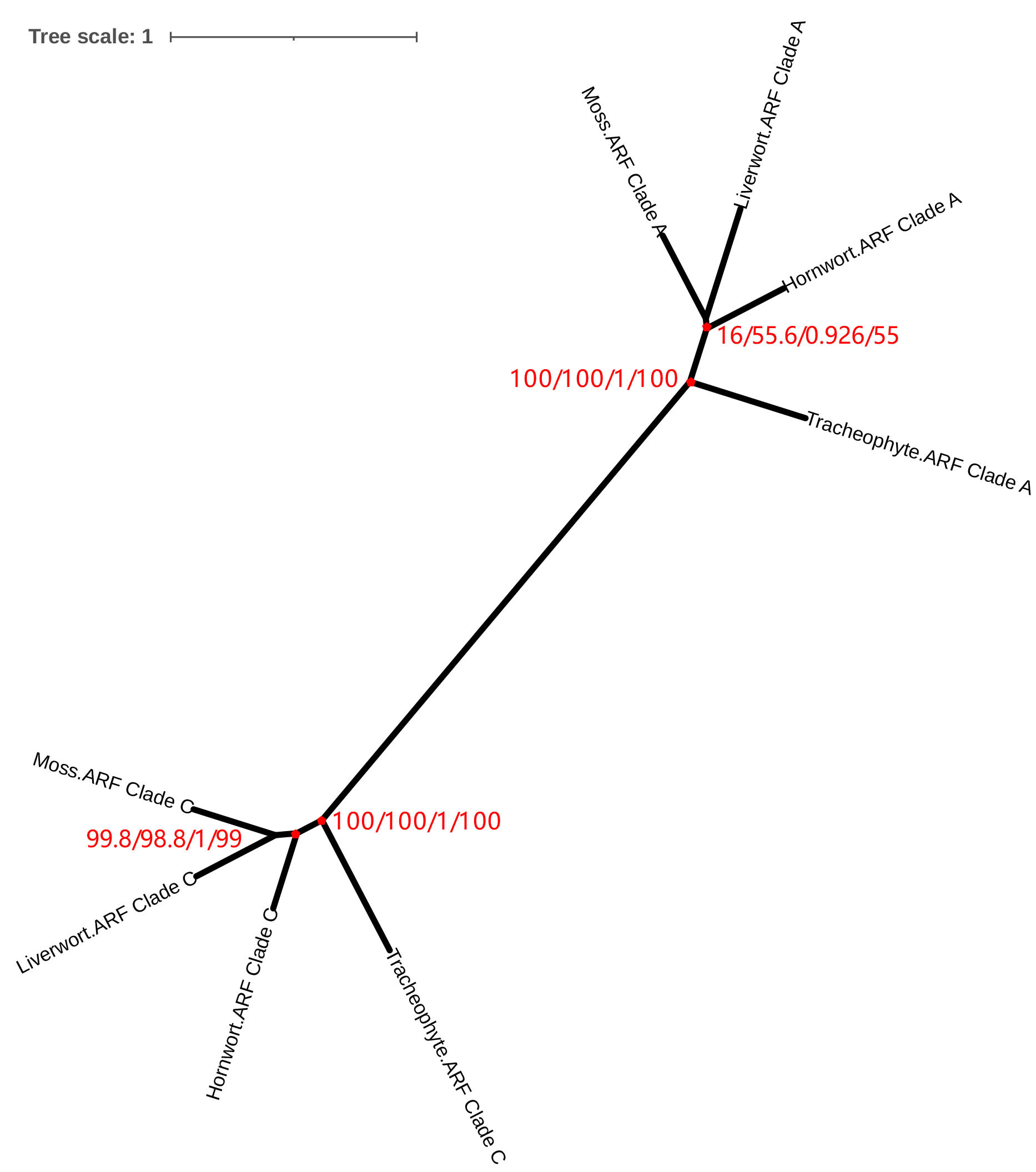

**Figure S9. ML gene tree inferred from the composite alignment consisting of clade-A and clade-C ARFs for the four major embryophyte clades (i.e., hornworts, liverworts, mosses, tracheophytes) using the LG model.**

The supporting level for the crown bryophyte and embryophyte node is measured by 1,000 ultrafast bootstrap replicates, SH-like approximate likelihood ratio test, approximate Bayes test, and the local bootstrap probability. The branch length represents the number of substitutions per site.
