## Supplementary Table S2 for "Rooting the deep divergence of land plants"

**Table S2. Genome data source and taxonomic information**

| **Species** | **Order** | **Class** | **Data source** |
| --- | --- | --- | --- |
| *Chlamydomonas reinhardtii* | Chlamydomonadales | Chlorophyceae | https://phytozome-next.jgi.doe.gov/info/CreinhardtiiCC_4532_v6_1 |
| *Klebsormidium nitens* | Klebsormidiales | Klebsormidiophyceae | https://www.ncbi.nlm.nih.gov/datasets/genome/GCA_000708835.1/ |
| *Chlorokybus atmophyticus* | Chlorokybales | Chlorokybophyceae | CNGB Nucleotide Sequence Archive (accession no. CNP0000228) |
| *Mesostigma viride* | Mesostigmatales | Mesostigmatophyceae | CNGB Nucleotide Sequence Archive (access |
| *Chara braunii* | Charales | Charophyceae | https://www.ncbi.nlm.nih.gov/datasets/genome/GCA_003427395.1/ |
| *Spirogloea muscicola* | Spirogloeales | Zygnematophyceae | https://figshare.com/articles/Genomes_of_subaerial_Zygnematophyceae_provide_insights_into_land_plant_evolution/9911876/1 |
| *Penium margaritaceum* | Desmidiales | Zygnematophyceae | http://bioinfo.bti.cornell.edu/ftp/Penium/ |
| *Mesotaenium endlicherianum* | Zygnematales | Zygnematophyceae | https://figshare.com/articles/Genomes_of_subaerial_Zygnematophyceae_provide_insights_into_land_plant_evolution/9911876/1 |
| *Zygnema cf. cylindricum* | Zygnematales | Zygnematophyceae | https://genome.jgi.doe.gov/portal/pages/dynamicOrganismDownload.jsf?organism=Zygcyl6981a_1 |
| *Z. circumcarinatum* SAG 698-1b | Zygnematales | Zygnematophyceae | https://genome.jgi.doe.gov/portal/pages/dynamicOrganismDownload.jsf?organism=Zygcir6981b_2 |
| *Z. circumcarinatum* UTEX 1559 | Zygnematales | Zygnematophyceae | https://phycocosm.jgi.doe.gov/Zygcir1559_1/Zygcir1559_1.home.html |
| *Z. circumcarinatum* UTEX 1560 | Zygnematales | Zygnematophyceae | https://genome.jgi.doe.gov/portal/pages/dynamicOrganismDownload.jsf?organism=Zygcir1560_1 |
| *Anthoceros agrestis* [Bonn] | Anthocerotales | Anthocerotopsida | https://www.hornworts.uzh.ch/en/download.html |
| *Anthoceros agrestis* [Oxford] | Anthocerotales | Anthocerotopsida | https://hornwortbase.org/ftp/Anthoceros_agrestis_Oxford/v1.0/ |
| *Anthoceros fusiformis* | Anthocerotales | Anthocerotopsida | https://hornwortbase.org/ftp/Anthoceros_fusiformis/v1.0/ |
| *Anthoceros punctatus* | Anthocerotales | Anthocerotopsida | https://hornwortbase.org/ftp/Anthoceros_punctatus/v1.0/ |
| *Notothylas yunnanensis* | Notothyladales | Anthocerotopsida | https://ftp.cngb.org/pub/CNSA/data2/CNP0002895/CNS0548206/CNA0052665/ |
| *Notothylas orbicularis* | Notothyladales | Anthocerotopsida | https://hornwortbase.org/ftp/Notothylas_orbicularis/v1.0/ |
| *Paraphymatoceros pearsonii* | Notothyladales | Anthocerotopsida | https://hornwortbase.org/ftp/Paraphymatoceros_pearsonii/v1.0/ |
| *Phaeoceros carolinianus* | Notothyladales | Anthocerotopsida | https://hornwortbase.org/ftp/Phaeoceros_carolinianus/v1.0/ |
| *Phaeoceros laevis* 902 | Notothyladales | Anthocerotopsida | https://ftp.cngb.org/pub/CNSA/data2/CNP0002895/CNS0548208/CNA0052666/ |
| *Phaeoceros laevis* 1032 | Notothyladales | Anthocerotopsida | https://ftp.cngb.org/pub/CNSA/data2/CNP0002895/CNS0538617/CNA0052667/ |
| *Phaeoceros* sp. | Notothyladales | Anthocerotopsida | https://hornwortbase.org/ftp/Phaeoceros_sp/v1.0/ |
| *Megaceros flagellaris* | Dendrocerotales | Anthocerotopsida | https://hornwortbase.org/ftp/Megaceros_flagellaris/v1.0/ |
| *Phaeomegaceros chiloensis* | Dendrocerotales | Anthocerotopsida | https://hornwortbase.org/ftp/Phaeomegaceros_chiloensis/v1.0/ |
| *Phymatoceros phymatodes* | Phymatocerotales | Anthocerotopsida | https://hornwortbase.org/ftp/Phymatoceros_phymatodes/v1.0/ |
| *Leiosporoceros dussii* | Leiosporocerotales | Leiosporocerotopsida | https://hornwortbase.org/ftp/Leiosporoceros_dussii/v1.0/ |
| *Haplomitrium mnioides* | Calobryales | Haplomitriopsida | https://ftp.cngb.org/pub/CNSA/data2/CNP0002895/CNS0544944/CNA0052668/ |
| *Treubia lacunosa* | Treubiales | Haplomitriopsida | https://ftp.cngb.org/pub/CNSA/data2/CNP0002895/CNS0539616/CNA0052669/ |
| *Blasia pusilla* 349 | Blasiales | Marchantiopsida | https://ftp.cngb.org/pub/CNSA/data2/CNP0002895/CNS0539593/CNA0052670/ |
| *Blasia pusilla* 890 | Blasiales | Marchantiopsida | https://ftp.cngb.org/pub/CNSA/data2/CNP0002895/CNS0538584/CNA0052671/ |
| *Sphaerocarpos donnellii* | Sphaerocarpales | Marchantiopsida | https://ftp.cngb.org/pub/CNSA/data2/CNP0002895/CNS0544948/CNA0052672/ |
| *Lunularia cruciata* | Lunulariales | Marchantiopsida | https://ftp.cngb.org/pub/CNSA/data2/CNP0002895/CNS0539612/CNA0052673/ |
| *Plagiochasma appendiculatum* | Marchantiales | Marchantiopsida | https://ftp.cngb.org/pub/CNSA/data2/CNP0002895/CNS0538592/CNA0052674/ |
| *Conocephalum conicum* | Marchantiales | Marchantiopsida | https://ftp.cngb.org/pub/CNSA/data2/CNP0002895/CNS0538575/CNA0052675/ |
| *Cyathodium cavernarum* | Marchantiales | Marchantiopsida | https://ftp.cngb.org/pub/CNSA/data2/CNP0002895/CNS0538588/CNA0052676/ |
| *Ricciocarpos natans* | Marchantiales | Marchantiopsida | https://ftp.cngb.org/pub/CNSA/data2/CNP0002895/CNS0548193/CNA0052677/ |
| *Marchantia polymorpha* | Marchantiales | Marchantiopsida | https://phytozome-next.jgi.doe.gov/info/Mpolymorpha_v3_1 |
| *Marchantia paleacea* | Marchantiales | Marchantiopsida | https://ftp.ncbi.nlm.nih.gov/genomes/all/GCA/014/180/765/GCA_014180765.2_ASM1418076v2/ |
| *Marchantia quadrata* | Marchantiales | Marchantiopsida | https://doi.org/10.6084/m9.figshare.27055057 |
| *Riccia sorocarpa* | Marchantiales | Marchantiopsida | https://ftp.ncbi.nlm.nih.gov/genomes/all/GCA/046/195/345/GCA_046195345.1_ASM4619534v1/ |
| *Fossombronia cristula* | Fossombroniales | Jungermanniopsida | https://ftp.cngb.org/pub/CNSA/data2/CNP0002895/CNS0539613/CNA0052678/ |
| *Noteroclada confluens* | Pelliales | Jungermanniopsida | https://ftp.cngb.org/pub/CNSA/data2/CNP0002895/CNS0539607/CNA0052679/ |
| *Pallavicinia ambigua* | Pallaviciniales | Jungermanniopsida | https://ftp.cngb.org/pub/CNSA/data2/CNP0002895/CNS0538591/CNA0052680/ |
| *Pleurozia purpurea* | Pleuroziales | Jungermanniopsida | https://ftp.cngb.org/pub/CNSA/data2/CNP0002895/CNS0538585/CNA0052681/ |
| *Metzgeria furcata* | Metzgeriales | Jungermanniopsida | https://ftp.cngb.org/pub/CNSA/data2/CNP0002895/CNS0548198/CNA0052682/ |
| *Metzgeria hamata* | Metzgeriales | Jungermanniopsida | https://ftp.cngb.org/pub/CNSA/data2/CNP0002895/CNS0548192/CNA0052683/ |
| *Ptilidium pulcherrimum* | Ptilidiales | Jungermanniopsida | https://ftp.cngb.org/pub/CNSA/data2/CNP0002895/CNS0539611/CNA0052684/ |
| *Gackstroemia magellanica* | Porellales | Jungermanniopsida | https://ftp.cngb.org/pub/CNSA/data2/CNP0002895/CNS0539597/CNA0052685/ |
| *Acrolejeunea sandvicensis* | Porellales | Jungermanniopsida | https://ftp.cngb.org/pub/CNSA/data2/CNP0002895/CNS0538583/CNA0052686/ |
| *Frullania moniliata* | Porellales | Jungermanniopsida | https://ftp.cngb.org/pub/CNSA/data2/CNP0002895/CNS0538603/CNA0052687/ |
| *Porella chinensis* | Porellales | Jungermanniopsida | https://ftp.cngb.org/pub/CNSA/data2/CNP0002895/CNS0538614/CNA0052688/ |
| *Porella platyphylla* | Porellales | Jungermanniopsida | https://ftp.cngb.org/pub/CNSA/data2/CNP0002895/CNS0539610/CNA0052689/ |
| *Porella caespitans var. nipponica* | Porellales | Jungermanniopsida | https://ftp.cngb.org/pub/CNSA/data2/CNP0002895/CNS0538610/CNA0052690/ |
| *Mesoptychia sp.* | Jungermanniales | Jungermanniopsida | https://ftp.cngb.org/pub/CNSA/data2/CNP0002895/CNS0548202/CNA0052691/ |
| *Odontoschisma sphagni* | Jungermanniales | Jungermanniopsida | https://ftp.cngb.org/pub/CNSA/data2/CNP0002895/CNS0539591/CNA0052692/ |
| *Scapania nepalensis* | Jungermanniales | Jungermanniopsida | https://ftp.cngb.org/pub/CNSA/data2/CNP0002895/CNS0548197/CNA0052693/ |
| *Tritomaria exsectiformis* | Jungermanniales | Jungermanniopsida | https://ftp.cngb.org/pub/CNSA/data2/CNP0002895/CNS0538609/CNA0052694/ |
| *Plicanthus hirtellus* | Jungermanniales | Jungermanniopsida | https://ftp.cngb.org/pub/CNSA/data2/CNP0002895/CNS0538607/CNA0052695/ |
| *Tetralophozia filiformis* | Jungermanniales | Jungermanniopsida | https://ftp.cngb.org/pub/CNSA/data2/CNP0002895/CNS0538608/CNA0052696/ |
| *Acrobolbus urvilleanus* | Jungermanniales | Jungermanniopsida | https://ftp.cngb.org/pub/CNSA/data2/CNP0002895/CNS0539608/CNA0052697/ |
| *Jungermannia erectum* | Jungermanniales | Jungermanniopsida | https://ftp.cngb.org/pub/CNSA/data2/CNP0002895/CNS0548194/CNA0052698/ |
| *Plagiochila semidecurrens* | Jungermanniales | Jungermanniopsida | https://ftp.cngb.org/pub/CNSA/data2/CNP0002895/CNS0539594/CNA0052699/ |
| *Lepidozia reptans* | Jungermanniales | Jungermanniopsida | https://ftp.cngb.org/pub/CNSA/data2/CNP0002895/CNS0538606/CNA0052701/ |
| *Bazzania tridens* | Jungermanniales | Jungermanniopsida | https://ftp.cngb.org/pub/CNSA/data2/CNP0002895/CNS0538615/CNA0052702/ |
| *Vetaforma dusenii* | Jungermanniales | Jungermanniopsida | https://ftp.cngb.org/pub/CNSA/data2/CNP0002895/CNS0539606/CNA0052703/ |
| *Herbertus kurzii* | Jungermanniales | Jungermanniopsida | https://ftp.cngb.org/pub/CNSA/data2/CNP0002895/CNS0538599/CNA0052704/ |
| *Sphagnum girgensohnii* | Sphagnales | Sphagnopsida | https://ftp.cngb.org/pub/CNSA/data2/CNP0002895/CNS0538587/CNA0052705/ |
| *Sphagnum palustre* | Sphagnales | Sphagnopsida | https://ftp.cngb.org/pub/CNSA/data2/CNP0002895/CNS0538598/CNA0052706/ |
| *Sphagnum magellanicum* | Sphagnales | Sphagnopsida | https://phytozome-next.jgi.doe.gov/info/Smagellanicum_v1_1 |
| *Sphagnum fallax* | Sphagnales | Sphagnopsida | https://phytozome-next.jgi.doe.gov/info/Sfallax_v1_1 |
| *Andreaeobryum macrosporum* | Andreaeobryales | Andreaeobryopsida | https://ftp.cngb.org/pub/CNSA/data2/CNP0002895/CNS0544945/CNA0052707/ |
| *Andreaea rupestris* | Andreaeales | Andreaeopsida | https://ftp.cngb.org/pub/CNSA/data2/CNP0002895/CNS0538573/CNA0052708/ |
| *Andreaea wilsonii* | Andreaeales | Andreaeopsida | https://ftp.cngb.org/pub/CNSA/data2/CNP0002895/CNS0539614/CNA0052709/ |
| *Tetraphis pellucida* 328 | Tetraphidales | Tetraphidopsida | https://ftp.cngb.org/pub/CNSA/data2/CNP0002895/CNS0539582/CNA0052710/ |
| *Tetraphis pellucida* 1011 | Tetraphidales | Tetraphidopsida | https://ftp.cngb.org/pub/CNSA/data2/CNP0002895/CNS0539570/CNA0052711/ |
| *Polytrichadelphus magellanicus* | Polytrichales | Polytrichopsida | https://ftp.cngb.org/pub/CNSA/data2/CNP0002895/CNS0539602/CNA0052712/ |
| *Atrichum angustatum* | Polytrichales | Polytrichopsida | https://ftp.cngb.org/pub/CNSA/data2/CNP0002895/CNS0539571/CNA0052713/ |
| *Pogonatum microstomum* | Polytrichales | Polytrichopsida | https://ftp.cngb.org/pub/CNSA/data2/CNP0002895/CNS0538602/CNA0052714/ |
| *Pogonatum subfuscatum* | Polytrichales | Polytrichopsida | https://ftp.cngb.org/pub/CNSA/data2/CNP0002895/CNS0538619/CNA0052715/ |
| *Polytrichum strictum* | Polytrichales | Polytrichopsida | https://ftp.cngb.org/pub/CNSA/data2/CNP0002895/CNS0539599/CNA0052716/ |
| *Polytrichum commune* | Polytrichales | Polytrichopsida | https://ftp.cngb.org/pub/CNSA/data2/CNP0002895/CNS0538618/CNA0052718/ |
| *Polytrichastrum ohioense* | Polytrichales | Polytrichopsida | https://ftp.cngb.org/pub/CNSA/data2/CNP0002895/CNS0539617/CNA0052717/ |
| *Polytrichastrum alpinum* | Polytrichales | Polytrichopsida | https://doi.org/10.6084/m9.figshare.28595150 |
| *Buxbaumia aphylla* | Buxbaumiales | Bryopsida | https://ftp.cngb.org/pub/CNSA/data2/CNP0002895/CNS0538570/CNA0052719/ |
| *Diphyscium fulvifolium* | Diphysciales | Bryopsida | https://ftp.cngb.org/pub/CNSA/data2/CNP0002895/CNS0538589/CNA0052720/ |
| *Gigaspermum repens* | Gigaspermales | Bryopsida | https://ftp.cngb.org/pub/CNSA/data2/CNP0002895/CNS0539620/CNA0052721/ |
| *Encalypta ciliata* 1023 | Encalyptales | Bryopsida | https://ftp.cngb.org/pub/CNSA/data2/CNP0002895/CNS0538612/CNA0052722/ |
| *Encalypta ciliata* 1125 | Encalyptales | Bryopsida | https://ftp.cngb.org/pub/CNSA/data2/CNP0002895/CNS0548207/CNA0052723/ |
| *Ceratodon purpureus* | Pseudoditrichales | Bryopsida | https://phytozome-next.jgi.doe.gov/info/CpurpureusGG1_v1_1 |
| *Timmia megapolitana* | Timmiales | Bryopsida | https://ftp.cngb.org/pub/CNSA/data2/CNP0002895/CNS0539569/CNA0052724/ |
| *Bryoxiphium norvegicum* | Bryoxiphiales | Bryopsida | https://ftp.cngb.org/pub/CNSA/data2/CNP0002895/CNS0538590/CNA0052725/ |
| *Ptychomitrium wilsonii* | Grimmiales | Bryopsida | https://ftp.cngb.org/pub/CNSA/data2/CNP0002895/CNS0538578/CNA0052726/ |
| *Racomitrium nitidulum* | Grimmiales | Bryopsida | https://ftp.cngb.org/pub/CNSA/data2/CNP0002895/CNS0548195/CNA0052727/ |
| *Schistidium cupulare* | Grimmiales | Bryopsida | https://ftp.cngb.org/pub/CNSA/data2/CNP0002895/CNS0548199/CNA0052728/ |
| *Grimmia obtusifolia* | Grimmiales | Bryopsida | https://ftp.cngb.org/pub/CNSA/data2/CNP0002895/CNS0539573/CNA0052729/ |
| *Niphotrichum japonicum* | Grimmiales | Bryopsida | https://figshare.com/articles/dataset/_em_strong_Niphotrichum_japonicum_strong_em_strong_genome_project_strong_/23573514 |
| *Archidium alternifolium* | Archidiales | Bryopsida | https://ftp.cngb.org/pub/CNSA/data2/CNP0002895/CNS0539575/CNA0052730/ |
| *Leucobryum albidum* | Dicranales | Bryopsida | https://ftp.cngb.org/pub/CNSA/data2/CNP0002895/CNS0539578/CNA0052731/ |
| *Leucobryum bowringii* | Dicranales | Bryopsida | https://ftp.cngb.org/pub/CNSA/data2/CNP0002895/CNS0538579/CNA0052732/ |
| *Fissidens javanicus* | Dicranales | Bryopsida | https://ftp.cngb.org/pub/CNSA/data2/CNP0002895/CNS0538586/CNA0052733/ |
| *Dicranoloma chilense* | Dicranales | Bryopsida | https://ftp.cngb.org/pub/CNSA/data2/CNP0002895/CNS0539600/CNA0052734/ |
| *Dicranum fulvum* | Dicranales | Bryopsida | https://ftp.cngb.org/pub/CNSA/data2/CNP0002895/CNS0548196/CNA0052735/ |
| *Paraleucobryum enerve* | Dicranales | Bryopsida | https://ftp.cngb.org/pub/CNSA/data2/CNP0002895/CNS0539574/CNA0052736/ |
| *Chorisodontium acidophyllum* | Dicranales | Bryopsida | https://ftp.cngb.org/pub/CNSA/data2/CNP0002895/CNS0548200/CNA0052737/ |
| *Schistostega pennata* | Dicranales | Bryopsida | https://ftp.cngb.org/pub/CNSA/data2/CNP0002895/CNS0544949/CNA0052738/ |
| *Pleuridium subulatum* | Pseudoditrichales | Bryopsida | https://ftp.cngb.org/pub/CNSA/data2/CNP0002895/CNS0539615/CNA0052739/ |
| *Barbula amplexifolia* | Pottiales | Bryopsida | https://ftp.cngb.org/pub/CNSA/data2/CNP0002895/CNS0538595/CNA0052740/ |
| *Gymnostomum aurantiacum* | Pottiales | Bryopsida | https://ftp.cngb.org/pub/CNSA/data2/CNP0002895/CNS0548204/CNA0052741/ |
| *Syntrichia ruralis* | Pottiales | Bryopsida | https://ftp.cngb.org/pub/CNSA/data2/CNP0002895/CNS0539572/CNA0052742/ |
| *Hedwigia ciliata* 329 | Hedwigiales | Bryopsida | https://ftp.cngb.org/pub/CNSA/data2/CNP0002895/CNS0539583/CNA0052743/ |
| *Hedwigia ciliata* 1009 | Hedwigiales | Bryopsida | https://ftp.cngb.org/pub/CNSA/data2/CNP0002895/CNS0539568/CNA0052744/ |
| *Splachnum ampullaceum* | Splachnales | Bryopsida | https://ftp.cngb.org/pub/CNSA/data2/CNP0002895/CNS0544947/CNA0052745/ |
| *Tayloria subglabra* | Splachnales | Bryopsida | https://ftp.cngb.org/pub/CNSA/data2/CNP0002895/CNS0539619/CNA0052746/ |
| *Philonotis turneriana* | Bartramiales | Bryopsida | https://ftp.cngb.org/pub/CNSA/data2/CNP0002895/CNS0548189/CNA0052747/ |
| *Bartramia mossmaniana* | Bartramiales | Bryopsida | https://ftp.cngb.org/pub/CNSA/data2/CNP0002895/CNS0539601/CNA0052748/ |
| *Bartramia ithyphylla* | Bartramiales | Bryopsida | https://ftp.cngb.org/pub/CNSA/data2/CNP0002895/CNS0538616/CNA0052749/ |
| *Plagiomnium ciliare* | Bryales | Bryopsida | https://ftp.cngb.org/pub/CNSA/data2/CNP0002895/CNS0539589/CNA0052750/ |
| *Pseudobryum cinclidioides* | Bryales | Bryopsida | https://ftp.cngb.org/pub/CNSA/data2/CNP0002895/CNS0539580/CNA0052751/ |
| *Mnium hornum* | Bryales | Bryopsida | https://ftp.cngb.org/pub/CNSA/data2/CNP0002895/CNS0539587/CNA0052752/ |
| *Ptychostomum knowltonii* | Bryales | Bryopsida | https://doi.org/10.6084/m9.figshare.27186099 |
| *Pohlia nutans* | Bryales | Bryopsida | https://ngdc.cncb.ac.cn/gwh/Assembly/24396/show |
| *Pyrrhobryum spiniforme* | Rhizogoniales | Bryopsida | https://ftp.cngb.org/pub/CNSA/data2/CNP0002895/CNS0538576/CNA0052753/ |
| *Leptotheca gaudichaudii* | Rhizogoniales | Bryopsida | https://ftp.cngb.org/pub/CNSA/data2/CNP0002895/CNS0539603/CNA0052754/ |
| *Aulacomnium turgidum* | Rhizogoniales | Bryopsida | https://doi.org/10.6084/m9.figshare.28595150 |
| *Aulacomnium androgynum* | Rhizogoniales | Bryopsida | https://ftp.cngb.org/pub/CNSA/data2/CNP0002895/CNS0544946/CNA0052757/ |
| *Aulacomnium palustre* | Rhizogoniales | Bryopsida | https://ftp.cngb.org/pub/CNSA/data2/CNP0002895/CNS0539579/CNA0052758/ |
| *Ulota hutchinsiae* | Orthotrichales | Bryopsida | https://ftp.cngb.org/pub/CNSA/data2/CNP0002895/CNS0539581/CNA0052755/ |
| *Orthotrichum anomalum* | Orthotrichales | Bryopsida | https://ftp.cngb.org/pub/CNSA/data2/CNP0002895/CNS0538582/CNA0052756/ |
| *Racopilum cuspidigerum* | Hypnodendrales | Bryopsida | https://ftp.cngb.org/pub/CNSA/data2/CNP0002895/CNS0539576/CNA0052759/ |
| *Ptychomnion cygnisetum* | Ptychomniales | Bryopsida | https://ftp.cngb.org/pub/CNSA/data2/CNP0002895/CNS0539598/CNA0052760/ |
| *Distichophyllum collenchymatosum* | Hookeriales | Bryopsida | https://ftp.cngb.org/pub/CNSA/data2/CNP0002895/CNS0538596/CNA0052761/ |
| *Hypopterygium flavolimbatum* | Hookeriales | Bryopsida | https://ftp.cngb.org/pub/CNSA/data2/CNP0002895/CNS0548191/CNA0052762/ |
| *Hypopterygium elatum* | Hookeriales | Bryopsida | https://ftp.cngb.org/pub/CNSA/data2/CNP0002895/CNS0538611/CNA0052763/ |
| *Catagonium nitens* 617 | Hypnales | Bryopsida | https://ftp.cngb.org/pub/CNSA/data2/CNP0002895/CNS0539604/CNA0052764/ |
| *Catagonium nitens* 600 | Hypnales | Bryopsida | https://ftp.cngb.org/pub/CNSA/data2/CNP0002895/CNS0539595/CNA0052765/ |
| *Catagonium nitens* 618 | Hypnales | Bryopsida | https://ftp.cngb.org/pub/CNSA/data2/CNP0002895/CNS0539605/CNA0052766/ |
| *Lepyrodon lagurus* | Hypnales | Bryopsida | https://ftp.cngb.org/pub/CNSA/data2/CNP0002895/CNS0539596/CNA0052767/ |
| *Fontinalis sullivantii* | Hypnales | Bryopsida | https://ftp.cngb.org/pub/CNSA/data2/CNP0002895/CNS0539584/CNA0052768/ |
| *Fontinalis antipyretica* | Hypnales | Bryopsida | http://gigadb.org/dataset/100748 |
| *Climacium americanum* | Hypnales | Bryopsida | https://ftp.cngb.org/pub/CNSA/data2/CNP0002895/CNS0539577/CNA0052769/ |
| *Rhytidiadelphus subpinnatus* | Hypnales | Bryopsida | https://ftp.cngb.org/pub/CNSA/data2/CNP0002895/CNS0539609/CNA0052770/ |
| *Calliergon cordifolium* | Hypnales | Bryopsida | https://ftp.cngb.org/pub/CNSA/data2/CNP0002895/CNS0539618/CNA0052771/ |
| *Aerobryopsis subdivergens* | Hypnales | Bryopsida | https://ftp.cngb.org/pub/CNSA/data2/CNP0002895/CNS0538577/CNA0052772/ |
| *Bryoandersonia illecebra* | Hypnales | Bryopsida | https://ftp.cngb.org/pub/CNSA/data2/CNP0002895/CNS0539586/CNA0052773/ |
| *Kindbergia praelongum* | Hypnales | Bryopsida | https://ftp.cngb.org/pub/CNSA/data2/CNP0002895/CNS0548203/CNA0052774/ |
| *Brachythecium laetum* | Hypnales | Bryopsida | https://ftp.cngb.org/pub/CNSA/data2/CNP0002895/CNS0539585/CNA0052775/ |
| *Bryhnia novae-angliae* | Hypnales | Bryopsida | https://ftp.cngb.org/pub/CNSA/data2/CNP0002895/CNS0539590/CNA0052776/ |
| *Homaliodendron scalpellifolium* | Hypnales | Bryopsida | https://ftp.cngb.org/pub/CNSA/data2/CNP0002895/CNS0538613/CNA0052777/ |
| *Anomodon attenuatus* | Hypnales | Bryopsida | https://ftp.cngb.org/pub/CNSA/data2/CNP0002895/CNS0539588/CNA0052778/ |
| *Thamnobryum sandei* | Hypnales | Bryopsida | https://ftp.cngb.org/pub/CNSA/data2/CNP0002895/CNS0538593/CNA0052779/ |
| *Regmatodon serrulatus* | Hypnales | Bryopsida | https://ftp.cngb.org/pub/CNSA/data2/CNP0002895/CNS0548205/CNA0052780/ |
| *Entodon concinnus* | Hypnales | Bryopsida | https://ftp.cngb.org/pub/CNSA/data2/CNP0002895/CNS0538594/CNA0052781/ |
| *Entodon seductrix* | Hypnales | Bryopsida | https://figshare.com/articles/dataset/Entodon_and_Hypnum_genome_project/17097077 |
| *Hypnum curvifolium* | Hypnales | Bryopsida | https://figshare.com/articles/dataset/Entodon_and_Hypnum_genome_project/17097077 |
| *Calohypnum plumiforme* | Hypnales | Bryopsida | https://ngdc.cncb.ac.cn/gwh/Assembly/8871/show |
| *Gollania philippinensis* | Hypnales | Bryopsida | https://ftp.cngb.org/pub/CNSA/data2/CNP0002895/CNS0538601/CNA0052782/ |
| *Ectropothecium obtusulum* | Hypnales | Bryopsida | https://ftp.cngb.org/pub/CNSA/data2/CNP0002895/CNS0538600/CNA0052783/ |
| *Hygrohypnum luridum* | Hypnales | Bryopsida | https://ftp.cngb.org/pub/CNSA/data2/CNP0002895/CNS0538604/CNA0052784/ |
| *Hygroamblystegium varium* | Hypnales | Bryopsida | https://ftp.cngb.org/pub/CNSA/data2/CNP0002895/CNS0539592/CNA0052785/ |
| *Cratoneuron filicinum* | Hypnales | Bryopsida | https://ftp.cngb.org/pub/CNSA/data2/CNP0002895/CNS0538605/CNA0052786/ |
| *Pleurozium schreberi* | Hypnales | Bryopsida | https://github.com/PycnopodiaD/Pleurozium_schreberi_annotated_genome_files/tree/master/assembly |
| *Physcomitrella patens* | Funariales | Bryopsida | https://phytozome-next.jgi.doe.gov/info/Ppatens_v6_1 |
| *Funaria hygrometrica* | Funariales | Bryopsida | https://doi.org/10.6084/m9.figshare.19720216 |
| *Physcomitrellopsis africana* | Funariales | Bryopsida | https://doi.org/10.6084/m9.figshare.25724079 |
| *Takakia lepidozioides* | Takakiales | Takakiopsida | https://www.takakia.com/download.html |
| *Lycopodium clavatum* | Lycopodiales | Lycopodiopsida | https://doi.org/10.6084/m9.figshare.20493417.v3 |
| *Huperzia asiatica* | Lycopodiales | Lycopodiopsida | https://figshare.com/projects/Huperzia_asiatica_genome/169145 |
| *Diphasiastrum complanatum* | Lycopodiales | Lycopodiopsida | https://phytozome-next.jgi.doe.gov/info/Dcomplanatum_v3_1 |
| *Selaginella lepidophylla* | Selaginellales | Lycopodiopsida | Personal correspondence with the first author Robert VanBuren |
| *Selaginella moellendorffii* | Selaginellales | Lycopodiopsida | https://phytozome-next.jgi.doe.gov/info/Smoellendorffii_v1_0 |
| *Selaginella tamariscina* | Selaginellales | Lycopodiopsida | Personal correspondence with first author Zhichao Xu |
| *Selaginella kraussiana* | Selaginellales | Lycopodiopsida | https://sk.ccgg.fun/download |
| *Isoetes taiwanensis* | Isoetales | Lycopodiopsida | https://genomevolution.org/coge/GenomeInfo.pl?gid=61511 |
| *Isoetes sinensis* | Isoetales | Lycopodiopsida | https://ftp.cngb.org/pub/CNSA/data5/CNP0004710/CNS0885329/CNA0072254/ |
| *Dipteris shenzhenensis* | Gleicheniales | Polypodiopsida | https://figshare.com/articles/dataset/The_genome_assembly_and_annotation_of_Dipteris_shenzhenensis/27419877/1 |
| *Alsophila spinulosa* | Cyatheales | Polypodiopsida | https://doi.org/10.6084/m9.figshare.19075346 |
| *Adiantum capillus-veneris* | Polypodiales | Polypodiopsida | https://www.ncbi.nlm.nih.gov/datasets/genome/GCA_014529385.2/ |
| *Ceratopteris richardii* | Polypodiales | Polypodiopsida | https://phytozome-next.jgi.doe.gov/info/Crichardii_v2_1 |
| *Azolla filiculoides* | Salviniales | Polypodiopsida | https://fernbase.org/ |
| *Salvinia cucullata* | Salviniales | Polypodiopsida | https://fernbase.org/ |
| *Marsilea vestita* | Salviniales | Polypodiopsida | https://fernbase.org/ |
| *Torreya grandis* | Cupressales | Pinopsida | https://doi.org/10.6084/m9.figshare.21089869 |
| *Taxus wallichiana* | Cupressales | Pinopsida | https://doi.org/10.5524/102659 |
| *Taxus chinensis* | Cupressales | Pinopsida | https://www.ncbi.nlm.nih.gov/datasets/genome/GCA_019776745.2/ |
| *Sequoiadendron giganteum* | Cupressales | Pinopsida | https://treegenesdb.org/FTP/Genomes/Segi/v2.0/ |
| *Pinus densiflora* | Pinales | Pinopsida | https://doi.org/10.25452/figshare.plus.25546534 |
| *Picea mariana* | Pinales | Pinopsida | https://doi.org/10.5281/zenodo.7830121 |
| *Ginkgo biloba* | Ginkgoales | Ginkgoopsida | https://ngdc.cncb.ac.cn/gwh/Assembly/18742/show |
| *Cycas panzhihuaensis* | Cycadales | Cycadopsida | https://db.cngb.org/codeplot/datasets/public_dataset?id=PwRftGHfPs5qG3gE |
| *Gnetum montanum* | Gnetales | Gnetopsida | https://doi.org/10.5061/dryad.ht76hdrdr |
| *Welwitschia mirabilis* | Welwitschiales | Gnetopsida | https://doi.org/10.5061/dryad.ht76hdrdr |
| *Amborella trichopoda* | Amborellales | Magnoliopsida | https://ngdc.cncb.ac.cn/gwh/Assembly/92602/show |
| *Brasenia schreberi* | Nymphaeales | Magnoliopsida | https://doi.org/10.6084/m9.figshare.27011128.v1 |
| *Nymphaea colorata* | Nymphaeales | Magnoliopsida | https://doi.org/10.6084/m9.figshare.27011128.v1 |
| *Euryale ferox* | Nymphaeales | Magnoliopsida | https://doi.org/10.6084/m9.figshare.22099592.v2 |
| *Ceratophyllum demersum* | Ceratophyllales | Magnoliopsida | https://genomevolution.org/coge/GenomeInfo.pl?gid=67253 |
| *Aristolochia fimbriata* | Piperales | Magnoliopsida | https://ngdc.cncb.ac.cn/gwh/Assembly/21819/show |
| *Piper nigrum* | Piperales | Magnoliopsida | https://cotton.hzau.edu.cn/EN/Download.htm |
| *Saururus chinensis* | Piperales | Magnoliopsida | https://doi.org/10.6084/m9.figshare.23735505.v1 |
| *Chloranthus spicatus* | Chloranthales | Magnoliopsida | https://ngdc.cncb.ac.cn/gwh/Assembly/23169/show |
| *Chloranthus sessilifolius* | Chloranthales | Magnoliopsida | https://github.com/yongzhiyang2012/Chloranthus-sessilifolius-genome/tree/main/Annotation/Chromosomes |
| *Warburgia ugandensis* | Canellales | Magnoliopsida | https://doi.org/10.6084/m9.figshare.23735505.v1 |
| *Annona cherimola* | Magnoliales | Magnoliopsida | https://ihsmsubtropicals.uma.es/easy_gdb/downloads.php |
| *Magnolia sieboldii* | Magnoliales | Magnoliopsida | https://doi.org/10.6084/m9.figshare.26825587.v1 |
| *Liriodendron chinense* | Magnoliales | Magnoliopsida | https://doi.org/10.6084/m9.figshare.27011128.v1 |
| *Persea americana* | Laurales | Magnoliopsida | https://doi.org/10.57760/sciencedb.07602 |
| *Chimonanthus salicifolius* | Laurales | Magnoliopsida | https://doi.org/10.6084/m9.figshare.27011128.v1 |
| *Acorus gramineus* | Acorales | Magnoliopsida | https://ngdc.cncb.ac.cn/gwh/Assembly/37749/show |
| *Wolffia australiana* | Alismatales | Magnoliopsida | https://www.lemna.org/download/Wo_australiana_8730/Wo_australiana_8730-REF-CSHL-1.0/ |
| *Acanthochlamys bracteata* | Pandanales | Magnoliopsida | https://doi.org/10.6084/m9.figshare.27011128.v1 |
| *Dioscorea alata* | Dioscoreales | Magnoliopsida | https://ftp.cngb.org/pub/CNSA/data6/CNP0006270/CNS1201353/CNA0263387/ |
| *Angraecum sesquipedale* | Asparagales | Magnoliopsida | source:https://ngdc.cncb.ac.cn/gwh/Assembly/88029/show |
| *Gloriosa superba* | Liliales | Magnoliopsida | https://doi.org/10.6084/m9.figshare.27933375.v1 |
| *Lilium sargentiae* | Liliales | Magnoliopsida | https://doi.org/10.6084/m9.figshare.27933375.v1 |
| *Oryza sativa* | Poales | Magnoliopsida | https://phytozome-next.jgi.doe.gov/info/Osativa_v7_0 |
| *Vriesea erythrodactylon* | Poales | Magnoliopsida | https://doi.org/10.6084/m9.figshare.25887199.v4 |
| *Elaeis guineensis* | Arecales | Magnoliopsida | http://genomsawit.mpob.gov.my |
| *Pontederia cordata* | Commelinales | Magnoliopsida | https://doi.org/10.6084/m9.figshare.24866487.v1 |
| *Musa schizocarpa* | Zingiberales | Magnoliopsida | https://banana-genome-hub.southgreen.fr/node/50/3777229 |
| *Stephania japonica* | Ranunculales | Magnoliopsida | http://eegr.bio2db.com/Download-Sequence |
| *Protea cynaroides* | Proteales | Magnoliopsida | https://doi.org/10.6084/m9.figshare.27011128.v1 |
| *Tetracentron sinense* | Trochodendrales | Magnoliopsida | https://doi.org/10.6084/m9.figshare.27011128.v1 |
| *Buxus austroyunnanensis* | Buxales | Magnoliopsida | https://doi.org/10.6084/m9.figshare.27011128.v1 |
| *Cornus wilsoniana* | Cornales | Magnoliopsida | https://doi.org/10.6084/m9.figshare.27011128.v1 |
| *Rhododendron liliiflorum* | Ericales | Magnoliopsida | http://www.tegr.com.cn/Download-Sequence |
| *Apocynum pictum* | Gentianales | Magnoliopsida | https://doi.org/10.6084/m9.figshare.25060931.v1 |
| *Dracocephalum rupestre* | Lamiales | Magnoliopsida | https://doi.org/10.6084/m9.figshare.25778184.v2 |
| *Ehretia macrophylla* | Boraginales | Magnoliopsida | https://ngdc.cncb.ac.cn/gwh/Assembly/83111/show |
| *Eucommia ulmoides* | Garryales | Magnoliopsida | https://ngdc.cncb.ac.cn/gwh/Assembly/13/show |
| *Solanum lycopersicum* | Solanales | Magnoliopsida | https://phytozome-next.jgi.doe.gov/info/Slycopersicum_ITAG5_0 |
| *Nicotiana benthamiana* | Solanales | Magnoliopsida | https://ngdc.cncb.ac.cn/gwh/Assembly/26127/show |
| *Helwingia omeiensis* | Aquifoliales | Magnoliopsida | https://doi.org/10.6084/m9.figshare.22817414.v3 |
| *Lactuca sativa* | Asterales | Magnoliopsida | https://figshare.com/s/f5f0e8068d5a236ea408 |
| *Centella asiatica* | Apiales | Magnoliopsida | https://doi.org/10.6084/m9.figshare.28022099.v1 |
| *Triplostegia glandulifera* | Dipsacales | Magnoliopsida | https://doi.org/10.6084/m9.figshare.25018103.v1 |
| *Escallonia herrerae* | Escalloniales | Magnoliopsida | https://www.ncbi.nlm.nih.gov/datasets/genome/GCA_033070095.1/ |
| *Malania oleifera* | Santalales | Magnoliopsida | https://ngdc.cncb.ac.cn/gwh/Assembly/24427/show |
| *Beta vulgaris* | Caryophyllales | Magnoliopsida | https://phytozome-next.jgi.doe.gov/info/Bvulgarisssp_vulgaris_EL10_2_2 |
| *Liquidambar styraciflua* | Saxifragales | Magnoliopsida | https://doi.org/10.6084/m9.figshare.26076514.v1 |
| *Vitis rotundifolia* | Vitales | Magnoliopsida | https://zenodo.org/records/7944875 |
| *Glycine max* | Fabales | Magnoliopsida | https://phytozome-next.jgi.doe.gov/info/Gmax_Wm82_a6_v1 |
| *Malus doumeri* | Rosales | Magnoliopsida | https://ngdc.cncb.ac.cn/gwh/Assembly/34332/show |
| *Quercus variabilis* | Fagales | Magnoliopsida | https://ngdc.cncb.ac.cn/gwh/Assembly/30668/show |
| *Citrullus ecirrhosus* PI_673135_v2 | Cucurbitales | Magnoliopsida | http://www.watermelondb.cn/#/download |
| *Tetraena mongolica* | Zygophyllales | Magnoliopsida | https://doi.org/10.6084/m9.figshare.20463798.v1 |
| *Lagerstroemia speciosa* | Myrtales | Magnoliopsida | https://doi.org/10.6084/m9.figshare.26861248.v1 |
| *Euscaphis japonica* | Crossosomatales | Magnoliopsida | https://doi.org/10.6084/m9.figshare.27011128.v1 |
| *Populus tremula* | Malpighiales | Magnoliopsida | https://phytozome-next.jgi.doe.gov/info/PtremulaxPopulusalbaHAP1_v5_1 |
| *Averrhoa carambola* | Oxalidales | Magnoliopsida | https://doi.org/10.6084/m9.figshare.27011128.v1 |
| *Tripterygium wilfordii* | Celastrales | Magnoliopsida | https://www.ncbi.nlm.nih.gov/datasets/genome/GCF_013401445.1 |
| *Gossypium longicalyx* | Malvales | Magnoliopsida | https://doi.org/10.6084/m9.figshare.17280200.v1 |
| *Arabidopsis thaliana* | Brassicales | Magnoliopsida | https://phytozome-next.jgi.doe.gov/info/Athaliana_TAIR10 |
| *Acer yangbiense* | Sapindales | Magnoliopsida | https://doi.org/10.6084/m9.figshare.27011128.v1 |
| *Iodes seguinii* | Icacinales | Magnoliopsida | Personal correspondence with Min Tang |

**Table S3. *R*^2^ of the Michaelis–Menten saturation curve and the exponential saturation curve on the p distance against the corrected distance**

| **Removed outgroup representatives** | **Michaelis–Menten saturation curve** | **Exponential saturation curve** |
| --- | --- | --- |
| None | 0.7946 | 0.7750 |
| *Chlamydomonas reinhardtii* | 0.8029 | 0.7847 |
| *Chlamydomonas reinhardtii* and *Chlorokybus atmophyticus* | 0.8059 | 0.7877 |
| *Chlamydomonas reinhardtii*, *Chlorokybus atmophyticus*, and *Mesostigma viride* | 0.8161 | 0.7988 |
| *Chlamydomonas reinhardtii*, *Chlorokybus atmophyticus*, *Mesostigma viride*, and *Klebsormidium nitens* | 0.8131 | 0.7956 |
| *Chlamydomonas reinhardtii*, *Chlorokybus atmophyticus*, *Mesostigma viride*, *Klebsormidium nitens* and *Chara braunii* | 0.8156 | 0.7981 |
| *Chlamydomonas reinhardtii*, *Chlorokybus atmophyticus*, *Mesostigma viride*, *Klebsormidium nitens*, *Chara braunii*, and *Spirogloea muscicola* | 0.8153 | 0.7980 |
| *Chlamydomonas reinhardtii*, *Chlorokybus atmophyticus*, *Mesostigma viride*, *Klebsormidium nitens*, *Chara braunii*, *Spirogloea muscicola*, and *Penium margaritaceum* | 0.8179 | 0.7996 |
| *Chlamydomonas reinhardtii*, *Chlorokybus atmophyticus*, *Mesostigma viride*, *Klebsormidium nitens*, *Chara braunii*, *Spirogloea muscicola*, *Penium margaritaceum*, and *Mesotaenium endlicherianum* | 0.8228 | 0.8030 |
| *Chlamydomonas reinhardtii*, *Chlorokybus atmophyticus*, *Mesostigma viride*, *Klebsormidium nitens*, *Chara braunii*, *Spirogloea muscicola*, *Penium margaritaceum*, *Mesotaenium endlicherianum* and *Zygnema cf. cylindricum* | 0.8176 | 0.7992 |
| *Chlamydomonas reinhardtii*, *Chlorokybus atmophyticus*, *Mesostigma viride*, *Klebsormidium nitens*, *Chara braunii*, *Spirogloea muscicola*, *Penium margaritaceum*, *Mesotaenium endlicherianum*, *Zygnema cf. cylindricum* and *Z. circumcarinatum UTEX 1560* | 0.8168 | 0.7982 |
| *Chlamydomonas reinhardtii*, *Chlorokybus atmophyticus*, *Mesostigma viride*, *Klebsormidium nitens*, *Chara braunii*, *Spirogloea muscicola*, *Penium margaritaceum*, *Mesotaenium endlicherianum*, *Zygnema cf. cylindricum*, *Z. circumcarinatum UTEX 1560*, and *Z. circumcarinatum SAG 698-1b* | 0.8244 | 0.8039 |
