## Supplementary Information for "Rooting the deep divergence of land plants"

**Supplementary Information 1 Species trees inferred from the concatenation and MSC methods with stepwise removal of outgroup representatives**

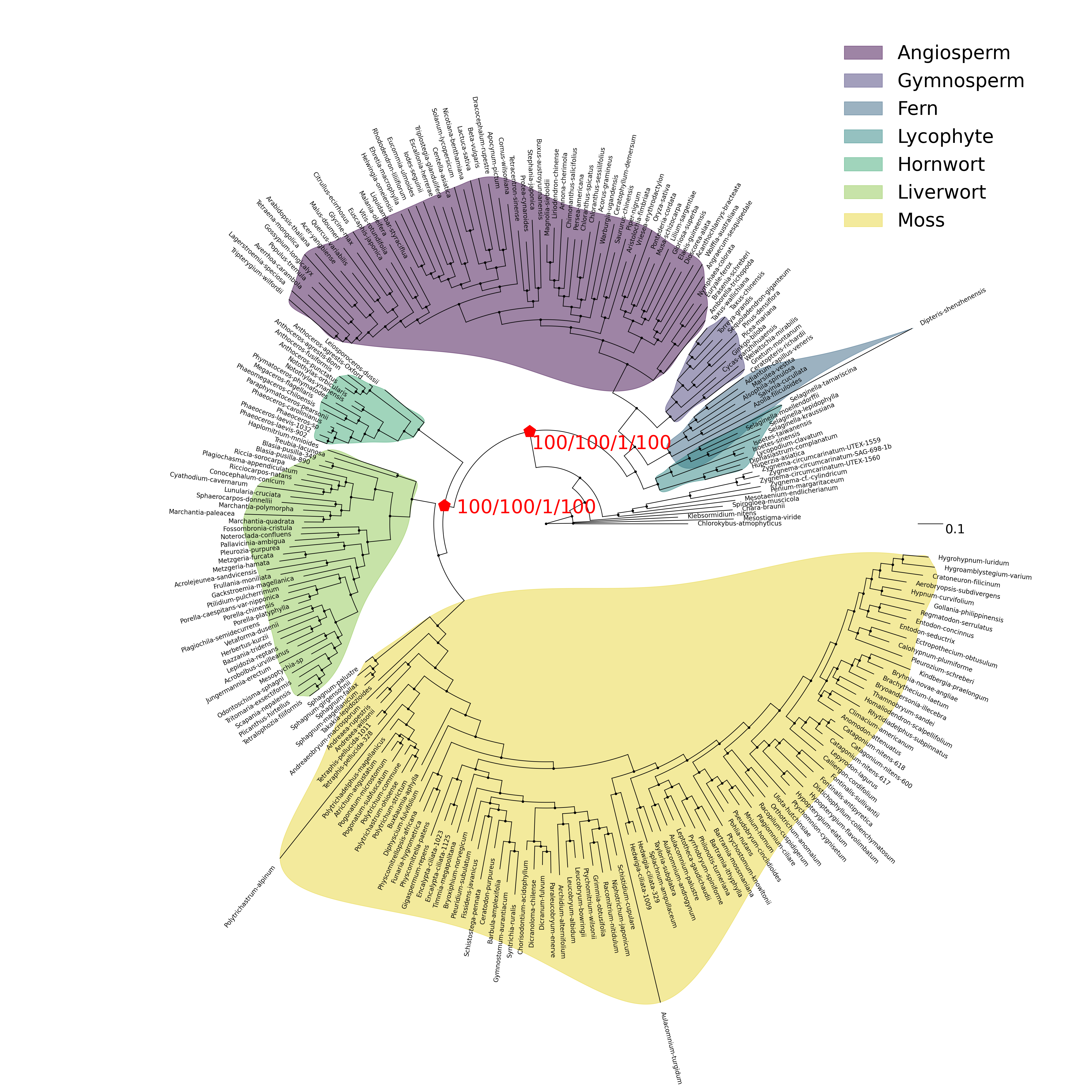

**Figure SI 1. Species tree inferred by the concatenation method based on the 137-rSOG dataset with *Chlamydomonas reinhardtii* removed.**

Taxonomical affiliations to hornworts, liverworts, mosses, lycophytes, ferns, angiosperms, and gymnosperms are marked with distinct colors. The supporting level for the crown node of monophyletic bryophytes and embryophytes is presented in terms of the ultrafast bootstrap support, SH-like approximate likelihood ratio test, approximate Bayes test, and local bootstrap probability, respectively. The branch length represents the number of substitutions per site.

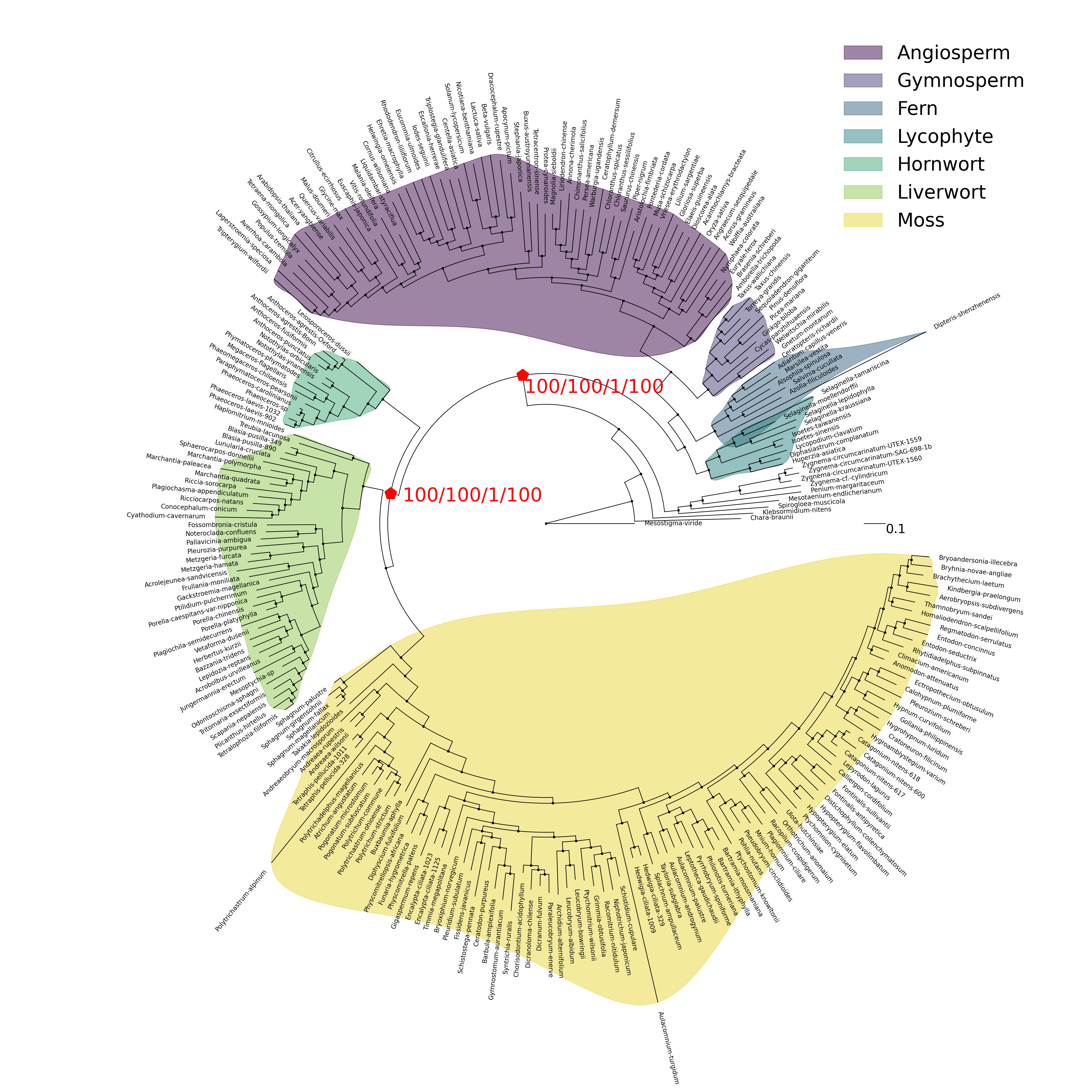

**Figure SI 2. Species tree inferred by the concatenation method based on the 137-rSOG dataset with *Chlamydomonas reinhardtii* and** ***Chlorokybus atmophyticus* removed.**

The caption stays the same as Figure SI 1.

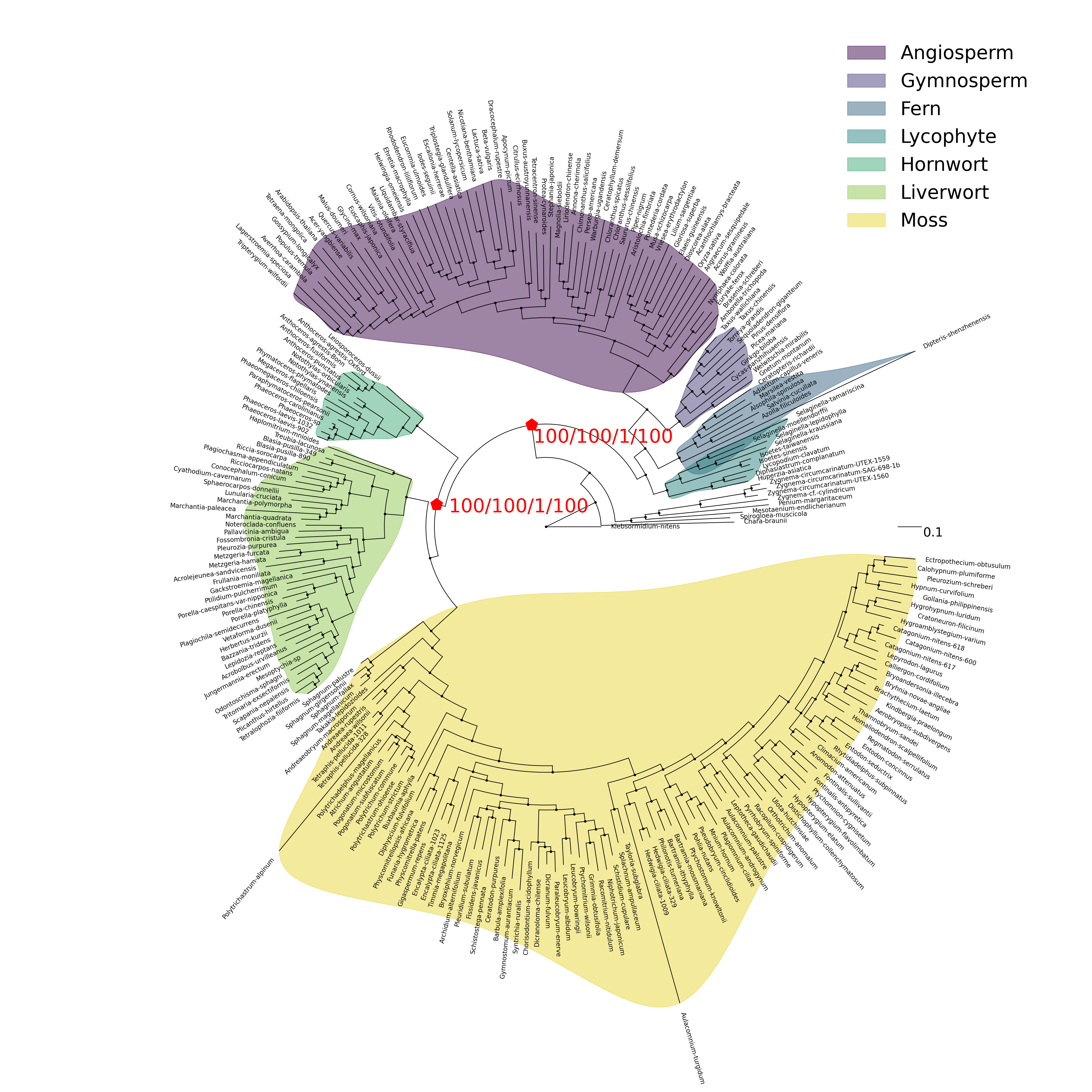

**Figure SI 3. Species tree inferred by the concatenation method based on the 137-rSOG dataset with *Chlamydomonas reinhardtii*,** ***Chlorokybus atmophyticus*, and *Mesostigma viride* removed.**

The caption stays the same as Figure SI 1.

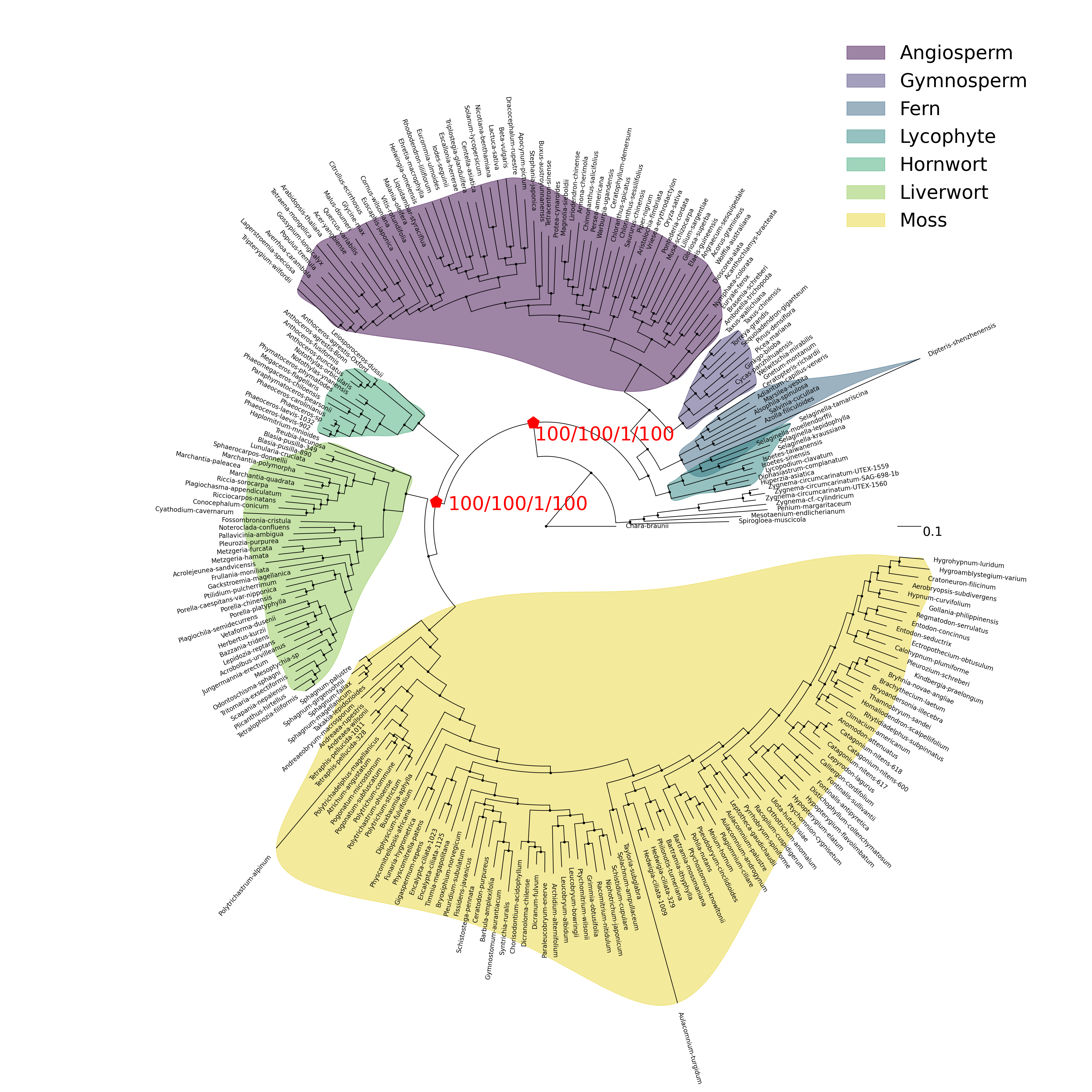

**Figure SI 4. Species tree inferred by the concatenation method based on the 137-rSOG dataset with *Chlamydomonas reinhardtii*,** ***Chlorokybus atmophyticus*, *Mesostigma viride*, and *Klebsormidium nitens* removed.**

The caption stays the same as Figure SI 1.

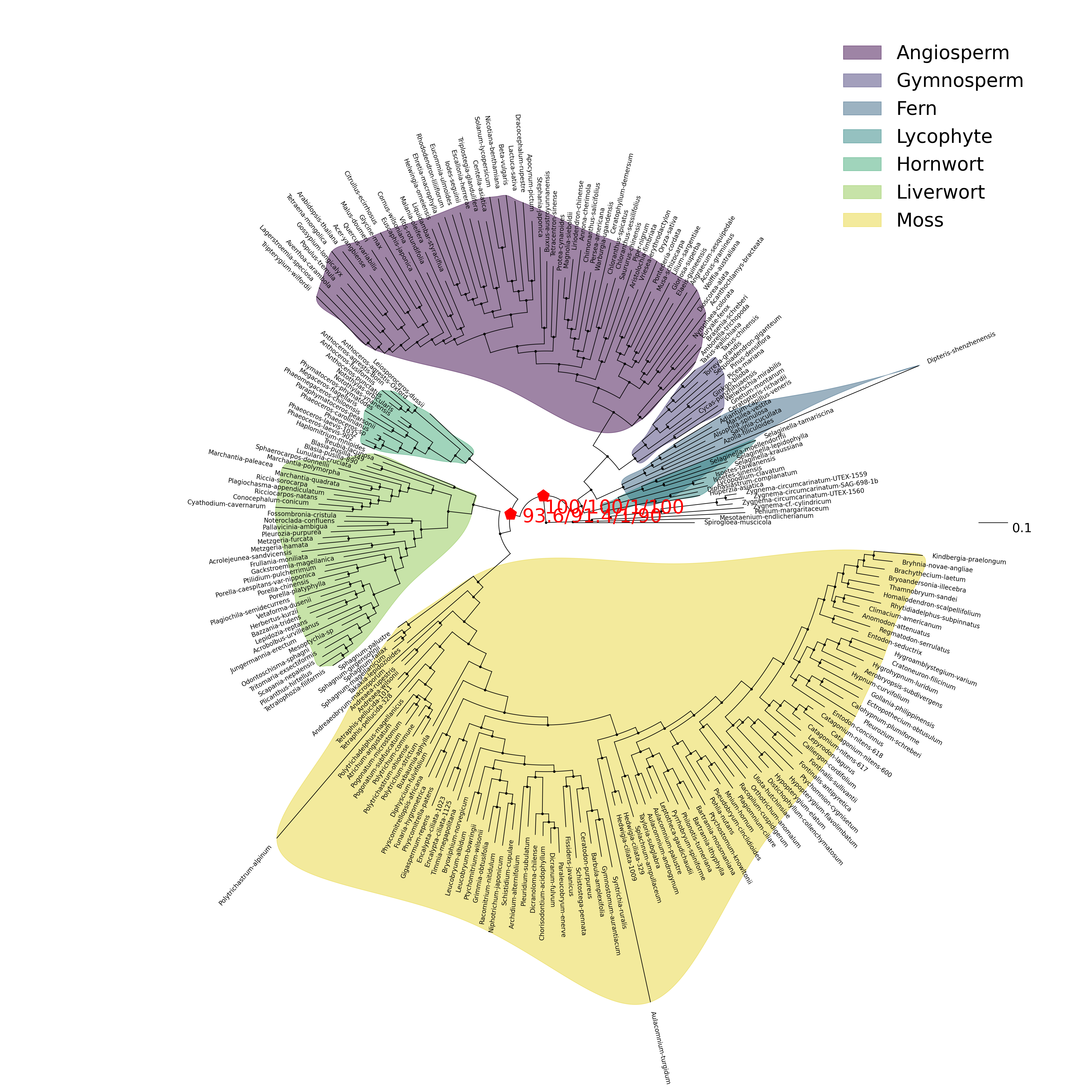

**Figure SI 5. Species tree inferred by the concatenation method based on the 137-rSOG dataset with *Chlamydomonas reinhardtii*,** ***Chlorokybus atmophyticus*, *Mesostigma viride*, *Klebsormidium nitens* and *Chara braunii* removed.**

The caption stays the same as Figure SI 1.

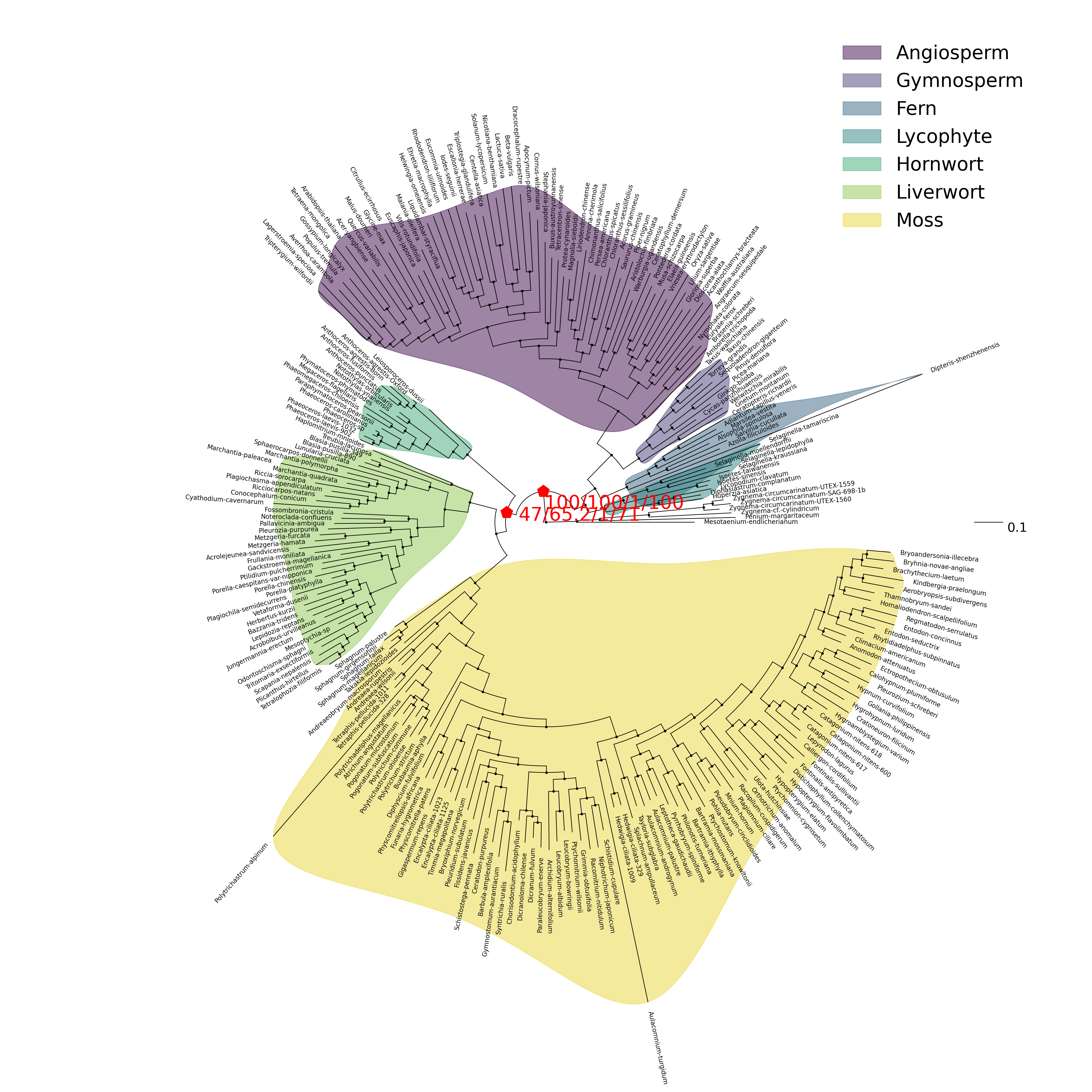

**Figure SI 6. Species tree inferred by the concatenation method based on the 137-rSOG dataset with *Chlamydomonas reinhardtii*,** ***Chlorokybus atmophyticus*, *Mesostigma viride*, *Klebsormidium nitens*, *Chara braunii*, and *Spirogloea muscicola* removed.**

The caption stays the same as Figure SI 1.

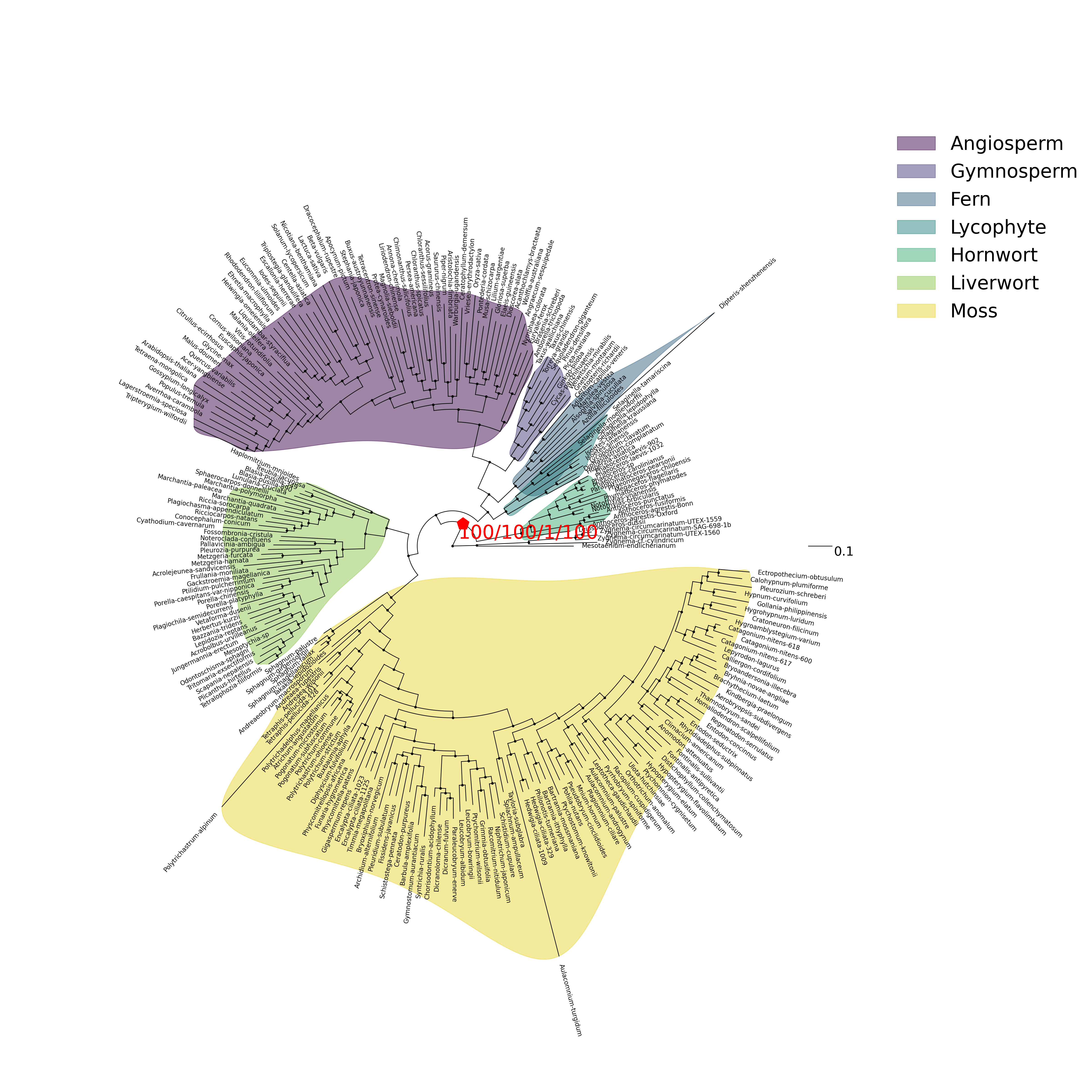

**Figure SI 7. Species tree inferred by the concatenation method based on the 137-rSOG dataset with *Chlamydomonas reinhardtii*,** ***Chlorokybus atmophyticus*, *Mesostigma viride*, *Klebsormidium nitens*, *Chara braunii*, *Spirogloea muscicola*, and *Penium margaritaceum* removed.**

Taxonomical affiliations to hornworts, liverworts, mosses, lycophytes, ferns, angiosperms, and gymnosperms are marked with distinct colors. The supporting level for the crown node of paraphyletic bryophytes is presented in terms of the ultrafast bootstrap support, SH-like approximate likelihood ratio test, approximate Bayes test, and local bootstrap probability, respectively. The branch length represents the number of substitutions per site.

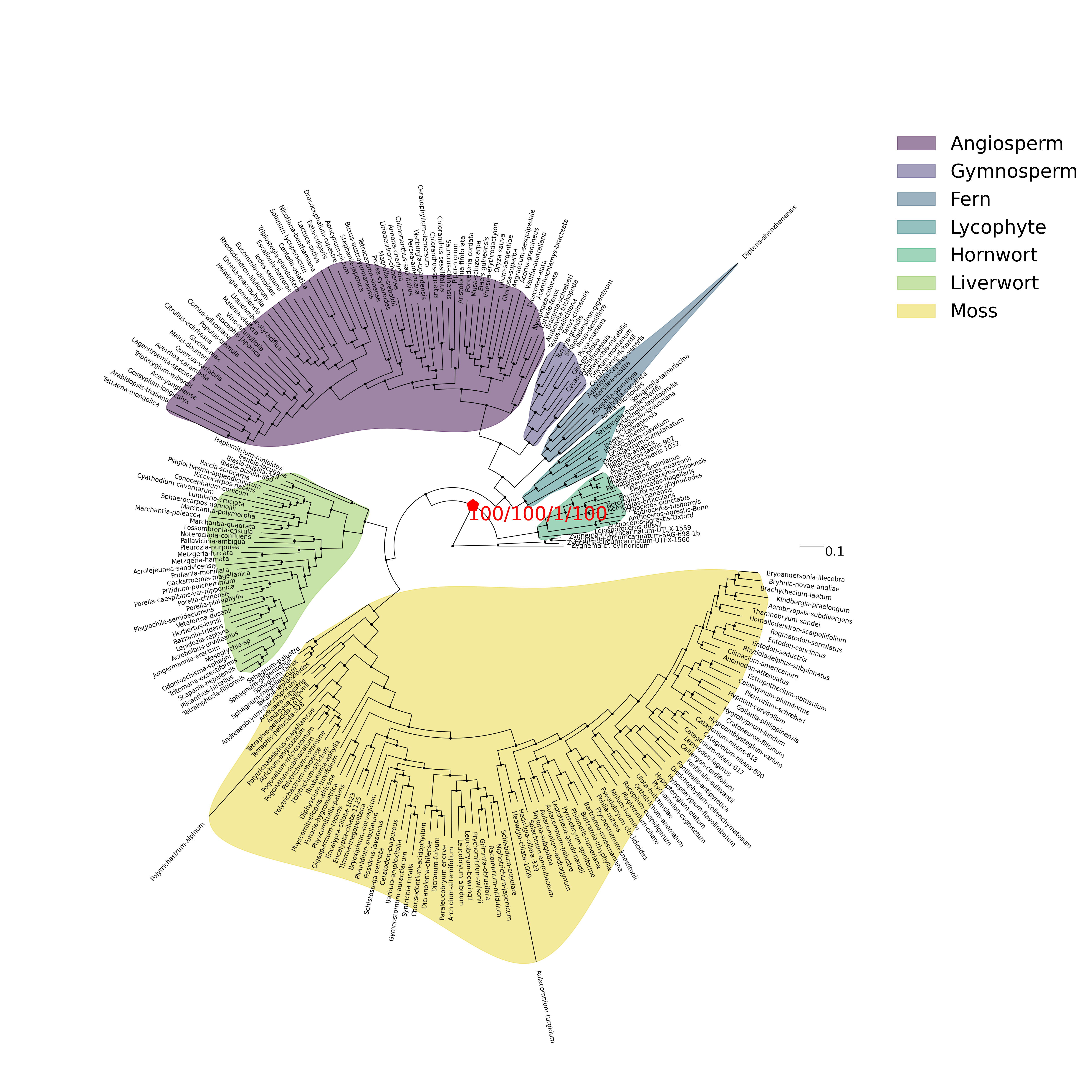

**Figure SI 8. Species tree inferred by the concatenation method based on the 137-rSOG dataset with *Chlamydomonas reinhardtii*,** ***Chlorokybus atmophyticus*, *Mesostigma viride*, *Klebsormidium nitens*, *Chara braunii*, *Spirogloea muscicola*, *Penium margaritaceum*, and *Mesotaenium endlicherianum* removed.**

The caption stays the same as Figure SI 7.

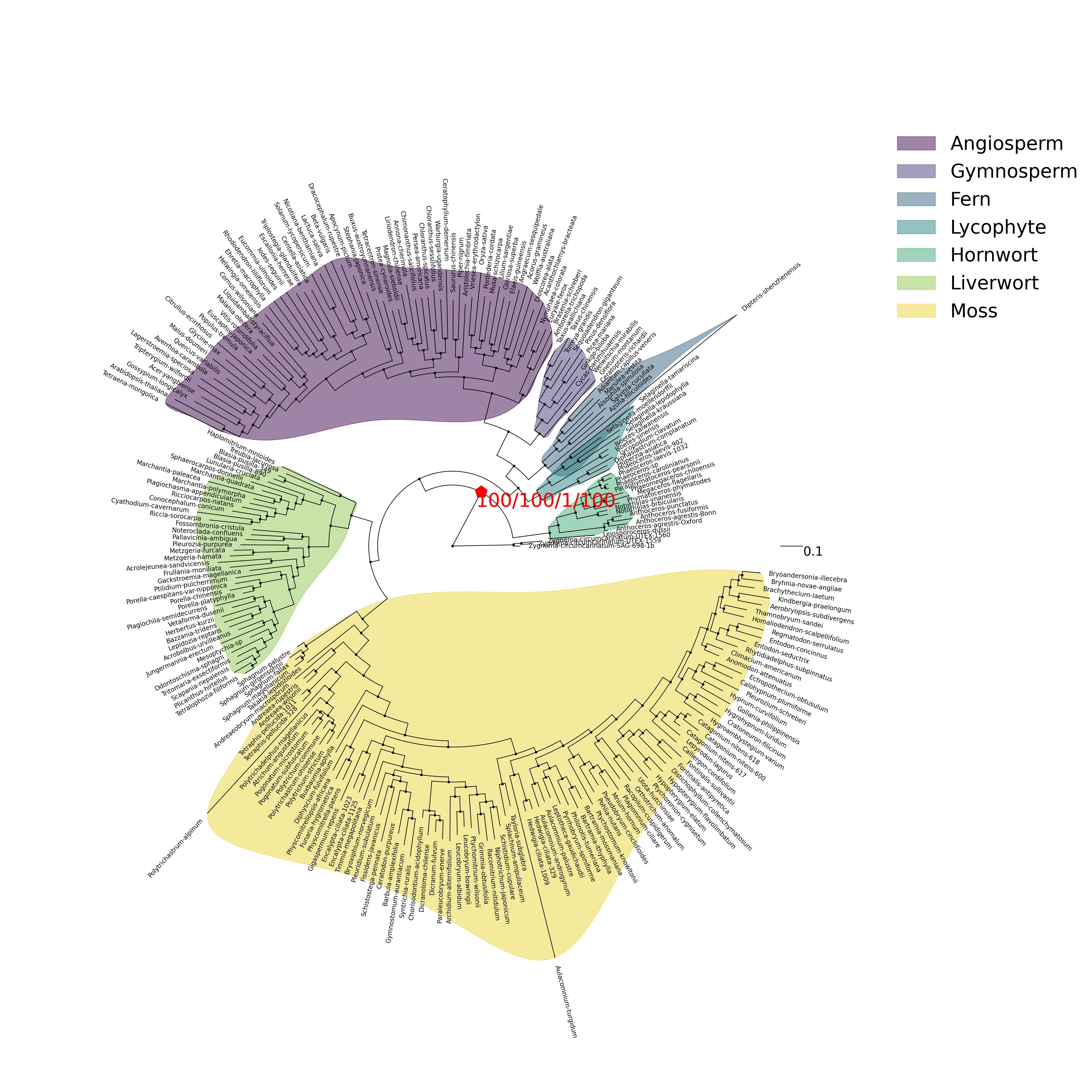

**Figure SI 9. Species tree inferred by the concatenation method based on the 137-rSOG dataset with *Chlamydomonas reinhardtii*,** ***Chlorokybus atmophyticus*, *Mesostigma viride*, *Klebsormidium nitens*, *Chara braunii*, *Spirogloea muscicola*, *Penium margaritaceum*, *Mesotaenium endlicherianum* and *Zygnema cf. cylindricum* removed.**

The caption stays the same as Figure SI 7.

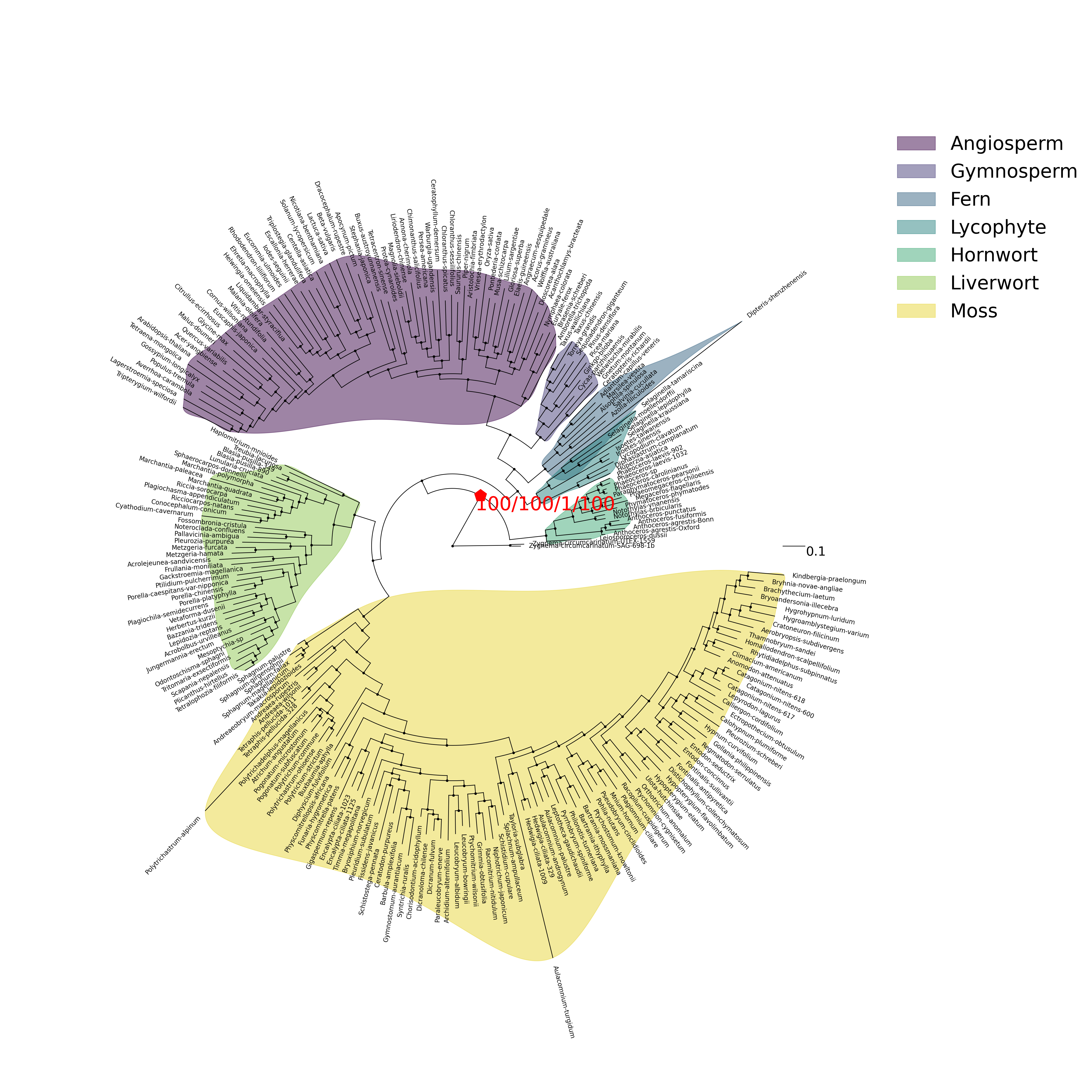

**Figure SI 10. Species tree inferred by the concatenation method based on the 137-rSOG dataset with *Chlamydomonas reinhardtii*,** ***Chlorokybus atmophyticus*, *Mesostigma viride*, *Klebsormidium nitens*, *Chara braunii*, *Spirogloea muscicola*, *Penium margaritaceum*, *Mesotaenium endlicherianum*, *Zygnema cf. cylindricum* and *Z. circumcarinatum UTEX 1560* removed.**

The caption stays the same as Figure SI 7.

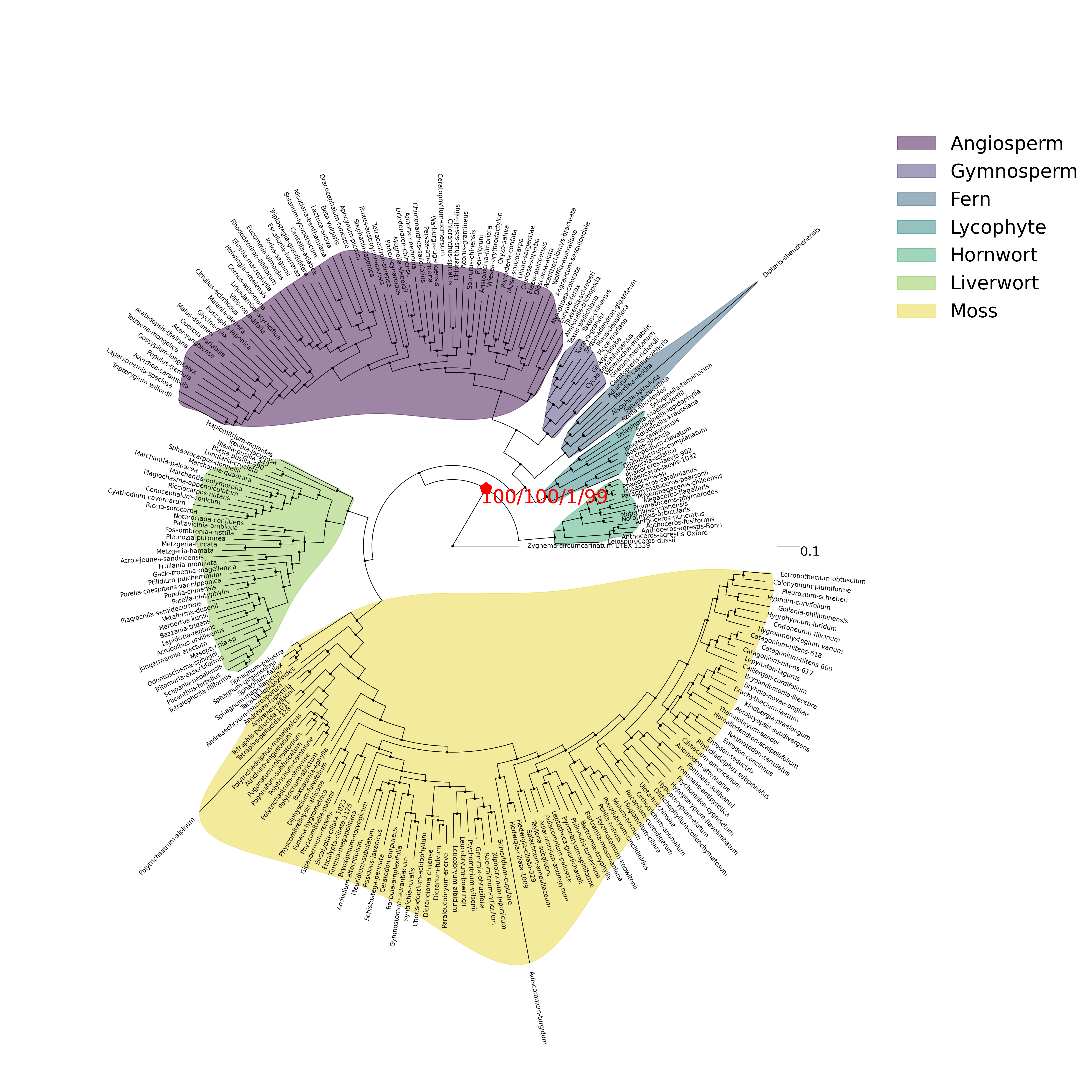

**Figure SI 11. Species tree inferred by the concatenation method based on the 137-rSOG dataset with *Chlamydomonas reinhardtii*,** ***Chlorokybus atmophyticus*, *Mesostigma viride*, *Klebsormidium nitens*, *Chara braunii*, *Spirogloea muscicola*, *Penium margaritaceum*, *Mesotaenium endlicherianum*, *Zygnema cf. cylindricum*, *Z. circumcarinatum UTEX 1560*, and *Z. circumcarinatum SAG 698-1b* removed.**

The caption stays the same as Figure SI 7.

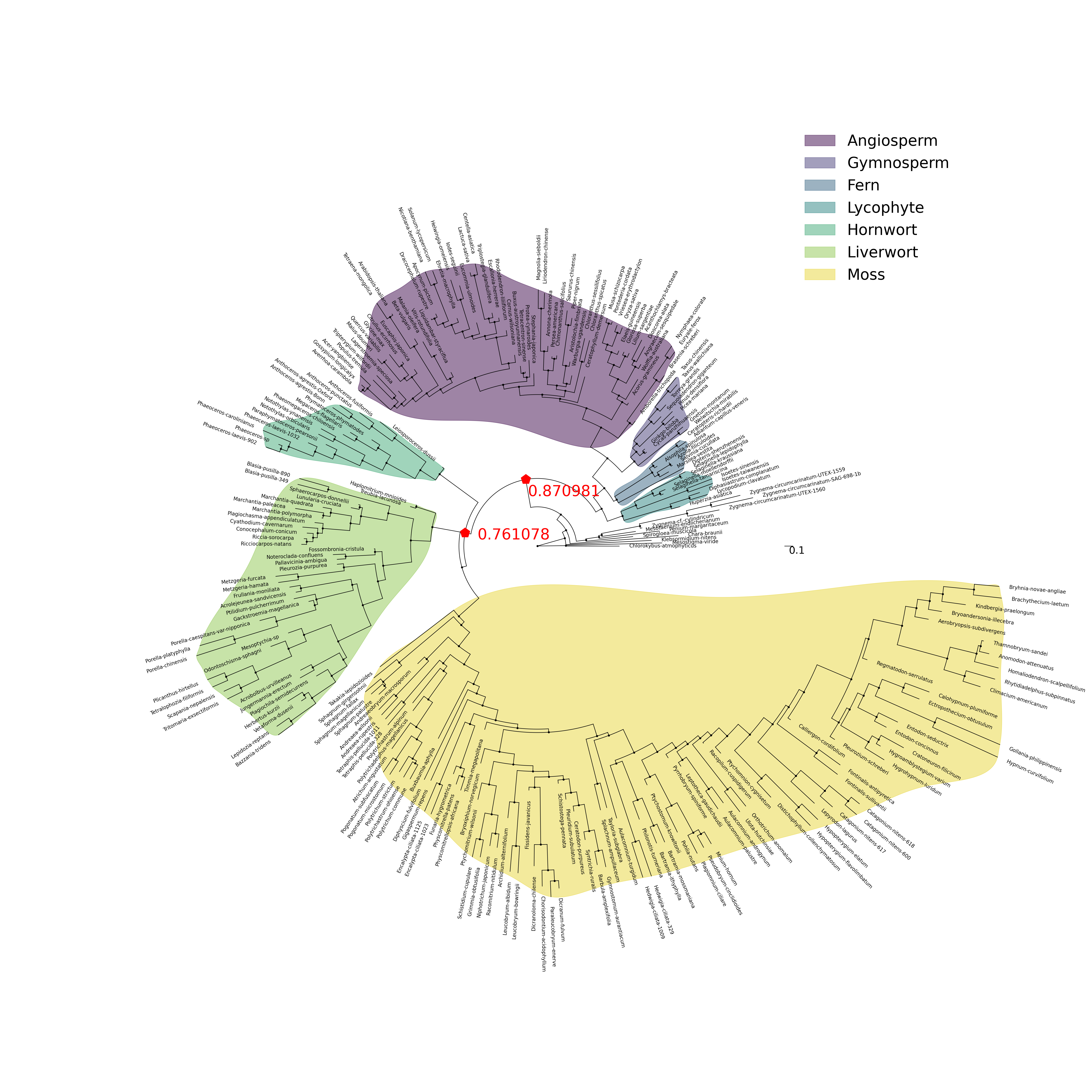

**Figure SI 12. Species tree inferred by the MSC method based on the 137-rSOG dataset with *Chlamydomonas reinhardtii* removed.**

Taxonomical affiliations to hornworts, liverworts, mosses, lycophytes, ferns, angiosperms, and gymnosperms are marked with distinct colors. The supporting level for the crown node of bryophytes and embryophytes is measured by the local posterior probability. The branch length represents the number of coalescent unit.

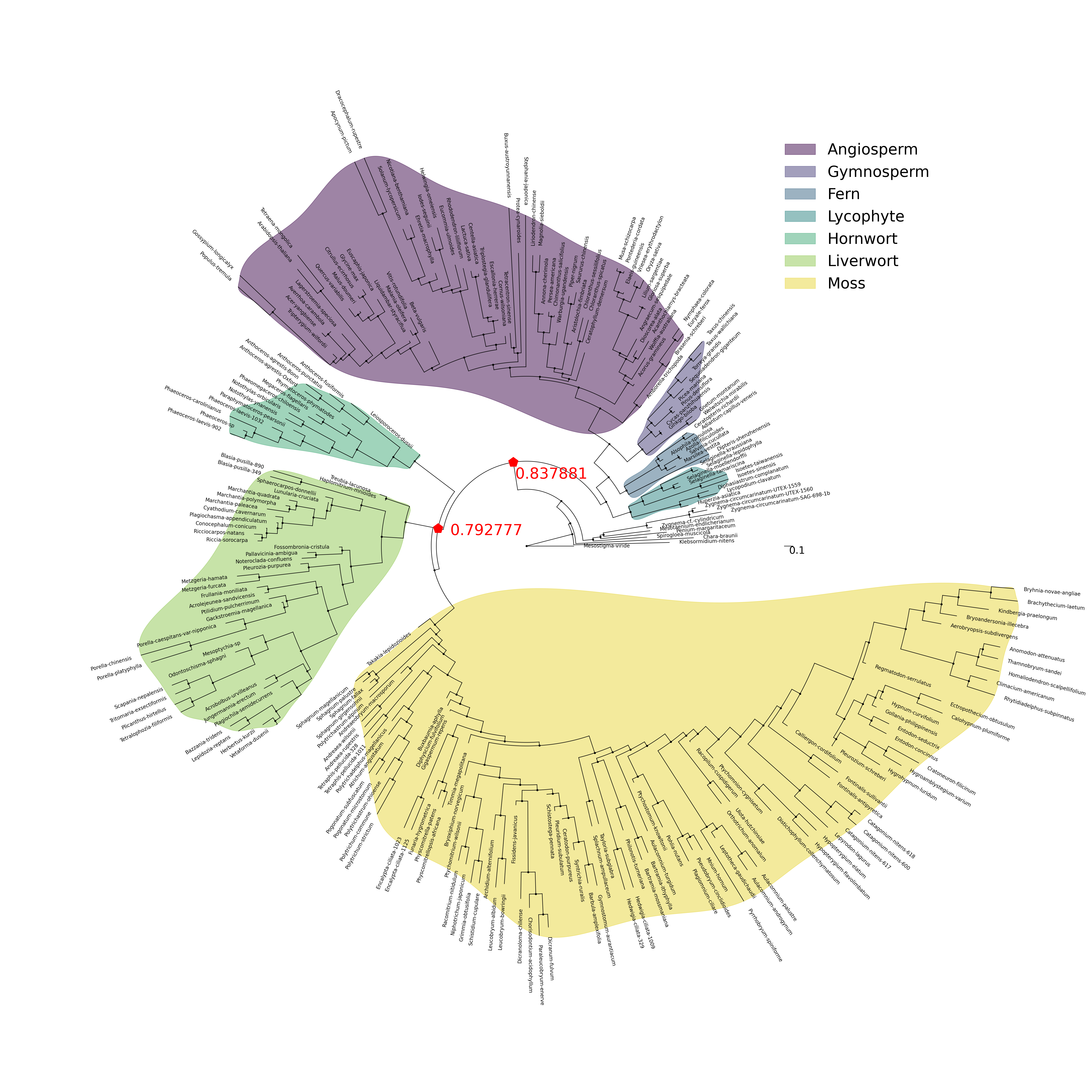

**Figure SI 13. Species tree inferred by the MSC method based on the 137-rSOG dataset with *Chlamydomonas reinhardtii* and** ***Chlorokybus atmophyticus* removed.**

The caption stays the same as Figure SI 12.

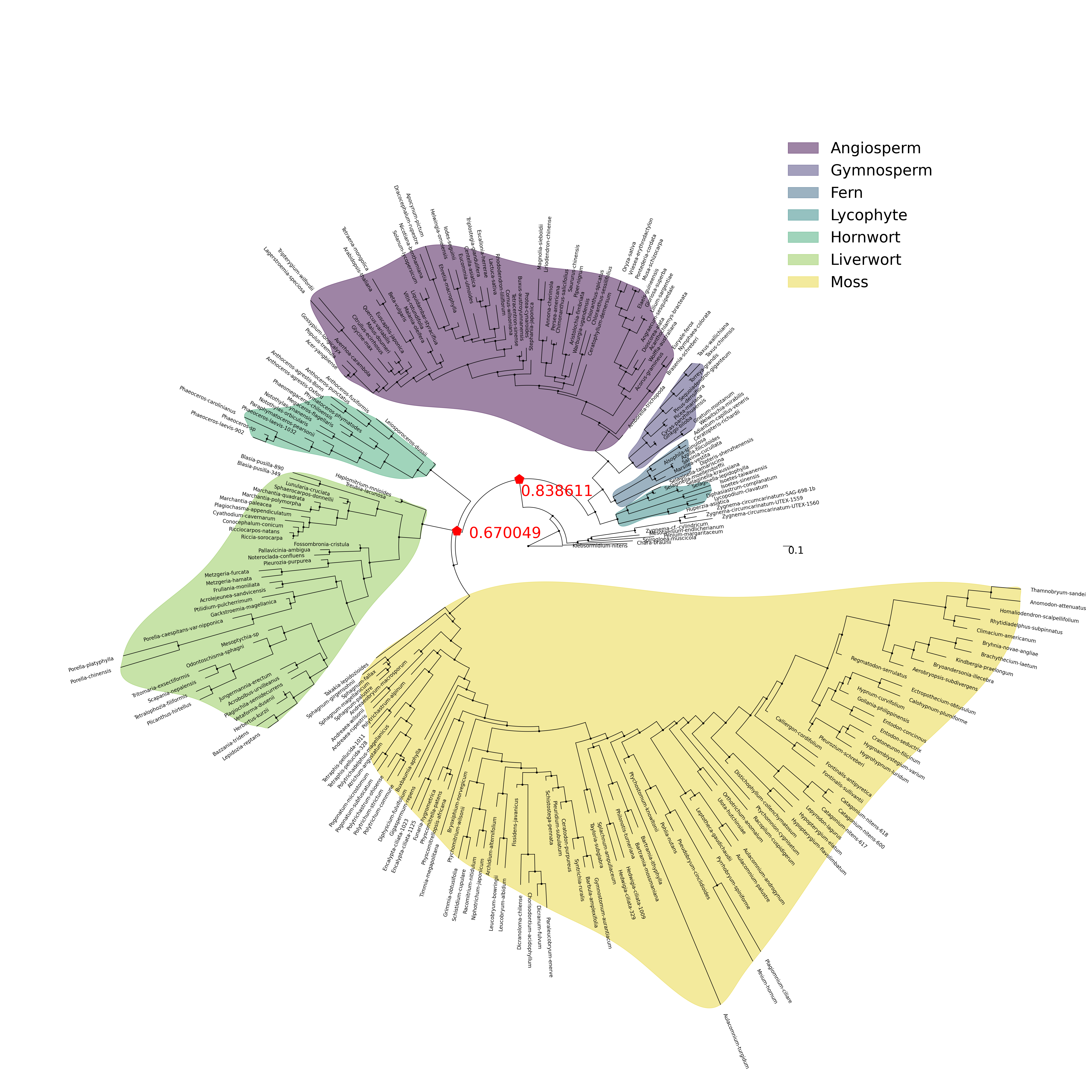

**Figure SI 14. Species tree inferred by the MSC method based on the 137-rSOG dataset with *Chlamydomonas reinhardtii*,** ***Chlorokybus atmophyticus*, and *Mesostigma viride* removed.**

The caption stays the same as Figure SI 12.

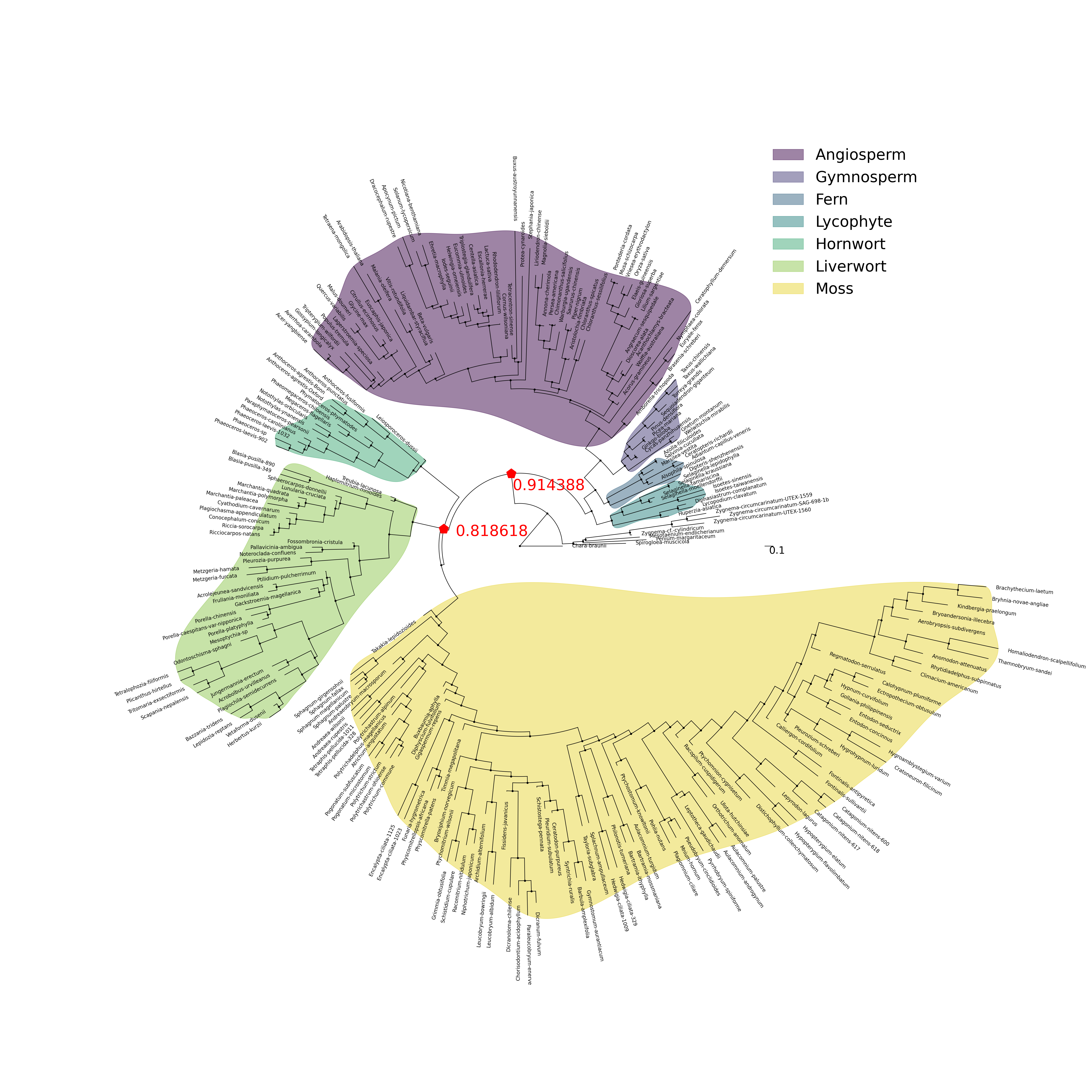

**Figure SI 15. Species tree inferred by the MSC method based on the 137-rSOG dataset with *Chlamydomonas reinhardtii*,** ***Chlorokybus atmophyticus*, *Mesostigma viride*, and *Klebsormidium nitens* removed.**

The caption stays the same as Figure SI 12.

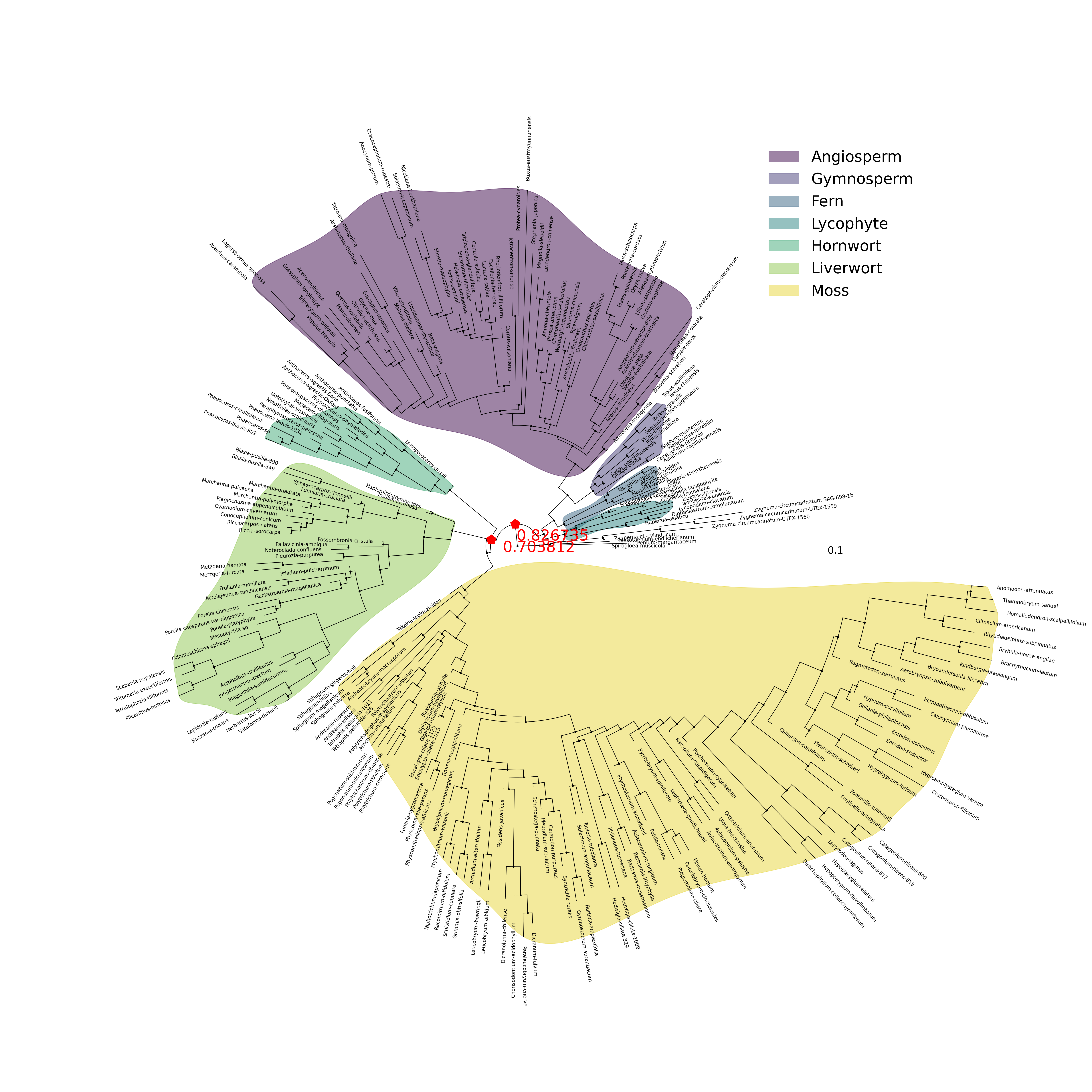

**Figure SI 16. Species tree inferred by the MSC method based on the 137-rSOG dataset with *Chlamydomonas reinhardtii*,** ***Chlorokybus atmophyticus*, *Mesostigma viride*, *Klebsormidium nitens*, and *Chara braunii* removed.**

The caption stays the same as Figure SI 12.

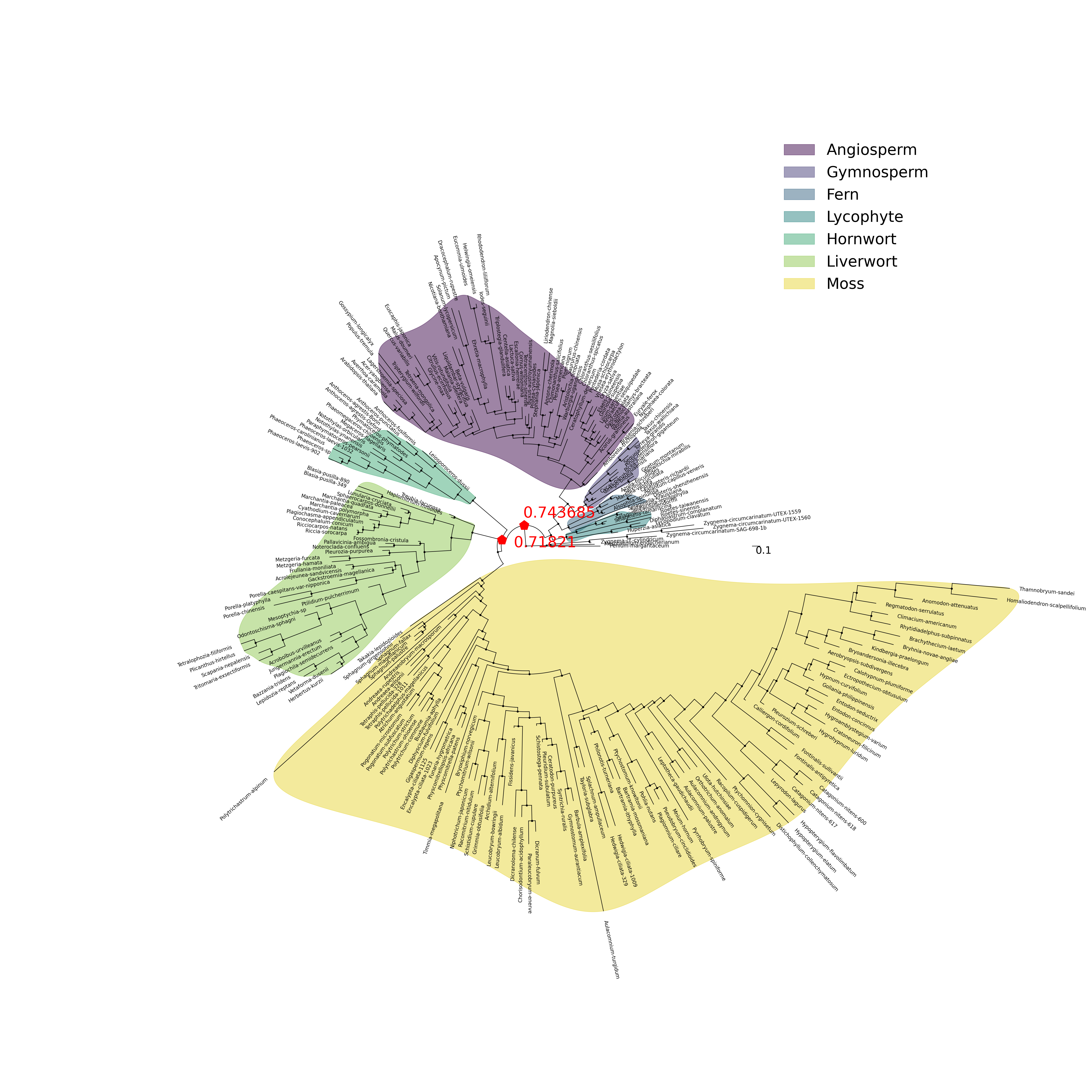

**Figure SI 17. Species tree inferred by the MSC method based on the 137-rSOG dataset with *Chlamydomonas reinhardtii*,** ***Chlorokybus atmophyticus*, *Mesostigma viride*, *Klebsormidium nitens*, *Chara braunii*, and *Spirogloea muscicola* removed.**

The caption stays the same as Figure SI 12.

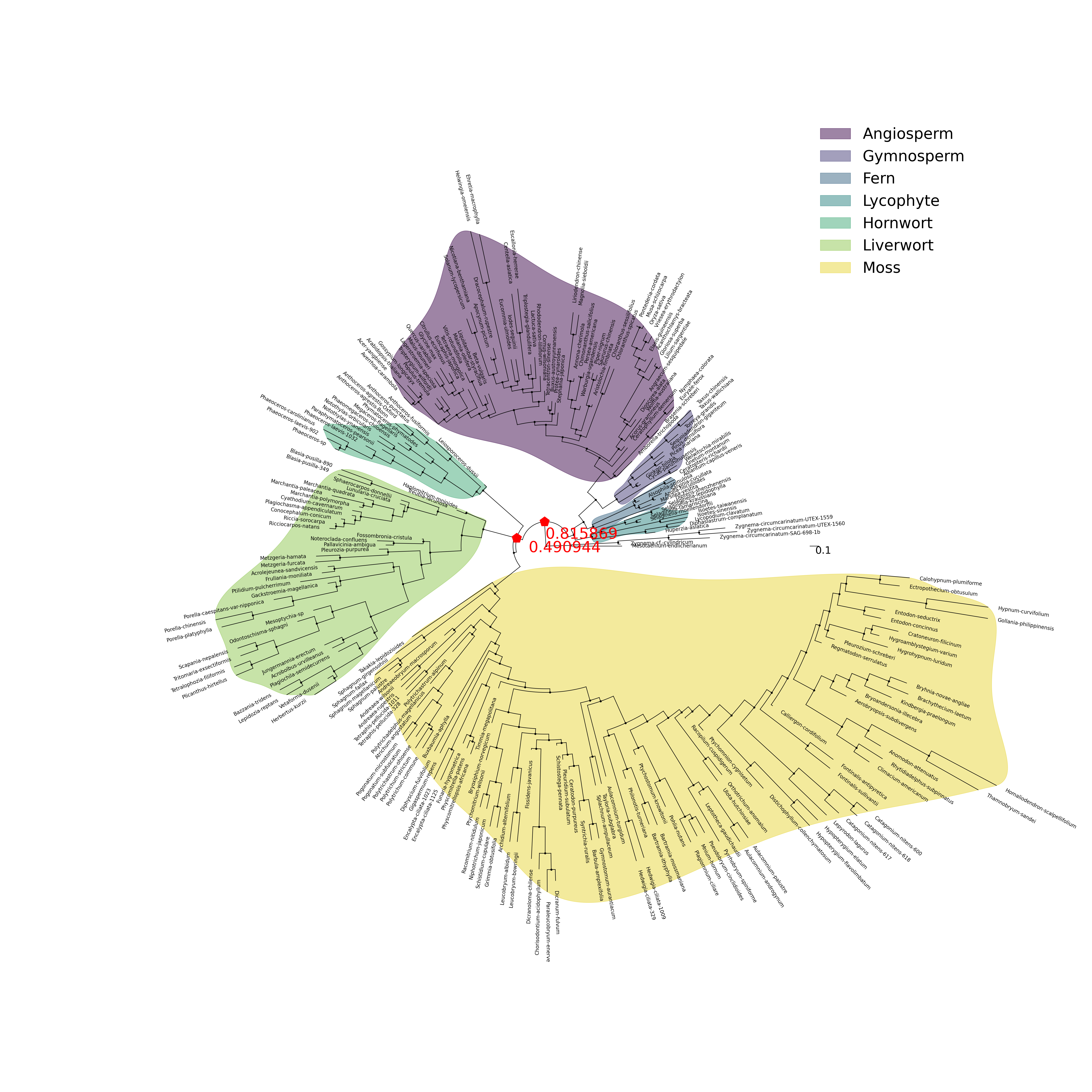

**Figure SI 18. Species tree inferred by the MSC method based on the 137-rSOG dataset with *Chlamydomonas reinhardtii*,** ***Chlorokybus atmophyticus*, *Mesostigma viride*, *Klebsormidium nitens*, *Chara braunii*, *Spirogloea muscicola*, and *Penium margaritaceum* removed.**

The caption stays the same as Figure SI 12.

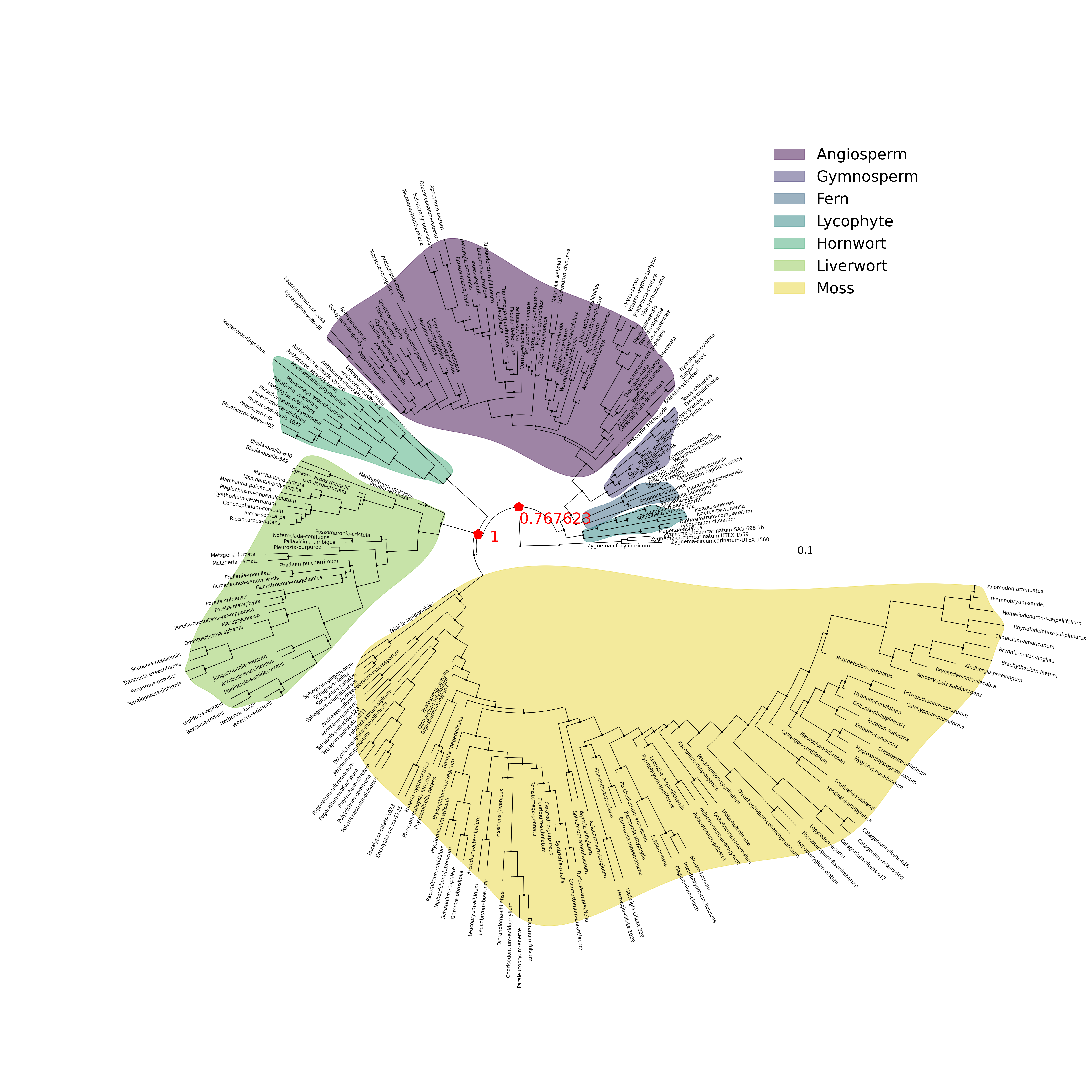

**Figure SI 19. Species tree inferred by the MSC method based on the 137-rSOG dataset with *Chlamydomonas reinhardtii*,** ***Chlorokybus atmophyticus*, *Mesostigma viride*, *Klebsormidium nitens*, *Chara braunii*, *Spirogloea muscicola*, *Penium margaritaceum*, and *Mesotaenium endlicherianum* removed.**

The caption stays the same as Figure SI 12.

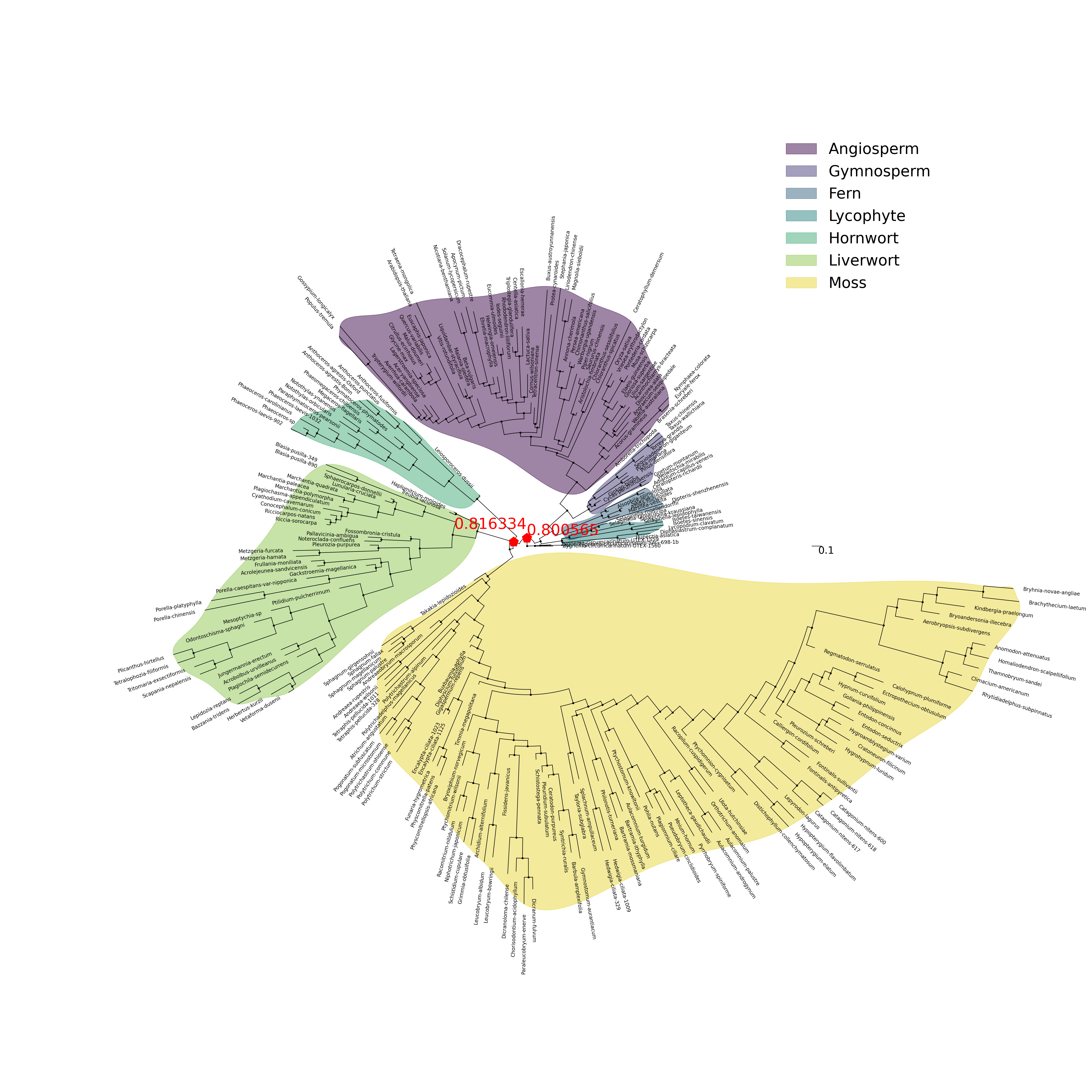

**Figure SI 20. Species tree inferred by the MSC method based on the 137-rSOG dataset with *Chlamydomonas reinhardtii*,** ***Chlorokybus atmophyticus*, *Mesostigma viride*, *Klebsormidium nitens*, *Chara braunii*, *Spirogloea muscicola*, *Penium margaritaceum*, *Mesotaenium endlicherianum*, and *Zygnema cf. cylindricum* removed.**

The caption stays the same as Figure SI 12.

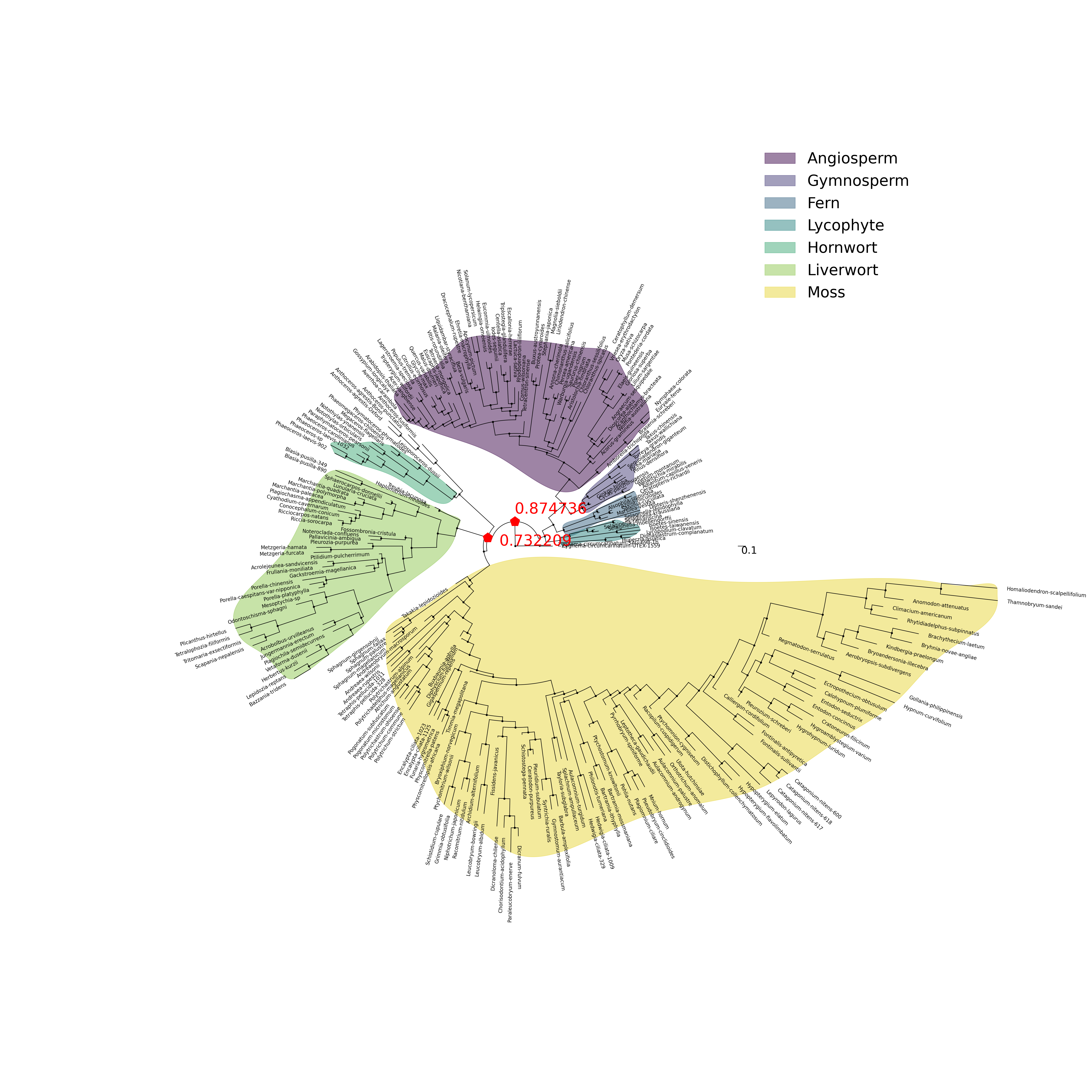

**Figure SI 21. Species tree inferred by the MSC method based on the 137-rSOG dataset with *Chlamydomonas reinhardtii*,** ***Chlorokybus atmophyticus*, *Mesostigma viride*, *Klebsormidium nitens*, *Chara braunii*, *Spirogloea muscicola*, *Penium margaritaceum*, *Mesotaenium endlicherianum*, *Zygnema cf. cylindricum*, and *Z. circumcarinatum UTEX 1560* removed.**

The caption stays the same as Figure SI 12.

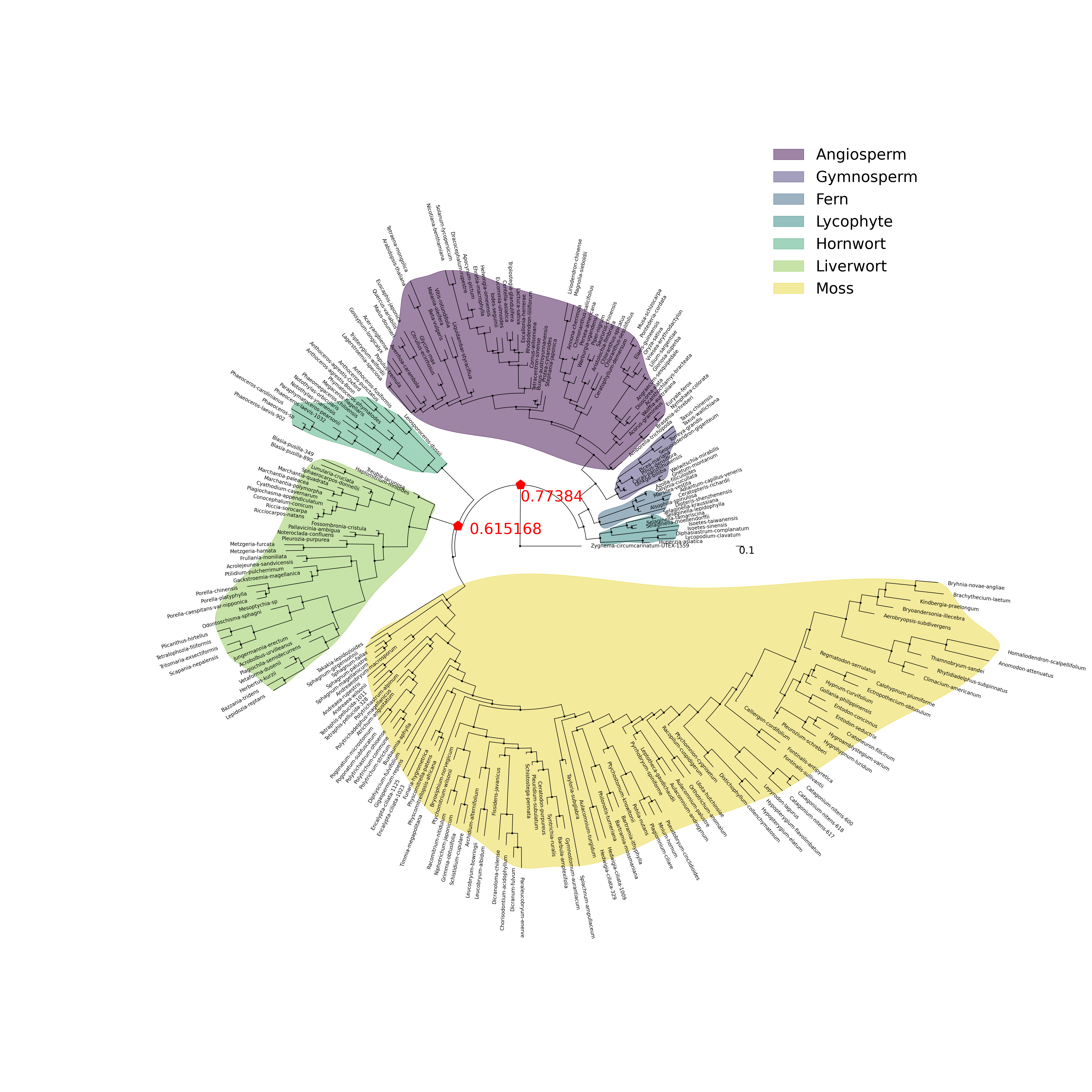

**Figure SI 22. Species tree inferred by the MSC method based on the 137-rSOG dataset with *Chlamydomonas reinhardtii*,** ***Chlorokybus atmophyticus*, *Mesostigma viride*, *Klebsormidium nitens*, *Chara braunii*, *Spirogloea muscicola*, *Penium margaritaceum*, *Mesotaenium endlicherianum*, *Zygnema cf. cylindricum*, *Z. circumcarinatum UTEX 1560*, and *Z. circumcarinatum SAG 698-1b* removed.**

The caption stays the same as Figure SI 12.
